# Fossil-calibrated inference of divergence times among the Volvocine algae enables reconstruction of the steps that led to differentiated multicellularity

**DOI:** 10.1101/2023.09.19.558489

**Authors:** Charles Ross Lindsey, Andrew H. Knoll, Matthew D. Herron, Frank Rosenzweig

## Abstract

Throughout its nearly four-billion-year history, life has undergone evolutionary transitions in which simpler subunits have become integrated to form a more complex whole. Many of these transitions opened the door to innovations that resulted in increased biodiversity and/or organismal efficiency. The evolution of multicellularity from unicellular forms represents one such transition, one that paved the way for cellular differentiation, including differentiation of male and female gametes. A useful model for studying the evolution of multicellularity and cellular differentiation is the volvocine algae, a clade of freshwater green algae whose members range from unicellular to colonial, from undifferentiated to completely differentiated, and whose gamete types can be isogamous, anisogamous, or oogamous. To better understand how multicellularity, differentiation, and gametes evolved in this group, we used comparative genomics and fossil data to establish a geologically calibrated roadmap of when these innovations occurred. Our results, presented as ancestral-state reconstructions, show that multicellularity arose independently twice in this clade. Our chronograms indicate multicellularity evolved during the Carboniferous-Triassic periods in Goniaceae + Volvocaceae, and possibly as early as the Cretaceous in Tetrabaenaceae. Using divergence time estimates we inferred when, and in what order, specific developmental changes occurred that led to differentiated multicellularity and oogamy. We find that in the volvocine algae the temporal sequence of developmental changes leading to differentiated multicellularity is much as proposed by David Kirk, and that multicellularity is correlated with the acquisition of anisogamy and oogamy. Lastly, morphological, molecular, and divergence time data suggest the possibility of cryptic species in Tetrabaenaceae.

## Introduction

Major evolutionary transitions occur when multiple autonomous units (e.g., genes) combine to form an interdependent autonomous unit (e.g., chromosomes) capable of storing and transmitting information in a novel way.^1^ Over the past four billion years a relatively small number of such transitions have resulted in a myriad of innovations that have contributed to the diversification of life on Earth.^1 2^ Among the most conspicuous of these is the transition from organisms whose individuals consist of one cell to individuals that consist of many.

Multicellularity has independently evolved from a unicellular ancestor at least 45 times,^3^ and has been documented in all three domains of life: Bacteria,^4^ Archaea,^5^ and Eukarya.^6^ In most cases, this transition opened the door to evolution of <u>d</u>ivision <u>o</u>f labor (DOL), which in turn paved the way for cellular differentiation, where cells in a multicellular body take on specific tasks. Task specialization has the potential to boost a multicellular organism’s fitness, provided that the programmed partitioning of functions provides metabolic, structural, and/or genetic advantages over retaining all functions in all cells. Cellular differentiation occurs in most but not all multicellular eukaryotes.

Though multicellularity and differentiation have repeatedly evolved, we have limited knowledge of what selects for them and the genetic steps that enable their initial evolution.^7^ To fill this knowledge gap, we can use comparative genomics to study evolutionary lineages, or clades, in which some species are unicellular and others multicellular, and in which the multicellular species exhibit varying degrees of cellular differentiation. Several eukaryotic clades fulfill these requirements^8,9^, notably the volvocine algae^10^, a small extant group of freshwater green algae nested within Viridiplantae in the Chlorophycean order Volvocales (**Figure 1**).

**Figure 1.**
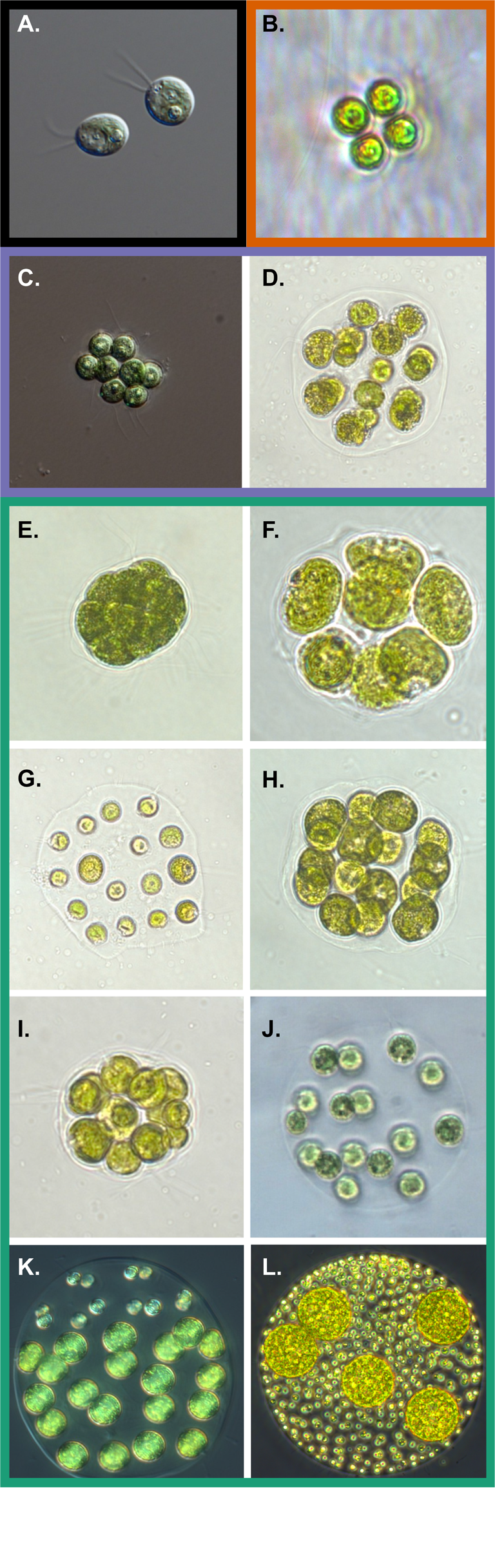
Diversity amongst volvocine genera. (A.) Chlamydomonas reinhardtii, (B.) Tetrabaena socialis, (C.) Gonium pectorale, (D.) Astrephomene gubernaculifera, (E.) Pandorina morum, (F.) Volvulina compacta, (G.) Platydorina caudata, (H.) Colemanosphaera charkowiensis, (I.) Yamagishiella unicocca, (J.) Eudorina elegans, (K.) Pleodorina starrii, (L.) Volvox carteri. Multicellular taxa have orange, purple, and green borders corresponding to Tetrabaenaceae, Goniaceae, and Volvocaceae, respectively. Unicellularity is designated by the black border. Photos are not to scale.

The volvocine algae are facultatively sexual,^11^ thus in addition to being a model system for the study of multicellularity and differentiation, they can also provide insight into another major transition: the evolution of male and female sexes via gametic differentiation. The definition and elaboration of the sexes, as well as the evolution of sexually selected traits, is predicated on the advent of anisogamy, gamete types that differ in size. Oogamy constitutes a specialized form of anisogamy wherein small male gametes are motile and large female gametes are not (e.g., vertebrates and land plants are oogamous). Anisogamy is hypothesized to have evolved from isogamy,^12^ in which the genetic determinants of mating type segregate during meiosis but the resulting gametes are identical in size and shape.^13^ Because multicellular lineages such as animals and plants lack extant isogamous forms, the origins of anisogamy cannot be deduced using these groups. By contrast, the volvocine algae range from unicellular, isogamous species like *Chlamydomonas* to multicellular, oogamous species like *Volvox*, which exhibits complete germ-soma differentiation.

Multicellular volvocine species are grouped within three families: Tetrabaenaceae, Goniaceae, and Volvocaceae. According to recent phylogenetic evidence,^14^ these families are paraphyletic. The unicellular *Vitreochlamys ordinata* is sister to the Tetrabaenaceae. This clade of *V. ordinata* + Tetrabaenaceae is sister to several *Chlamydomonas* species plus the multicellular Goniaceae + Volvocaceae. Both *Vitreochlamys* and *Chlamydomonas* are isogamous, undifferentiated unicells. The genera composing the Tetrabaenaceae, *Tetrabaena* and *Basichlamys*, are 4-celled colonies that are isogamous and undifferentiated. The Goniaceae comprises two genera, *Gonium* and *Astrephomene*. *Gonium* consists of flattened 8-32 celled colonies that are isogamous and undifferentiated. *Astrephomen*e, also isogamous, includes 32-128 celled spheroidal colonies that exhibit somatic cell differentiation. Somatic cells engage in specific cellular functions, principally motility, are mortal, and do *not* pass on genetic information to subsequent generations. Among the 8 genera that comprise the Volvocaceae, *Platydorina*, *Volvulina*, and *Yamagishiella* are isogamous and undifferentiated; *Colemanosphaera*, *Eudorina*, and *Pandorina* are anisogamous and undifferentiated; *Pleodorina* is anisogamous with somatic differentiation, and *Volvox* is oogamous with specialized germ and somatic cells. Specialized germ cells pass on genetic information and are distinct from undifferentiated cells in that they do not significantly contribute to colony motility.

Because the volvocine algae are closely related, but vary in cellularity, differentiation, and sexuality, comparative genomics offers the prospect of understanding how and when each of these traits evolved. Nearly 15 years ago Herron et al.^15^ estimated divergence times for this group while analyzing the temporal sequence of certain developmental traits. Since then, a wealth of molecular, paleontological, and phylogenetic data has come to light, motivating us to reevaluate their conclusions. For example, multiple volvocine genomes^16–20^ have been published, as well as the transcriptomes of all known extant volvocine genera,^14,21^ including a new genus;^22^ furthermore, genomes and transcriptomes have been sampled among different strains within many volvocine species.^14,21^ Also, the fossil record has been enriched by the discovery of a billion-year-old multicellular chlorophyte,^23^ firm dates of more than a billion years have been established for simple, multicellular red algae,^24^ and red algal fossils have been observed that may be 1.6 billion years old.^25^ Using a dataset of 40 nuclear-protein coding genes, Lindsey et al.^14^ recently reported that *(i)* multicellularity evolved at least twice in this group, *(ii)* cell differentiation evolved at least four times, *(iii)* anisogamy evolved from isogamous ancestors at least three times, and *(iv)* oogamy evolved at least three times. The first two of these findings have since been bolstered by Ma et al.^26^

When divergence times were estimated by Herron et al.,^15^ their molecular dataset consisted of 5 chloroplast protein-coding genes plus the 18S nuclear, small ribosomal subunit across 35 volvocine taxa. Subsequent studies undertaking ancestral-state reconstruction^27–30^ of this group have relied solely on the 5 chloroplast gene dataset for their analyses. Unsurprisingly, the resulting trees exhibited essentially the same branching order, none of which reflect the new insights gained by Lindsey et al.^14^ As noted, Ma et al.^26^ also inferred divergence times of the volvocine algae using large nuclear datasets, employing a single relaxed clock model. However, these researchers did not use their phylogenetic inferences to perform ancestral-state reconstruction of sexual and developmental traits.

Here, we provide a geologically-calibrated roadmap of when multicellularity, differentiation, and anisogamy arose in the volvocine algae. Since no reliable fossils exist for this group, we sampled fossils across the Archaeplastida, or kingdom Plantae (*sensu lato*), where reliable fossils are plentiful. Specifically, we sampled 14 fossil taxa across the three major Archaeplastida clades (Rhodophyta, Streptophyta, and Chlorophyta) selecting them so as to calibrate our time-tree over an interval of at least one billion years. Our molecular data consist of amino acid sequences for 263 single-copy nuclear genes drawn from 164 taxa across the Archaeplastida. Sequences were obtained both from publicly available genomic and transcriptomic datasets as well as from our own published RNA-Seq dataset.^14^ Our goal was to reanalyze the divergence times and the gain and/or loss of traits related to multicellularity, differentiation, and sexuality for the volvocine algae. Our data, presented as ancestral-state reconstructions, lead us to conclude that there were two independent origins of multicellularity in the volvocine algae, as predicted by Lindsey et al.^14^ and Ma et al.^26^ Moreover, under one of our four relaxed clock models reassessment of divergence times indicates that multicellularity may have originated in the Goniaceae + Volvocaceae sometime between the Carboniferous and Triassic, much older than previous inferrences.^15,26^ Among the Tetrabaenaceae we find that multicellularity may have arisen as early as the Carboniferous or as recently as the Cretaceous. Using our chronogram, we also reassessed the developmental milestones proposed in David L. Kirk’s 12-Step Program^31^ to differentiated multicellularity in the volvocine algae. Through ancestral state reconstructions, we show that the temporal sequence of events he hypothesized almost 20 years ago is essentially correct. To investigate multicellularity’s role in the evolution of sex in the volvocine algae, we reconstructed 7 sexual traits. When the gain and loss of these traits are mapped onto our time tree, multicellularity’s role as a driver of sexual evolution becomes evident, consistent with the results of Hanschen et al.^30^ Finally, our analyses of morphological, molecular, and divergence time data suggest the possibility of a cryptic species in the multicellular Tetrabaenaceae.

## Results and Discussion

### Selection of orthologues in taxa across the Archaeplastida provides the basis for inferring how multicellularity and cellular differentiation evolved in the volvocine algae

We sampled a total of 164 taxa representing all major clades of Archaeplastida: Rhodophyta, Streptophyta, and Chlorophyta. Specifically, we sampled 12 rhodophytes across 3 red algal subclades, 45 streptophytes representing 13 major green algal and land plant subclades, and 107 chlorophytes, representing prasinophytes, Ulvophyceae, Trebouxiophyceae, and Chlorophyceae, including 68 unicellular and multicellular volvocine algae (**Figure 2**). We chose the red algae for our outgroup, as multiple studies indicate that they are sister to the green algae + land plants (Embryophyta), and that the red and green algae share a common plastid ancestor.^32,33^ Although our chief aim was to discern, on a geological timescale, the evolution of traits leading to multicellularity and differentiation in the volvocine algae, we included other green algal genera such as *Coleochaete* and *Tetraselmis* to further illuminate the history of Viridiplantae. The former is generally considered to be relatively closely related to land plants,^34,35^ while the latter is believed to be sister to the three major Chlorophyta clades.^36^ Because all multicellular lineages necessarily evolved from unicellular ancestors, clarifying which lineages are sister to which multicellular clades will enable a deeper understanding of how this major transition occurred.

**Figure 2.**
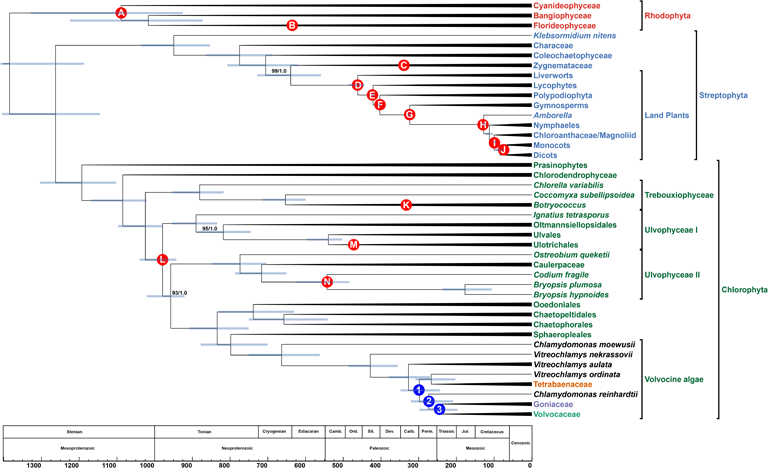
Time-calibrated phylogeny of the Archaeplastida. Branching order of the tree was inferred under maximum-likelihood analysis from an aligned amino acid, concatenated dataset of 263 nuclear genes. Numbers on branches represent bootstrap and posterior probability values, respectively. Branch lengths, corresponding to time, were inferred under the CIR relaxed clock model using 8 most clock-like genes as determined by Sortadate. Blue bars correspond to the inferred 95% HPD interval for each node. Red bubbles correspond to calibrated nodes (Table 1), and blue bubbles correspond to key divergences among the volvocine algae (Figure 3B). Members of the multicellular volvocine algae (Tetrabaenaceae, Goniaceae, and Volvocaceae) are denoted in orange, purple, and green. Taxa in black font are unicellular volvocine algae.

**Table 1.** Fossils used to calibrate nodes in Archaeplastida tree.

| Node | Calibration | Age |
| --- | --- | --- |
| A | *Bangiomorpha* | 1047 +13/-17 |
| B | Oldest Florideophycidae | 609 +/-5 |
| C | Oldest Zgnemataceae | MIN: 350 |
| D | First Land Plants | MIN: 480 |
| E | Oldest Tracheophytes<br>(*Cooksonia* with tracheids) | 423-419 |
| F | Fern-Seed Plant Split | MIN: 385 |
| G | Extant Gymnosperms-Angiosperms Split (based on<br>oldest *Cordaite*s) | 330-323 |
| H | *Amborella* and Nymphaeles<br>Split from other Plants<br>(Barremian flowers and<br>pollen) | 129-125 |
| I | Chloroanthaceae/Magnoliid<br>versus monocots plus<br>eudicot split<br>(Chloranthaceous fossils) | MIN: 125 |
| J | Monocot-Eudicot Split<br>(*Liliacidites* monocot pollen) | MIN: 113 |
| K | *Botryococcus braunii* | 358-356 |
| L | *Proterocladus* | 1056-948 |
| M | Ulotrichales: *Vermiporella* | 470-458 |
| N | *Protocodium* | MIN: 541 |

A single dataset consisting of 263 single-copy, protein coding genes was compiled and analyzed under Maximum-Likelihood (ML), Bayesian inference (BI), and coalescent-based (CB) phylogenetic methods. These 263 genes were conserved across the three main clades of Archaeplastida. Our concatenated alignment for ML and BI analyses represents an aggregate of 79,844 amino acids, equivalent to 239,532 nucleotide positions, with a total of 62,106 parsimony-informative sites. All raw reads used to complete our single-gene and concatenated alignments encompassing 164 total taxa were mined from previously published data located in public repositories (**Supplementary Table S1**).

### Phylogenies of the Archaeplastida inferred by different analytical methods are largely congruent

Maximum-Likelihood (ML), Bayesian inference (BI), and coalescent-based (CB) phylogenies are largely congruent, with nearly all clades well-supported (**Supplementary Figure 1**, **Supplementary Figure 2**, and **Supplementary Figure 3**). The ML and BI phylogenies have identical branching orders (**Supplementary Figure 1** and **Supplementary Figure 2**), and the only differences between the ML + BI and CB trees occur in Chlorophyta, the major green algal and seaweed clade. Within Rhodophyta, our analyses indicate *Galdieria* + *Cyanidiococcus + Cyanidioschyzon* as sister to the Bangiophyceae + Florideophyceae with all branches having high support (ML bootstraps (MLBS) = 100, Bayesian posterior probabilities (BPP) and Coalescent posterior probabilities >= 0.97). Within the Streptophyta, all three analyses inferred an identical branching order, which in “newick” tree format^37^ consists of (Klebsormidiales,(Characeae,(Coleochaetophyceae,(Zygnemataceae, Embryophyta)))).

Regarding the position occupied by the Coleochaetophyceae, our ML + BI and CB trees indicate this clade to be sister to Zygnemataceae + Embryophyta with high support (MLBS = 100, BPP = 1.0, CPP = 1.0). Thus, our results show that the filamentous charophyte Zygnemataceae is the sister group to land plants, consistent with other published works.^38–40^ Among the chlorophyte green algae, our analyses also show that *Tetraselmis* is sister to the three main Chlorophyta families: Trebouxiophyceae, Ulvophyceae, and Chlorophyceae. And lastly, our ML + BI and CB analyses do not accord in the position of the ulvophycean Bryopsidales. All three analyses indicate that the Ulvophyceae are paraphyletic, as noted in recent studies^41,42^ based on dense ulvophyte taxonomic sampling. ML and BI analyses indicate that the Bryopsidales are sister to the Chlorophyceae (MLBS = 93, BPP = 1.0), as previously reported.^41,42^ Our CB tree, however, suggests a branching order of Bryopsidales sister to Ulvophyceae I + Chlorophyceae (CPP = 1.0). Due to these findings, we suggest that the Ulvophyceae are indeed paraphyletic, with the Bryopsidales likely a sister group to the Chlorophyceaen green algae.

### Phylogenetic analyses of the volvocine algae indicate multiple independent origins of multicellularity and differentiation

Our ML, BI, and CB analyses all indicate two independent origins of multicellularity among the volvocine algae: one in the lineage leading to the Tetrabaenaceae and another in the lineage leading to the Goniaceae + Volvocaceae (**Figure 2**, **Supplementary Figure 1**, **Supplementary Figure 2**, and **Supplementary Figure 3**). These results bolster the findings of Lindsey et al.^14^ and Ma et al.^26^ As before, our ML and BI analyses have identical branching orders. The Tetrabaenaceae + *Vitreochlamys ordinata* are shown to be sister to several *Chlamydomonas* species + the Goniaceae and Volvocaceae in our ML and BI trees (MLBS = 100, BPP = 1.0) (**Figure 2**, **Supplementary Figure 1**, and **Supplementary Figure 2**), and this topology both corroborates and expands on with those presented in several earlier studies.^14,26^ Our CB tree, however, indicates a slightly different branching order for the Tetrabaenaceae. The resulting coalescent-based tree shows the Tetrabaenaceae + *V. ordinata* forming a clade with *Chlamydomonas reinhardtii* and its relatives, and this clade is shown to be sister to the Goniaceae + Volvocaceae (CPP = 1.0) (**Supplementary Figure 3**). Of note, the sister relationship between the Tetrabaenaceae + *V. ordinata* and the *Chlamydomonas* clade is poorly supported in our CB analysis (CPP = 0.34).

In accordance with the findings of Lindsey et al.,^14^ all three of our phylogenetic analyses indicate a minimum of 4 independent origins of somatic cellular differentiation and a minimum of 3 origins of anisogamy among the volvocine algae. Origins of somatic cell differentiation occur in the following lineages: (*i*) *Astrephomene*, (*ii*) section *Volvox*, (*iii*) *Pleodorina thompsonii*, and (*iv*) the *Pleodorina japonica* + *Volvox carteri* clade. Anisogamy evolved from isogamous ancestors at least three times in the following lineages: (*i*) *Astrephomene,* (*ii*) section *Volvox*, and (*iii*) in the *Eudorina* + *Volvox* + *Pleodorina* (EVP) clade.

In contrast to several recent volvocine studies,^14,20^ all three of our phylogenetic analyses support the major conclusion that the Goniaceae are monophyletic, albeit with varying support values (MLBS = 95, BPP = 1.0, CPP = 0.67). This conclusion was reached by multiple earlier investigations.^15,26,29,30,43–50^ Consistent with results of Lindsey et al.^14^ and Ma et al.,^26^ we find that *Volvox* section *Volvox* is sister to the *Pandorina* + *Volvulina* + *Colemanosphaera* (PVC) and the (EVP) clades within the Volvocaceae. Our ML + BI and CB trees contain minor branching order differences in the EVP clade, but these have no bearing on our major findings.

### Variation in divergence times inferred under different relaxed molecular clock models necessitate validation tests

All divergence time estimates were inferred by Phylobayes 4.1b^51^ under a Bayesian approach for our inferred ML and CB species trees. For each topology, a total of 14 nodes were calibrated across Rhodophyta, Streptophyta, and Chlorophyta (**Figure 2** and **Table 1**), and four relaxed clock models were tested during estimation of divergence times: the autocorrelated lognormal^52^ (LN) and CIR^53^ models, and the uncorrelated gamma (UGAM) and white-noise (WN) models.^54^ Inferred divergence times established under all clock models are largely consistent across the three major red and green algal clades (**Figure 3**). However, there are nodes such as the root age, earliest rhodophyte divergence, and major divergences within the volvocine algae where one model infers a date markedly younger or older than the others (**Figure 3**).

**Figure 3.**
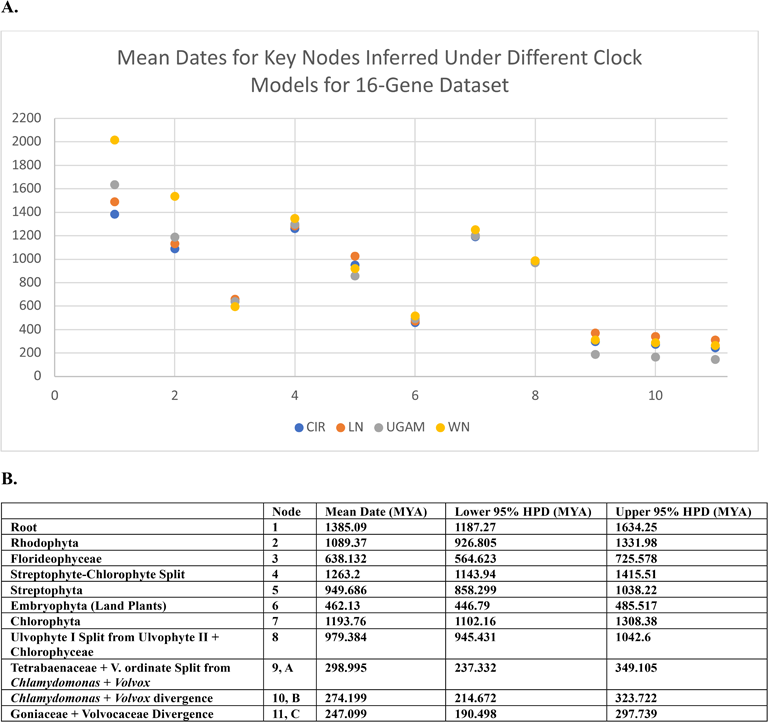
Scatter plot of mean dates for key nodes across Archaeplastida inferred under different clock models. The y-axis represents time in MYA (millions-of-years ago), and the x-axis corresponds to specific nodes labeled in table. The four clock models represented in each key node include: LN (blue), CIR (orange), WN (gray), and UGAM (yellow).

The estimated mean root age of all tested clock models varies by ∼630 million years (MY) with the WN model estimating the earliest red algal divergence as ∼2016 MYA, and the CIR model estimating it as late as ∼1385 MYA (**Figure 2B**). Averaging the mean root ages of the LN, CIR, and UGAM models produces an average root age of ∼1503 MYA, reducing the variance in estimated mean root age between WN and the others to ∼500 MY. Similarly, for the earliest rhodophyte divergence, the WN model inferred a mean date significantly older than all other models by at least 300 MY, whereas the LN, CIR, and UGAM models estimated mean ages within <100 MY of each other. Large date discrepancies such as these may be solely due to differences in clock algorithms.^55,56^

For all volvocine algae divergences, the UGAM model estimated ages markedly younger than the three other models we tested (**Figure 3A, B** (divergences 9-11)). It is noteworthy that the inferred ages of the UGAM model for the volvocine algae are very similar to the dates inferred by Ma et al.,^26^ who used a single relaxed clock model. When taking the average of the mean dates inferred by the LN, CIR, and WN models, there is a consistent ∼130 MY difference between the average and a mean node date inferred by the UGAM model for this group. Significantly, the UGAM 95% highest posterior density (HPD) intervals for the volvocine algal divergence do not overlap with any of the 95% HPD intervals inferred by other models, conversely the other three models see a comfortable overlap in their 95% HPD intervals for volvocine divergence estimates.

Given that the WN and UGAM models inferred outlier dates for certain key divergence events, we decided to exclude both as clock models. Ages inferred by the CIR model in this study were never observed to be outliers for key divergence events, and the ages inferred by the CIR model for the volvocine algae are nearly identical to the WN model’s estimates for this group. Additionally, divergence estimates inferred under the CIR model for the volvocine algae were the most conservative among the LN, WN, and CIR results. Altogether, the foregoing considerations prompted us to report dates for major divergence events using data produced by the CIR relaxed clock model (**Figure 3A, B**). Fossil cross validation tests were performed as described earlier for each relaxed clock model, with the objective of identifying a best-performing analysis. However, no relaxed clock model performed markedly better than any of the others (**Supplementary Table 2**).

### Fossil selections and Archaeplastida divergence times

Since no reliable fossils exist for the volvocine algae, we selected 14 fossil taxa across the Archaeplastida where reliable fossils are abundant (**Figure 2** and **Table 1**). For each primary fossil calibration used in this study, each calibration point was constrained to a range rather than a fixed-point estimate, acknowledging the inherent uncertainty in fossil ages. With each date range, we specified a soft bound where 2.5% of the total probability mass is positioned outside of the specified lower and/or upper bounds. Detailed information regarding each fossil taxon and calibration point may be found in **Supplementary Methods.** Moreover, our results indicate that divergence times among the Archaeplastida inferred under the CIR model are largely congruent with those of prior studies. With the volvocine algae being our principal clade of interest, we elected to report general Archaeplastida divergence times in **Supplementary Methods.**

### Molecular clock analysis reveals that the crown Tetrabaenaceae-Goniaceae-Volvocaceae clade arose sometime between the Carboniferous and Triassic, and ancestral state reconstructions confirm our prior inference that multicellularity arose twice

According to our <u>a</u>ncestral-<u>s</u>tate <u>r</u>econstructions (ASR) (**Supplementary Figures 4** and **5**), multicellular lineages share a common unicellular ancestor ∼298 MYA (CIR relaxed clock model, 95% HPD interval 349-237 MYA), marking the emergence of the crown <u>T</u>etrabaenanceae-<u>G</u>oniaceae-<u>V</u>olvocaceae (TGV) clade sometime in the Carboniferous to the Triassic periods (**Figure 3B** and **Figure 4, node A**). Our 95% HPD interval for the TGV clade partly overlaps with ranges reported in previous studies.^15,26^ For example, the range reported by Herron et al.^15^ is mostly confined to the Triassic (260-209 MYA), whereas that reported by Ma et al.^26^ extends across the Early Triassic and Early Jurassic (251-180 MYA).

**Figure 4.**
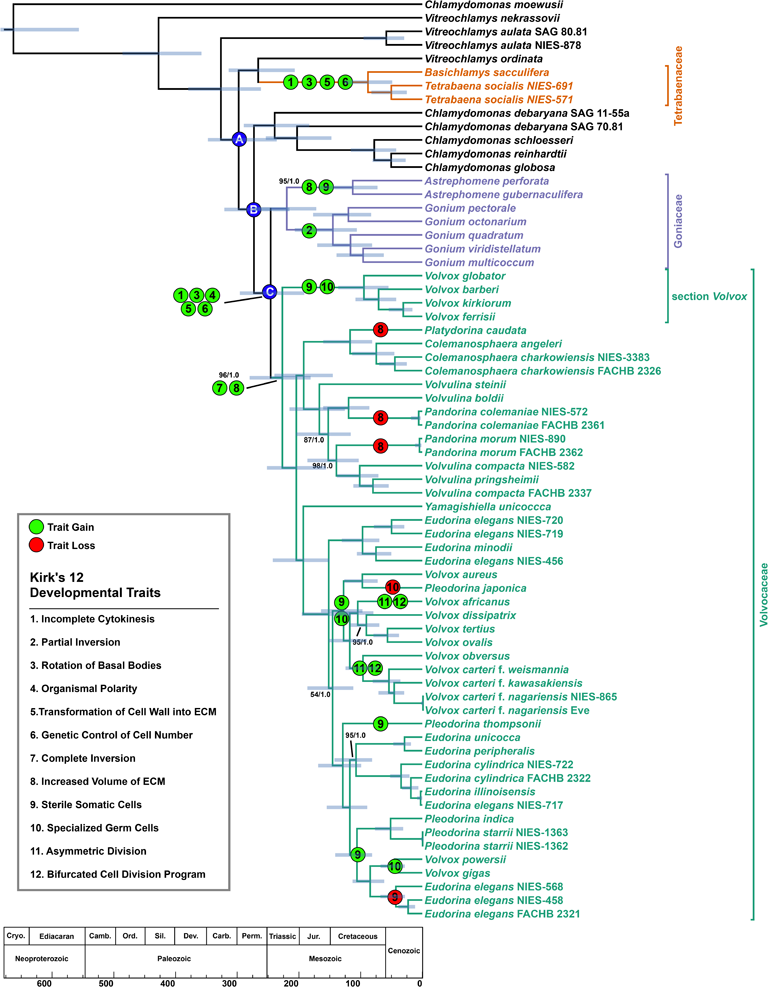
Estimated divergence times of the volvocine algae. Branching order of the tree was inferred under maximum-likelihood analysis from an aligned amino acid, concatenated dataset of 263 nuclear genes. Numbers on branches represent bootstrap and posterior probability values, respectively. Branch lengths, corresponding to time, were inferred under the CIR relaxed clock model using 8 most clock-like genes as determined by Sortadate. Blue bars correspond to the inferred 95% HPD interval for each node. Green bubbles correspond to a developmental trait gain, and red bubbles corresponds to loss of a trait. The figure table lists the 12 developmental traits identified by Kirk in their original order. Blue bubbles indicate key divergences in the volvocine algae. Members of the multicellular volvocine algae (Tetrabaenaceae, Goniaceae, and Volvocaceae) are denoted in orange, purple, and green. Taxa in black font are unicellular volvocine algae.

From the TGV clade’s common ancestor (**Figure 4, node A**), our ASR results show that two unicellular lineages diverged (**Supplementary Figures 4** and **5**), each of which led to independent origins of multicellularity in the volvocine algae (**Figure 2**, **Figure 3**, and **Figure 4**). From one branch of this split, a multicellular ancestor of the Tetrabaenaceae diverged from a unicellular ancestor of *Vitreochlamys* ∼267 MYA (316-206 MYA), and from the other branch a multicellular ancestor of the Goniaceae + Volvocaceae diverged from a unicellular ancestor of *Chlamydomonas* ∼274 MYA (323-214 MYA (**Figure 3B** and **Figure 4**). These results confirm two independent origins of multicellularity among the volvocine algae as indicated by previous phylogenies.^14,26,45^ According to our data, multicellularity arose in Tetrabaenaceae sometime between the Jurassic and Cretaceous (126-49 MYA), while multicellularity arose in the Goniaceae + Volvocaceae during the Permian-Jurassic (297-190 MYA).

### Ancestral state reconstructions suggest that the steps leading to differentiated multicellularity in the volvocine algae occur in the order envisioned by Kirk

In 2005, David L. Kirk, a developmental and cell biologist, hypothesized how differentiated multicellularity, as manifest in *Volvox carteri*, might have evolved from a unicellular, *Chlamydomonas*-like ancestor.^31^ In his essay, attention was brought to 12 ontogenetic traits (**Figure 4)** that occur in the developmental cycle of *V. carteri* as it matures from a single-cell to a >500-celled organism. These 12 traits were then shown to be present in various extant Goniaceae and Volvocaceae, allowing them to be positioned in a stepwise manner along a simplified volvocine phylogeny, beginning with the most recent common ancestor of *C. reinhardtii* and *V. carteri*. Through the ordered mapping of these traits, Kirk suggested a developmental program that could explain how multicellularity with a division of labor might evolve. This hypothesis, known as Kirk’s 12-Step Program to differentiated multicellularity, has been formally tested once by Herron et al.^15^ using ancestral-state reconstructions from a previous study.^43^ Herron et al.^15^ suggested that the 12-Step Program was not as orderly as hypothesized due to the phylogenetic position of the Tetrabaenaceae, which are absent from Kirk’s hypothesis. Below, we show that, using ancestral-state reconstructions based on our more robust phylogeny, Kirk’s 12-Step Program occurs temporally much as originally hypothesized.

For continuity, Kirk’s original order of 12 developmental traits has been retained in our analysis (**Figure 4**). Approximately 298 MYA (349-237 MYA), two lineages emerged from the common unicellular ancestor of the colonial volvocine algae (**Figure 4**, **node A**). One of these lineages eventually gave rise to the Tetrabaenaceae (**Figure 4**). In this lineage, *Steps 1, 3, 5* and *6* evolved in the multicellular ancestor of *Basichlamys* and *Tetrabaena* (**Figure 4** and **Supplementary Figures 6-13**). According to our ASR, this multicellular ancestor was established ∼89 MYA (126-49 MYA); no other traits identified by Kirk are present either in this ancestor or in any of the known extant Tetrabaenaceae. Importantly, the number of Kirk’s traits gained in the lineage leading to the Tetrabaenaceae is twice that reported by Herron et al.^15^ This conclusion is supported by recent studies confirming incomplete cytokinesis (*Step 1*) and basal body rotation (*Step 3*) in *Tetrabaena*.^57,58^

The second lineage diverged from a *Chlamydomonas*-like ancestor ∼274 MYA (323-214 MYA), and eventually gave rise to the multicellular Goniaceae and Volvocaceae (**Figure 4**, **node B**). ASR indicates that the common ancestor to the Goniaceae and Volvocaceae was multicellular (**Supplementary Figures 4** and **5**), that it had evolved 5 of the 12-Step Program traits over a period of ∼27 MY (**Figure 4, node C**) (**Supplementary Figures 6-15**), and was most likely non-spheroidal (**Supplementary Figures 16 and 17**). By ∼247 MYA (297-190 MYA), the non-spheroidal, multicellular ancestors to the Goniaceae and Volvocaceae had evolved: (*Step 1*) an incomplete cytokinesis where daughter cells were left connected to each other via cytoplasmic bridges, (*Step 3*) peripheral colony cells with rotated flagellar basal bodies beating in parallel allowing the colony to efficiently mobilize through water, (*Step 4*) central-to-peripheral polarity similar to what is seen in *Gonium*, (*Step 5*) daughter cells embedded into an extracellular matrix with two distinct boundaries, and (*Step 6*) colony cell number controlled by its genetic code rather than the environment (**Figure 4**).

Due to the way that volvocine cells divide, embryos resulting in *Gonium* and Volvocaceae colonies must undergo a process known as “inversion.” Following cell division in embryogenesis, volvocine embryos (whether *Gonium* or volvocacean) contain all the cells that will be present in the mature adult colony. At this stage, a developmental hurdle must be overcome for the eventual colony to swim. For *Gonium*, embryos are in the shape of a bowl with their flagellar ends positioned in the concave region, and volvocacean embryos, in similar fashion, have their flagellar ends located inside the developing colony. By ∼146 MYA (206-106 MYA), the ancestor to *Gonium* had developed (*Step 2*) “partial” inversion whereby a curvature reversal in the embryo occurs, allowing the flagellar ends to be situated on the exterior of the colony. By ∼228 MYA (279-175 MYA), volvocacean embryos developed “complete” inversion (*Step 7*), where cells, in a coordinated fashion, move and curl outward along the anteroposterior axis, resulting in an embryo whose flagellar ends point outside the colony (**Supplementary Figure 18** and **19**) (*Step7*). In both examples, cells in colonies are held in position by cytoplasmic bridges formed via incomplete cytokinesis (*Step 1*). Along with *Step 7*, by ∼228 MYA (279-175 MYA) spheroidal volvocacean ancestors had evolved body plans with increased volume. Diminished colony volume occurred secondarily in lineages leading to *Platydorina* and the paraphyletic *Pandorina* (**Figure 4**).

The spheroidal body plan in the Volvocaceae and in the lineage leading to *Astrephomene* likely consisted of cells positioned along a colony’s periphery, resulting in a transparent organism with an interior filled with extracellular matrix. Despite the morphological similarity among spheroidal colonies, earlier studies^59,60^ and our ASR results indicate that such a body plan arose independently twice in the volvocine algae (**Supplementary Figures 16** and **17**). Yamashita et al.^59^ elucidated the developmental mechanisms that underlie the two independently evolved forms. During *Astrephomene* embryogenesis daughter protoplasts gradually rotate their apical ends so that flagella extend towards the posterior of the growing colony after each successive cell division. Development proceeds differently in the Volvocaceae. In volvocacean early embryos, chloroplasts are positioned at one end of each protoplast, facing outward. As the embryo matures, these cells change shape, with the chloroplast ends forming acute edges.^60^ This cell shape change results in bending the epithelium at an opening of the embryo known as the phialopore.^61^ Epithelial bending causes the colony to turn itself inside-out allowing flagella to be located on the exterior of the developing colony. This process in the Volvocaceae is known as “complete” inversion.^59^ Yamashita and Nozaki^60^ showed that neither of these developmental mechanisms occur in *Tetrabaena* or *Gonium* embryogenesis and thus concluded that spheroidal body plans in the volvocine algae are independently derived via separate mechanisms.

In our updated phylogeny, the most recent common ancestor of the Volvocaceae exhibited 7 of the 12 traits identified by Kirk (**Figure 4**). Moreover, there is only a single conflict in the order in which they evolved according to his hypothesis. This conflict arises from ambiguity as to when inversion first evolved in a volvocine ancestor; indeed, ASR yields alternative inferences depending upon which analytical assumptions are used. When we consider this conflict as an unordered 3-state problem (no inversion, “partial” inversion, or “complete” inversion) (**Supplementary Figure 20**), which assumes that no state is a prerequisite for another to evolve, the ancestor to the Goniaceae and Volvocaceae is shown not to have evolved inversion (**Figure 4**, **node C**). However, when the analysis is performed as an “ordered” 3-state problem, where “partial” inversion is required for “complete” inversion to evolve, the common ancestor to the Goniaceae and Volvocaceae is shown to have partial inversion (**Supplementary Figure 21**). Lastly, when inversion is treated as a 2-state problem (e.g., no inversion, inversion), the common ancestor of the two clades is shown to have evolved inversion (**Supplementary Figure 22**). This last result conflicts with Yamashita and Nozaki’s^60^ explanation for how inversion in *Gonium* and the Volvocaceae are mechanistically distinct. Inversion should be treated as different states (e.g., “partial” and “complete”), as in our first example of an “unordered” 3-state problem, because it is unknown whether “partial” inversion is necessary for “complete” inversion to evolve. Thus, we report that only the ancestor to *Gonium* had evolved “partial” inversion (*Step 2*) by ∼146 MYA (206-106 MYA).

The evolution of somatic cellular differentiation (*Step 9*) is the first step towards a complete reproductive division of labor where germ cells do *not* participate in motility. Somatic cells are distinct from two other cell types found in the volvocine algae: undifferentiated cells and germ cells. True somatic cells are distinct in that they assume specific cellular functions, notably motility, exhibit a finite lifespan, and do *not* pass on genetic information to subsequent generations. Undifferentiated cells participate in motility and can also undergo mitotic division to produce the next generation. By contrast, fully-differentiated germ cells also undergo mitotic division to generate progeny, but these cells never significantly participate in motility. Our ASR results compared to our phylogenetic results allow for a specific number of independent origins of sterile somatic cells to be stated. Thus, the gain of sterile somatic cells (*Step 9*) occurred five times in three different genera of the volvocine algae (**Figure 4**). According to our results, true somatic differentiation arose in (*i*) *Pleodorina thompsonii* sometime after it diverged from the common ancestor it shared with *V. carteri* (169-99 MYA) and in the common ancestors of: (*ii*) *Astrephomene* (170-75 MYA), (*iii*) section *Volvox* (136-57 MYA), (*iv*) the *Volvox* + *Pleodorina japonica* clade (166-98 MYA), and (*v*) *P. starrii* + *V. gigas* (140-82 MYA) (**Figure 4** and **Supplementary Figures 23** and **24**). In addition to this trait being gained, it was also lost once in the *Eudorina* clade sister to *V. gigas* + *V. powersii*.

The advent of true germ-soma differentiation in the volvocine algae is characterized by evolution of specialized germ cells (*Step 10*) that do not significantly contribute to colony motility. In genera like *Astrephomene* and *Pleodorina* that have not made this step, colonies consist of somatic cells as well as biflagellate gonidia, cells that contribute to motility and can undergo mitotic division. Indeed, in these taxa gonidia greatly outnumber somatic cells. In *A. gubernaculifera* a 32 or 64-celled colony may have 30 or 60 gonidia, respectively,^62^ and 128-celled *P. californica* colonies can have >70 gonidia.^63^ Gonidia in these instances contribute significantly to motility until cell division is initiated, which ultimately leads to hatched daughter colonies and subsequent death of the mother. In contrast, *Volvox* colonies are all composed of >500 somatic cells and far fewer flagellated or aflagellate gonidia. For example, at the end of *V. rousseletii*’s embryogenesis, a colony may consist of ∼2000-14,000 biflagellate, nearly indistinguishable cells with only 1-16 gonidia.^64,65^ In *V. rousseletii*, gonidia contribute to motility for <20% of a colony’s asexual life cycle before resorbing their flagella and undergoing mitotic division.^65^ *Volvox carteri* colonies are typically composed of ∼2000-6000 cells with ∼8 gonidia.^64^ Somatic and gonidial cells are clearly defined during *V. carteri*’*s* embryogenesis. Gonidial cells are aflagellate; thus, in this species gonidia never contribute to colony motility during its life cycle.^65^ Therefore, *Volvox* spp. are the only extant volvocine algae to have evolved specialized germ cells (*Step 10*). In this group, specialized germ cells evolved by ∼96 MYA (136-57 MYA) in section *Volvox*, ∼46 MYA (68-28 MYA) in the ancestor to *V. gigas* and *V. powersii,* and ∼129 MYA (166-98 MYA) in *V. aureus* and *V. carteri’*s most recent common ancestor (**Figure 4** and **Supplementary Figures 23** and **24**). This trait was lost once in *P. japonica* (**Figure 4** and **Supplementary Figures 23** and **24**).

The final two steps of the 12-Step Program are coupled, consisting of asymmetric cell division (*Step 11*) and a bifurcated cell division program (*Step 12*) (**Supplementary Figures 25-28**). Both traits evolved twice in the genus *Volvox*: (*i*) *V. africanus* sometime after ∼106 MYA and (*ii*) ∼98 MYA for the ancestor of *V. obversus* and *V. carteri*. During *V. carteri* embryogenesis, anterior embryo cells undergo asymmetric divisions producing both small and large cells. Size controls cell fate during embryogenesis. Cells <8μm may divide up to 6 times before they arrest, whereas larger cells undergo 1-2 asymmetric cell divisions. Smaller cells, and the remaining embryonic cells, eventually become soma, whereas the larger cells are destined to become specialized germ cells. Through these two coupled processes (*Steps 11* and *12*), complete division of labor is achieved between somatic and germ cell lines.

Ancestral state reconstruction shows that evolution did not progress linearly within the volvocine algae from a unicellular organism like *Chlamydomonas* to one that is multicellular and differentiated like *V. carteri.* Numerous trait gains and losses occurred in the history of this clade, exemplifying the non-linear nature of evolution. However, Kirk’s 12-Step Program does not attempt to linearize the evolutionary history of this group, but rather to provide a roadmap of steps needed for true differentiated multicellularity to evolve in the volvocine algae. Our results indicate that the temporal sequence in which Kirk organized his 12 traits aligns well with the inferred evolutionary history of this group.

### Ancestral-state reconstructions support previous findings that multicellularity drives evolution of sexual traits in the volvocine algae

The volvocine algae in their various forms have given rise to multicellular lineages that express a variety of sexual traits. For this group, asexual reproduction through mitosis is the typical method that gives rise to the next generation of unicells or colonies. Sexual reproduction, however, is facultative, and is precipitated by reduction in environmental nitrogen and in some cases a sex inducing hormone.^66^ In either case, a diploid zygote is formed after fertilization resulting in a spore. Meiosis only occurs in the volvocine algae when the spore germinates to produce haploid progeny.^66^

Unicellular forms, such as *Chlamydomonas* (*sensu* Pröschold et al.^67^), and in some multicellular forms (*Astrephomene*, *Basichlamys*, *Gonium*, *Pandorina*, *Tetrabaena*, *Volvulina*, and *Yamagishiella*), *plus* and *minus* type gametes are morphologically identical (isogamy) However, in many multicellular forms (*Colemanosphera*, *Eudorina*, *Platydorina*, and *Pleodorina*) the two gamete types differ in size, with the smaller of the two recognized as being male (anisogamy).^68^ In certain instances (*Volvox*), gametic differentiation may be further exaggerated, with the small male gamete being motile, and the large female gamete being sessile (oogamy). One longstanding question is: why does anisogamy evolve in the first place? Another is: what role does multicellularity play in gametic differentiation? Because the volvocine algae encompass both unicellular and multicellular forms, and because they exhibit the full range of known gamete types, they are especially well-suited to addressing these two fundamental questions.

Hanschen et al.^30^ systematically investigated whether multicellularity is a prerequisite for the evolution of anisogamy and whether multicellular volvocines exhibit greater complexity in the form of more elaborate sexual traits. Their study, like ours, concludes that multicellularity is correlated with the acquisition of anisogamy, as all anisogamous volvocines are multicellular, and anisogamy appears to evolve from isogamous ancestors (**Figure 5**).^30^ Using our updated phylogeny, with its newly inferred relationships, we conducted ASR for 7 sexual traits identified in previous studies.^29,30^ Except where noted (**Figure 5**), we focused on the same strains used in the two prior studies of how sexual traits are distributed among the volvocine algae. Like those earlier studies, we excluded *Vitreochlamys*, *Pleodorina thompsonii*, and *Volvox ovalis* from certain ASR analyses due to a lack of information about how they reproduce sexually.

**Figure 5.**
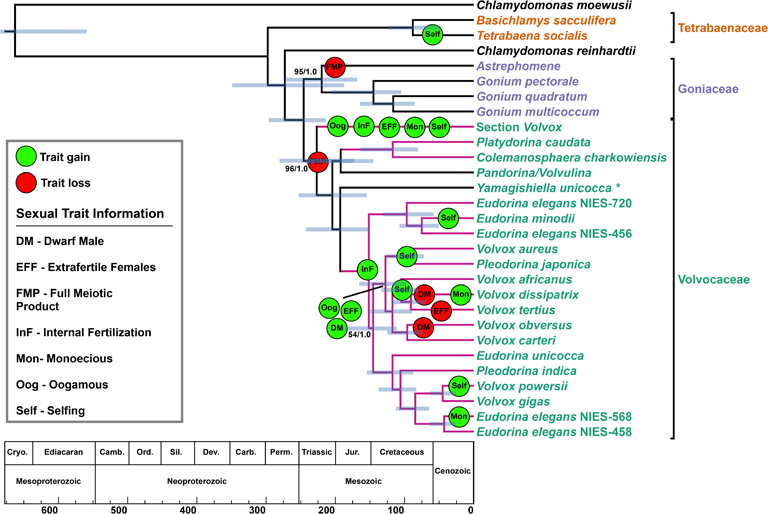
Sexual trait gain and loss in the volvocine algae. Branching order is of a collapsed tree inferred under maximum-likelihood analysis from an aligned amino acid, concatenated dataset of 263 nuclear genes. Numbers on branches represent bootstrap and posterior probability values, respectively. Branch lengths, corresponding to time, were inferred under the CIR relaxed clock model using 8 most clock-like genes as determined by Sortadate. Blue bars correspond to the inferred 95% HPD interval for each node. Green bubbles correspond to a sexual trait gain, and red bubbles corresponds to loss of a trait. The figure table lists the 7 mapped sexual traits and their acronyms. Members of the multicellular volvocine algae (Tetrabaenaceae, Goniaceae, and Volvocaceae) are denoted in orange, purple, and green. Taxa in black font are unicellular volvocine algae.

Unicellular ancestors of the volvocine algae, like *Chlamydomonas*, express the following sexual phenotypes: *(i)* isogamy, *(ii)* full meiotic hatching of zygospores, *(iii)* dioecy, *(iv)* outcrossing, *(v)* external fertilization of the egg, and no evidence of sexual dimorphisms such as (*vi*) “extrafertile” females or (*vii*) “dwarf” males. Prior to the evolution of anisogamy, the number of “gone cells” released from a zygospore decreases. (Note: a gone cell constitutes the haploid meiotic product arising from a diploid zygospore.^69^) *Chlamydomonas reinhardtii*, *Tetrabaena*, and *Gonium* produce all four haploid gone cells per diploid zygospore.^47^ According to our ASR results full meiotic hatching is lost either once or twice in the volvocine algae (**Supplementary Figures 29** and **30**) Our Mr. Bayes ASR indicates a single loss, but this could be an artifact arising from low sampling of taxa that exhibit full meiotic hatching. In contrast, and consistent with Hanschen et al.,^30^ our Phytools ASR indicates two losses of full meiotic hatching in the following lineages: once in *Astrephomene* sometime after diverging from *Gonium* ∼220 MYA (274-170 MYA), and again in the Volvocaceae ∼228 MYA (279-175 MYA) (**Figure 5** and **Supplementary Figures 29** and **30**). Each loss represents a reduction in meiotic products from four haploid gone cells to a single haploid gone cell and three polar bodies. Among our identified sexual traits, no other gain or loss occurs until the evolution of anisogamy.

The most significant difference between our findings and those of Hanschen et al.^30^ is the number of times we find anisogamy to have evolved independently. Their results inferred two independent origins of anisogamy from isogamous ancestors, whereas we conclude three, consistent with Lindsey et al.^14^ (**Figure 5** and **Supplementary Figures 31** and **32**).^14,30^ Section *Volvox*’s position as sister to all other Volvocaceans underlies this distinction. The three origins of anisogamy in our analysis occur in the ancestors of the following lineages: (*i*) section *Volvox* (136-57 MYA), (*ii*) *Platydorina* + *Colemanosphaera* (113-46 MYA) and (*iii*) the *Eudorina* + *Volvox* + *Pleodorina* (*EVP*) clade (196-117 MYA) (**Figure 5** and **Supplementary Figure 31** and **32**). It is noteworthy that we infer three independent origins of anisogamy, regardless of whether this trait is treated as a 2-state problem (e.g., isogamy and anisogamy) or a 3-state problem (e.g., isogamy, anisogamy, or oogamy) (**Supplementary Figures 31-34**).

Consistent with our mapping of developmental traits, anisogamy evolved in lineages that had already acquired a diverse suite of characters differing from a unicellular ancestor. In each case, the eventual anisogamous ancestor had evolved a spheroidal body plan with increased colony size, polarity, and in the case of section *Volvox*, may also have acquired true germ-soma differentiation. It is possible that in some anisogamous ancestors, the body plan was larger with distinct cell types compared to a *Pandorina*-like or *Volvulina*-like morphology, as hypothesized by Hanschen et al.^30^ In section *Volvox* and the *EVP* clade, internal fertilization was gained around the same time (sometime after ∼228 MYA (279-175 MYA) for section *Volvox* and ∼152 MYA (196-117 MYA) for the *EVP* clade) as anisogamy, suggesting that anisogamy might be a prerequisite for this trait (**Figure 5** and **Supplementary Figures 35** and **36**).

After gaining anisogamy and internal fertilization, oogamous lineages appear rapidly to gain several other sexual traits (**Figure 5**). This suggests that oogamy, an extreme form of anisogamy, may drive further sexual evolution. By ∼95 MYA (136-57 MYA), section *Volvox*’s ancestor had become monoecious (i.e., colonies could produce both sperm and ova) and capable of selfing. Necessarily, such an ancestor had lost dioecy (i.e., colonies able to produce only sperm or ova) as well as the capacity to outcross (**Supplementary Figures 37-40**). By the same time, this ancestor gained a sexual dimorphism in the form of “extrafertile” females (**Figure 5**). During sexual reproduction, an “extrafertile” female *Volvox* colony exhibits a doubling in the number of eggs inside its colony, thus giving sexual reproduction the potential to numerically outperform an asexual reproductive cycle.

Mr. Bayes and PhyTools ASR results indicate different ancestral states for when the “dwarf” male phenotype was gained. Results from Mr. Bayes ASR (MBASR) indicate “dwarf” males were gained independently in the lineages leading to *V. africanus*, *V. tertius*, and in the common ancestor to *V. carteri* f. *nagariensis* and *V. carteri* f. weismannia (**Supplementary Figure 43**). Phytools ASR, however, indicate that the common ancestor to *V. carteri* and *V. africanus* had evolved the “dwarf” male phenotype with two losses occurring in the lineages leading to *V. obversus* and *V. dissipatrix* (**Supplementary Figure 44**). These differences arise due to model selection. MBASR results are replicated in Phytools if the “equal rates” model is exclusively used (**Supplementary Figure 45**). When an ANOVA test is performed in Phytools comparing the “equal rates” and “all-rates-different” models, the “all-rates-different” model has a markedly higher weight (0.70 vs 0.29) indicating that it is highly favored over the “equal rates” model. Therefore, in the common ancestor to *V. carteri* and *V. africanus*, two sexual dimorphisms, “extrafertile” females and “dwarf” males, were gained by ∼118 MYA (152-90 MYA) (**Figure 5** and **Supplementary Figures 41-42** and **44**). The “dwarf” male phenotype is expressed as an exaggeratedly reduced male colony size when compared to female colonies.

Within this same clade, *V. africanus* and *V. dissipatrix* independently evolved monoecious, selfing colonies, in addition to the previously mentioned sexual dimorphisms. Sexual trait gains occur in various other lineages of the volvocine algae, but none are as clear as the ones occurring in the two oogamous lineages of *Volvox*.

In summary, we conclude that multicellularity drives the evolution of anisogamy. Once anisogamy is established, a variety of other sexual traits may arise, notably the evolution of oogamy, which in turn opens the door for more elaborate forms of sexual dimorphism such as extrafertile females and dwarf males. Thus, our more robust phylogeny of the volvocine algae, based on not 5 but on 263 protein coding loci, confirms previous findings^29,30^ that multicellularity ultimately drives the evolution and elaboration of diverse sexual traits.

### Molecular clock analysis also suggests cryptic volvocine species in the Tetrabaenaceae

Much like the longstanding debate on what constitutes a biological species,^70,71^ consensus is lacking for how we should recognize cryptic species.^72^ Most simply, cryptic species are those that have been binned into a single taxon at the species rank. Discriminating cryptic species can enlarge estimates of biodiversity and provide insight into the mechanisms that underlie adaptation and speciation. Following suggestions and criticisms offered by Struck *et al.*,^73^ we provide here divergence times and molecular evidence that point to the existence of cryptic species within the genus *Tetrabaena*. This insight is reinforced by previously published biogeographic,^74,75^ phenotypic,^75^ and genetic data^76^ on this enigmatic genus.

*Tetrabaena* is a monotypic, colonial volvocine alga and one of two genera composing the multicellular Tetrabanaeceae. Several strains of this genus have been described as far back as 1841;^77^ all described *Tetrabaena* spp. have been of freshwater origin except one Antarctic strain, NIES-691. Following collection of the Antarctic *T. socialis* from King George Island,^78^ Nozaki and Ohtani^75^ provided morphological data and temperature growth profiles that demonstrated phenotypic idiosyncrasies of this strain in comparison to the freshwater NIES-571. Phenotypic traits distinctive to NIES-691 included: (*i*) vegetative colonies measuring up to 50 μm in diameter (∼18 μm larger than NIES-571), (*ii*) immobile colonies that adhere to surfaces at 20 °C, and (*iii*) failure to grow at 25 °C. In contrast to NIES-691, NIES-571 was observed to produce smaller colonies and to grow normally at all tested temperatures ranging from 5-25 °C, with no phenotypic changes that produce immobile colonies or ones adhering to surfaces.^75^ These phenotypic differences, which may be local environmental adaptation, point towards the possibility of a cryptic species in *Tetrabaena*, arguing for the examination of detailed molecular data on these two strains.

In 1997, Mai and Coleman^76^ produced complete Internal Transcribed Spacer 2 (ITS-2) sequences for 111 Volvocales. including *Tetrabaena socialis* strains NIES-571 and NIES-691. When these two *T. socialis* ITS-2 sequences are subjected to pairwise alignment, the percent identity between them is 92.47% (**Supplementary Table 3**). Similar to the percent identity (91.25%) between ITS-2 sequences of *V. gigas* UTEX 1895 and *V. powersii* UTEX 1864, two sister taxa recognized as separate species, when these are subject to pairwise alignment (**Supplementary Table 3**). When the same test is applied to ITS-2 sequences of *Volvox carteri f. nagariensis* strains (Poona and 72-52), the percent identity between them is 98.06% (**Supplementary Table 3**). Considered in relation to previously reported phenotypic data, these molecular differences lead us to hypothesize that NIES-571 and NIES-691 are cryptic species within the genus *Tetrabaena*.

This hypothesis is bolstered by our clock analysis and molecular data. The two strains appear to have diverged from one another ∼51 MYA (82-25 MYA) (**Figure 4**). This divergence time estimate stands in contrast to that estimated between strains of *Volvox carteri* f. *nagariensis* (NIES-865 and HK10) as well as to that estimated between strains of *Pleodorina starrii* (NIES-1362 and NIES-1363). In each of these two examples, strains are thought to have diverged from one another <1 MYA (**Figure 4**). And in both cases, there is good reason to believe that each strain pair is a biological species *sensu* Mayr.^79^ *V. carteri* NIES-865 is an isolate of *V. carteri* HK10^80,81^, and mating experiments^82^ between *Pleodorina starrii* strains (NIES-1362 and NIES-1363) produce fertile offspring. Exploration of conserved regions from our 16 amino acid sequence alignments used in our molecular clock analyses further highlight differences in sequence percent identity. Pairwise percent identity between *V. carteri* and *P. starrii* strains was 100% and 99.65-100%, respectively, whereas the percent identity between each set of *T. socialis* sequences ranges between 88-99.27% indicating substantial evolutionary differences in the protein sequences (**Supplementary Table 3**).

Given the divergence times indicated by our dataset, and the phenotypic and molecular features known to distinguish these *T. socialis* strains, we propose recognizing *Tetrabaena socialis* NIES-571 as a cryptic species to clarify *socialis* N-691’s current status as the type species. Formal elevation of NIES-571 to species status will require additional data. For example, complete plastid sequences should be obtained to complement the pairwise alignments of nuclear genes provided here. Because chloroplast genomes are known to be highly conserved,^83^ marked differences between plastid sequences of these two *T. socialis* strains would strengthen the case for one to be recognized as a cryptic species. Also, NIES-571 and NIES-691 should be crossed. And, even if mating experiments result in viable F1 progeny, additional crosses should be undertaken to assess the fertility of F1 hybrids in relation to their parents and to one another.

## Conclusion

Through the use of a more comprehensive molecular and taxonomic dataset, we have confirmed the major findings of Lindsey et al.’s phylogeny^14^, and further supported them by carrying out ancestral-state reconstructions using fossil-calibrated molecular clock analyses. We find that the origin of the crown TGV clade may be much older than previously thought, possibly emerging as early as the Carboniferous Period. Also, multicellularity in the Goniaceae + Volvocaceae evolved sometime between the Carboniferous-Triassic Periods, and multicellularity in Tetrabaenaceae may have evolved as recent as the Cretaceous Period. Armed with our time-tree and ancestral-state reconstructions, we reassessed Kirk’s 12-Step Program to differentiated multicellularity using a new, more robust phylogeny of the volvocine algae; we found the essential elements of his hypothesis to be correct. Finally, we suggest the possibility of a cryptic species in the multicellular Tetrabaenaceae, a possibility that could change its monotypic status, and provide an exciting opportunity to investigate the genomics of two diverged species that have adapted to very different environments.

In ancestral character state reconstructions, it is nearly always the case that multiple, nearly equally likely reconstructions exist, and this limits the confidence we can have in the specifics of the (marginally) most parsimonious. For example, in the clade at the bottom of Fig. 4, which includes *P. indica, P. starrii, V. powersii, V. gigas,* and some *E. elegans* strains, our reconstruction shows a gain of sterile soma at the base of the clade and a loss in *E. elegans*. However, it would be only slightly less parsimonious to show independent gains in *P. indica + P. starrii* and in *V. powersii* + *V. gigas*, with no loss. There is, though, no way to reconstruct the history of this trait that doesn’t involve several independent gains and/or losses.

The distinction between mortal somatic cells, which do not pass on genetic information to subsequent generations, and potentially immortal germ cells, which do, has been recognized as a fundamentally important evolutionary trait as far back as Weismann’s germ-plasm theory.^84^ The evolution of anisogamy is the first step in the differentiation of male and female sexes, setting the stage for sexual selection to produce all of the spectacular sexual dimorphisms we see throughout the animal kingdom^12,85,86^ as well as the more modest dimorphisms in some species of *Volvox* (dwarf males and extrafertile females). Although nearly equally likely reconstructions preclude a high degree of confidence in the details of these traits’ histories, this much is clear: given the long-recognized fundamental evolutionary significance of these traits, they are surprisingly evolutionarily labile, with multiple independent losses and/or gains, within the relatively recently-evolved multicellular volvocine algae.

## Materials and Methods

### Retrieval of genomic and RNA-Seq data for Archaeplastida phylogeny

A total of 164 taxa were sampled across Archaeplastida based upon publicly available genomic and RNA-Seq data. Specifically, longest primary transcript files were downloaded for 23 taxa from the Phytozome and Ensembl databases, and protein files for 7 taxa were downloaded from NCBI, OIST Marine Genomics Unit, OrcAE, and Tokyo Institute of Technology. RNA-Seq data was downloaded for 132 taxa, and *de novo* transcriptomes were assembled for each taxon (**Supplementary Table 3**).

### Quality control of RNA-Seq reads

Quality of all RNA-Seqs was initially checked using FastQC v.0.11.8 with an additional FastQC assessment post-trimming. Quality trimming was conducted with either Trimmomatic v.0.39^87^ or BBDuk from the BBMap suite^88^. BBDuk was utilized for quality trimming only if Trimmomatic failed to remove all adapter content. A 4-base sliding window approach was used to trim the rest of the read once average quality fell below a Phred score of 15; reads that were below a minimum length of 36 bases were discarded (LEADING: 3 TRAILING:3 SLIDINGWINDOW:4:15 MINLEN:36). To trim adapter content, the ILLUMINACLIP:2:30:10 option was used with the “adapters.fa” file provided in the BBMap suite due to its comprehensive list of adapter sequences. ILLUMINACLIP option allows for 2 “seed” mismatches where the seed is a short segment of the adapter that is being aligned in every section of the read. If >2 mismatches occurred, no trimming of the read occurred. Additionally, there had to be at least 30 matched bases in the paired-end palindrome read alignment and at least 10 matched bases between an adapter sequence and read. For BBDuk (ktrim=r k=25 mink=5 qtrim=r qhdist=1 minlength=36 trimq=15 tpe tbo), the “adapters.fa” file was also used as the adapter reference list.

### De novo transcriptome assembly

The quality filtered, paired-end reads were used to assemble *de novo* transcriptomes with SOAPdenovo-Trans v1.0.4^89^ using a k-mer size of 25 (SOAPdenovo-Trans-31mer all -s <config input file> -o <outfile> -K 25), GapCloser was used to close gaps in each *de novo* transcriptome (-b <soapdenovo-Trans_config_file> -a <.scafSeq file> -o <outfile> -l <max read length>), and CDHIT v4.8.1^90^ was used under default parameters to cluster redundant transcripts from each assembled transcriptome.

### Annotation of de novo transcripts

Each assembled transcript had its longest open reading frame predicted using TransDecoder v5.5.0 (TransDecoder.LongOrfs -t <path to transcript file>). After the longest open reading frames were predicted for each transcript, the “longest_orfs.pep” file created by TransDecoder was subsequently used in a BLASTP search conducted by Diamond v2.0.11.149^91^ (diamond blastp -p <number of threads> -q longest_orfs.pep -d uniref90 -k 1 -e 0.00001 -o <.tsv outfile> --very-sensitive) against the Uniref90 database. Additional Diamond BLASTP options “k”, “e”, and “very-sensitive” correspond to max number of target sequences, e-value, and the sensitivity of Diamond when aligning longer sequences, respectively. Protein domain identification was accomplished by using HMMER v3.2.1^92^ (hmmscan -cpu <number of threads> --domtblout <outfile> <path to protein database> longest_orfs.pep) to conduct Pfam searches against the Pfam-A database using the “longest_orfs.pep” file as our input. The most likely coding region from each open read frame was predicted using TransDecoder v5.5.0 (TransDecoder.Predict -t <path to transcript file> --retain_pfam_hits <path to domtblout file> --retain_blastp_hits <path to blastp outfile>) by incorporating the results from our Diamond BLASTP and HMMER homology searches. As a result, nucleotide CDS and amino acid PEP files were created.

### Orthologous gene identification infers 284 single-copy genes shared among red and green algae

A training set of 10 Archaeplastida taxa (1 rhodophyte, 5 streptophytes, and 4 chlorophytes) were chosen from the Phytozome database to infer a putative list of single-copy genes present across Rhodophyta, Streptophyta, and Chlorophyta (**Supplementary Table 4**). These 10 taxa were chosen based upon the availability of a longest primary transcript file, their respective clade in Archaeplastida, and genome assembly quality. A primary transcript file was available on Phytozome for each chosen organism, and this first criterion is perhaps the most important due to alternative splice variants being accounted for in each genome. This limited our ability, however, to choose more than one rhodophyte. Nevertheless, we were still able to sample from all three major clades of Archaeplastida to infer a putative list of single-copy genes. Lastly, 4 out of the 10 genomes sampled have chromosome-level assembly and annotation, and a 5^th^, *Micromonas pusilla*, is in the targeted finish state with a 4 chromosome-level assembly with no gaps.

The primary transcript file for each taxon in our training set was used to determine a putative list of single-copy genes using Orthofinder v2.3.8^93^ (orthofinder -M msa -oa -f <path to transcript files>). Orthofinder returned 284 orthologous groups where all species were present and all genes single-copy. These 284 orthologous genes across all species were aligned using MAFFT v7.487^94^. HMMER (for f in $(ls *.fa); do echo “Processing file: ${f}”; hmmbuild ${f}.hmm ${f}; done) was subsequently utilized to build profile Hidden Markov Model (HMM) files for each gene alignment. These profile HMMs were then used by Orthofisher^95^ to mine our genomes and *de novo* transcriptomes for those single-copy genes.

### Ruling out putative organellar sequences from inferred single-copy gene dataset

All sequences identified as single-copy from our 10 genome training set were BLASTed against the NCBI non-redundant (nr) protein database (diamond makedb --in nr.gz --taxonmap prot.accession2taxid.FULL --taxonnodes nodes.dmp --taxonnames names.dmp -d nr) to ensure protein sequences identified as single-copy were also of nuclear origin using Diamond v2.0.11.149^91^ (diamond blastp -p 8 -q <query_file> -d nr -k 10 --outfmt 6 qseqid sseqid stitle staxids sscinames sskingdoms pident length mismatch gapopen qstart qend sstart send evalue bitscore -o <outfile> --very-sensitive --header). In total, 174 sequences contained at least one of the following terms identifying them as possible organellar sequences: “chloroplastic”, “chloroplast”, “mitochondrial” and/or “mitochondria”.

To address whether these sequences were of nuclear or organellar origin, GenPept files were fetched from NCBI using the protein accessions from the BLAST outfiles. For taxa, such as *Arabidopsis thaliana* and *Oryza sativa*, which have chromosome-level genome assemblies, all putative organellar proteins were shown to originate from the nuclear genome. An additional step of BLASTing these questionable sequences against the complete chloroplast genome found in the NCBI chloroplast database from either the species or genus of the 10 taxa chosen for orthologous, single-copy gene identification. For those sequences, all BLAST results resulted in poor matches, high e-values, and low bit scores, indicating that these sequences are not originating from the plastid genome. For the putative mitochondrial sequences in our dataset, complete mitochondrial genomes for all 10 taxa could not be located. For those which could be found in the NCBI mitochondrial genome database, those sequences were BLASTed against its organism’s mitochondrial genome. For all others, a broad BLAST search was done against all mitochondrial genomes. Again, all BLAST results indicated poor matches including high e-values and low bit scores for these sequences.

### Gene sequence alignments

Once the amino acid sequences were confirmed to be of nuclear origin and single-copy, they were aligned using MUSCLE v3.8.31, resulting in 284 untrimmed gene alignments. Furthermore, each alignment was manually inspected and aligned in Aliview v1.26^96^. During manual inspection poorly aligned sequences that included bacterial, fungal, and non-algal sequences were removed according to the taxon ID assigned to a particular sequence’s top BLAST result. Additionally, poorly aligned sequences that BLASTed to a similar sequence but different protein than the other well-aligned sequences were removed from each alignment to ensure that the same protein product was being compared and analyzed across taxa. Following manual quality control of alignments, ambiguously aligned regions from each alignment were trimmed using Trimal v1.4.1^97^ (trimal -in <infile> -automated1 -out <outfile> -htmlout <html_outfile>) allowing for only conserved and reliably aligned regions to later undergo phylogenetic analysis.

### Parameters for Phylogenetic analyses

Each single-gene alignment had its best evolutionary model predicted by ProtTest v3.4.2^98^ under the Akaike Information Criterion (AIC). Each predicted evolutionary model and its associated alignment may be found in **Supplementary Table 5**. All single-gene alignments underwent Maximum-Likelihood (ML) analysis using IQtree^99^ (iqtree -nt AUTO -s <input> -m <evolutionary_model> -bb 1000 -pre <outfile>).

Phyutility v2.7.1^100^ (phyutility -concat -in <infiles> -out <concatenated_outfile>) was utilized to concatenate all gene alignment files for ML and BI analyses. IQtree was used to run a concatenated, gene-partitioned ML analysis with 1000 rapid bootstrap replicates^101^ (iqtree -nt AUTO -s <aln_file> -spp <partition_file> -bb 1000 -pre <outfile>). The gene-partition strategy was based upon the best evolutionary model predicted by ProtTest. The Bayesian inference (BI) analysis was conducted with MrBayes 3.2.7a^102^ under the same gene-partitioning strategy previously described. A total of 4 runs each with 4 Markov chains (3 heated and 1 cold). Trees were sampled every 1000 generations over 1,000,000 generations during which convergence (average standard deviation of split frequencies = <0.01) between runs was reached, and a burn-in of 25% was used (ngen=1000000 nruns=4 samplefreq=1000 nchains=4 starttree=random).

Lastly, ASTRAL^103^ was utilized to perform a coalescent-based analysis using the 284 single-gene trees produced by IQtree. Each single-gene tree had branches with low bootstrap support (<10) collapsed using Newick Utilities v1.6^104^.

### Parameters for molecular clock analyses

For all clock analyses, concatenated datasets of 8, 12, and 16 genes were used to offset high computational cost. Each reduced molecular dataset was formed based on SortaDate^105^ results of our original XYZ gene dataset where each single-gene phylogeny is scored and ranked on similarity to a given species tree (bipartition score), least amount of root-to-tip variance (clock-likeness), and overall tree length (calculable amount of molecular evolution). For each of the smaller datasets, genes were ranked with an emphasis on their Sortadate bipartition score with the scores ranging from: 48-56% (8 genes), 46-56% (12 gene), and 45-56% (16 genes) (**Supplementary Table 6**).

Phylobayes 4.1b^51^ was used to perform relaxed molecular clock analyses on each dataset under lognormal autocorrelated (-ln) and uncorrelated gamma (-ugam) models with the LG protein substitution model applied (-lg), discrete gamma rate across sites (-dgam 4), and birth-death prior on divergence times (-bd). The red algae were specified as the outgroup taxa according to manual instructions (-r <outgroup.txt>). A total of 17 nodes were calibrated for each molecular clock analysis using fossil taxa with either a minimum bound or minimum + maximum bounds (**Figure 2**) with the soft bounds flag (-sb) allowing date sampling to occur slightly outside of the specified bounds (5% on a pure minimum date, and 2.5% on lower and upper bounds). No secondary node calibrations were used in this study. The root age (red and green algal divergence) was set to a maximum bound of 2000 MY. Separate runs for our maximum-likelihood and coalescent-based species tree topologies were conducted. A total of two MCMC chains were conducted per run for a minimum of 10,000 generations, and the convergence of the two chains was determined in Phylobayes. A summarization of the chains was done after discarding the first 2000 generations (∼20%) as burn-in.

### Ancestral state reconstructions of discrete volvocine characters

Two programs were used to estimate the probability of ancestral character states at internal nodes in the volvocine algae: the Phytools^106^ R package and Mr. Bayes^102^. For each of the two programs, our tree topology was constrained to the maximum-likelihood, molecular-clock tree inferred using 16 protein-coding genes under the CIR model. For each ASR method, the following discrete character states were ancestrally reconstructed in the volvocine algae: (*i*) cellularity, (*ii*) spheroidal body plan, (*iii*) developmental traits identified by Kirk in his 12-Step Program^31^, and (*iv*) 7 sexual traits from two previous studies.^29,30^

In Phytools, stochastic character mapping (“simmap”) was applied to estimate probabilities of ancestral character states. For each analysis, the equal rates (“ER”) and all-rates-different (“ARD”) models of character evolution were compared using the R “anova” function. When used, this function outputs Akaike Information Criterion (AIC) values and Akaike weights for each model, and a higher Akaike weight indicates the favored model.^107^ The highest weighted model was used for each “simmap” analysis where a total of 1000 stochastic simulations (nsim=1000) were conducted under an empirical Bayes method (Q=”empirical”). To conclude each run, the 1000 stochastic simulations were summarized into a single result, or tree, with calculated posterior probabilities as a pie chart at each internal node. The R script, input tree file, and input trait CSV file to replicate each run is available at Dryad.

To further analyze ancestral character states in the volvocine algae, the MBASR toolkit^108^, an R package, was used to estimate the previously mentioned discrete characters. The MBASR toolkit aims to circumvent the abstruse nature of ASR programs by automating the ASR workflow for discrete traits in Mr. Bayes.^102^ For each ancestral state reconstruction, a Markov chain Monte Carlo (MCMC) simulation was conducted for 1,000,000 generations (MBASR n.samples=10,000) with a sample taken every 100 generations. Burn-in is automatically set by MBASR to discard samples whose log likelihoods score below the upper 25% threshold. All runs were conducted as “unordered” unless specified. The R script, input tree file, and input trait CSV file to replicate each run is available at Dryad.

## Supporting information

Strain information

Pairwise sequence identities of ITS seqs

Pairwise sequence identities of ITS seqs between Tetrabaena, Pleodorina, and Volvox strains

Fossil cross validation tests

Ancestral state reconstruction trees

## Acknowledgments

The authors gratefully acknowledge the useful discussion of *Volvox* germ specialization provided by Hisayoshi Nozaki and Alexey Desnitskiy, as well as editorial suggestions provided by James Elser, Scott Miller, Gavin Sherlock. This material is based upon work while MDH was serving at the National Science Foundation.

## Author’s Contributions

Performed the experiments (CRL), carried out the analyses (CRL, MDH), wrote the manuscript (CRL, AHK, MDH, RFR). All authors have read and approved the final manuscript.

