## Supplementary material for "Fossil-calibrated inference of divergence times among the Volvocine algae enables reconstruction of the steps that led to differentiated multicellularity": Strain information

**Supplementary Table 1.** Strain information for sampled taxa

| Division | Class or Order | Species | Source | Type |
|---|---|---|---|---|
| Rhodophyta | Cyanidiophyceae | Cyanidioschyzon merolae | Ensembl | Proteome |
| Rhodophyta | Cyanidiophyceae | Galdieria sulphuraria | Ensembl | Proteome |
| Rhodophyta | Cyanidiophyceae | Cyanidiococcus yangmingshanensis | NCBI | Proteome |
| Rhodophyta | Bangiophyceae | Porphyra umbilicalis | JGI | Genome |
| Rhodophyta | Bangiophyceae | Pyropia yezoensis | 1kp-ERS1830191 | Transcriptome |
| Rhodophyta | Florideophyceae | Chondrus crispus | Ensembl | Proteome |
| Rhodophyta | Florideophyceae | Ahnfeltiopsis flabelliformis | 1kp-ERS1830200 | Transcriptome |
| Rhodophyta | Florideophyceae | Gloiopeltis furcata | 1kp-ERS1830197 | Transcriptome |
| Rhodophyta | Florideophyceae | Mazzaella japonica | 1kp-ERS1830199 | Transcriptome |
| Rhodophyta | Florideophyceae | Grateloupia livida | 1kp-ERS1830209 | Transcriptome |
| Rhodophyta | Florideophyceae | Grateloupia catenata | 1kp-ERS1830211 | Transcriptome |
| Rhodophyta | Florideophyceae | Grateloupia turuturu | 1kp-ERS1830210 | Transcriptome |
| Streptophyta | Klebsormidiophyceae | Klebsormidium nitens | Tokyo Institute of Technology | Proteome |
| Streptophyta | Charophyceae | Nitella mirabilis | NCBI-PRJNA158153 | Transcriptome |
| Streptophyta | Charophyceae | Chara braunii | Ensembl | Proteome |
| Streptophyta | Charophyceae | Chara globularis | Vries et al. (2018) | Transcriptome |
| Streptophyta | Coleochaetophyceae | Coleochaete orbicularis | NCBI-SRS10979585 | Transcriptome |
| Streptophyta | Coleochaetophyceae | Coleochaete scutata | Vries et al. (2018) | Transcriptome |
| Streptophyta | Zygnematophyceae | Staurastrum sebaldi | 1kp-ERS1830172 | Transcriptome |
| Streptophyta | Zygnematophyceae | Cosmarium granatum | 1kp-ERS1830163 | Transcriptome |
| Streptophyta | Zygnematophyceae | Euastrum affine | 1kp-ERS1830167 | Transcriptome |
| Streptophyta | Zygnematophyceae | Cylindrocystis cushleckae | 1kp-ERS368241 | Transcriptome |
| Streptophyta | Zygnematophyceae | Mesotaenium kramstae | 1kp-ERS1830178 | Transcriptome |
| Streptophyta | Zygnematophyceae | Zygnemopsis sp. | 1kp-ERS3670390 | Transcriptome |
| Streptophyta | Jungermanniopsida | Radula lindenbergiana | 1kp-ERS1830058 | Transcriptome |
| Streptophyta | Jungermanniopsida | Frullania sp. | 1kp-ERS3670367 | Transcriptome |
| Streptophyta | Jungermanniopsida | Scapania nemorea | 1kp-ERS1830046 | Transcriptome |
| Streptophyta | Jungermanniopsida | Barbilophozia barbata | 1kp-ERS1830047 | Transcriptome |
| Streptophyta | Marchantiopsida | Marchantia polymorpha | JGI | Genome |
| Streptophyta | Lycopodiopsida | Selaginella moellendorffii | JGI | Genome |
| Streptophyta | Lycopodiopsida | Isoetes tegetiformans | 1kp- ERS1829935 | Transcriptome |
| Streptophyta | Lycopodiopsida | Phylloglossum drummondii | 1kp- ERS1829929 | Transcriptome |
| Streptophyta | Lycopodiopsida | Huperzia lucidula | 1kp- ERS1829926 | Transcriptome |
| Streptophyta | Lycopodiopsida | Lycopodium annotinum | 1kp- ERS1829933 | Transcriptome |
| Streptophyta | Polypodiopsida | Polystichum tripteron | Qi et al. (2018) | Transcriptome |
| Streptophyta | Polypodiopsida | Cyrtomium fortunei | Qi et al. (2018) | Transcriptome |
| Streptophyta | Polypodiopsida | Dryopteris decipiens | Qi et al. (2018) | Transcriptome |
| Streptophyta | Polypodiopsida | Ctenitis subglandulosa | Qi et al. (2018) | Transcriptome |
| Streptophyta | Pinopsida | Thuja plicata | JGI | Genome |
| Streptophyta | Pinopsida | Pinus hwangshanensis | Jin et al. (2021) | Transcriptome |
| Streptophyta | Ginkgoopsida | Ginkgo biloba | 1kp- ERS368269 | Transcriptome |
| Streptophyta | Amborellales | Amborella trichopoda | JGI | Genome |
| Streptophyta | Nymphaeles | Nymphaea colorata | JGI | Genome |
| Streptophyta | Nymphaeles | Nymphaea caerulea | Zhang et al. (2019) | Transcriptome |
| Streptophyta | Laurales | Cinnamomum kanehirae | JGI | Genome |
| Streptophyta | Laurales | Peumus boldus | 1kp-ERS1829196 | Transcriptome |
| Streptophyta | Laurales | Gyrocarpus americanus | 1kp-ERS1829190 | Transcriptome |
| Streptophyta | Chloranthales | Sarcandra glabra | 1kp-ERS368201 | Transcriptome |
| Streptophyta | Chloranthales | Ascarina rubricaulis | 1kp-ERS1829208 | Transcriptome |

| | | | | |
|---|---|---|---|---|
| Streptophyta | Brassicales | Arabidopsis thaliana | JGI | Genome |
| Streptophyta | Brassicales | Capsella rubella | JGI | Genome |
| Streptophyta | Brassicales | Eutrema salsugineum | JGI | Genome |
| Streptophyta | Brassicales | Brassica rapa | JGI | Genome |
| Streptophyta | Poales | Brachypodium distachyon | JGI | Genome |
| Streptophyta | Poales | Oryza sativa | JGI | Genome |
| Streptophyta | Poales | Sorghum bicolor | JGI | Genome |
| Streptophyta | Poales | Zea Mays | JGI | Genome |
| Chlorophyta | Mamiellophyceae | Bathycoccus prasinos | NCBI | Proteome |
| Chlorophyta | Mamiellophyceae | Ostreococcus tauri | JGI | Genome |
| Chlorophyta | Mamiellophyceae | Micromonas pusilla | NCBI | Proteome |
| Chlorophyta | Chlorodendrophyceae | Tetraselmis cordiformis | 1kp-ERS1830070 | Transcriptome |
| Chlorophyta | Chlorodendrophyceae | Tetraselmis striata | NCBI-SRS10979578 | Transcriptome |
| Chlorophyta | Trebouxiophyceae | Coccomyxa subellipsoidea | JGI | Genome |
| Chlorophyta | Trebouxiophyceae | Botryococcus braunii | 1kp-ERS1830127 | Transcriptome |
| Chlorophyta | Trebouxiophyceae | Botryococcus terribilis | 1kp-ERS1830129 | Transcriptome |
| Chlorophyta | Trebouxiophyceae | Chlorella variabilis | NCBI | Proteome |
| Chlorophyta | Ulvophyceae | Ulva mutabilis | OrcAE; De Clerck et al. (2018) | Proteome |
| Chlorophyta | Ulvophyceae | Percursaria percursa | 1kp-ERS1830140 | Transcriptome |
| Chlorophyta | Ulvophyceae | Ochlochaete sp. | 1kp-ERS3670385 | Transcriptome |
| Chlorophyta | Ulvophyceae | Entocladia endozoica | 1kp-ERS1830144 | Transcriptome |
| Chlorophyta | Ulvophyceae | Acrosiphonia sp. | 1kp-ERS3670386 | Transcriptome |
| Chlorophyta | Ulvophyceae | Helicodictyon planctonicum | 1kp-ERS1830074 | Transcriptome |
| Chlorophyta | Ulvophyceae | Hazenia basiliensis | Hou et al. (2022) | Transcriptome |
| Chlorophyta | Ulvophyceae | Rhexinema paucicellulare | Hou et al. (2022) | Transcriptome |
| Chlorophyta | Ulvophyceae | Planophila laetevirens | 1kp-ERS1830148 | Transcriptome |
| Chlorophyta | Ulvophyceae | Planophila sp | 1kp-ERS1830149 | Transcriptome |
| Chlorophyta | Ulvophyceae | Halochlorococcum marinum | 1kp-ERS1830151 | Transcriptome |
| Chlorophyta | Ulvophyceae | Oltmannsiellopsis unicellularis | NCBI- SRS10979569 | Transcriptome |
| Chlorophyta | Ulvophyceae | Oltmannsiellopsis viridis | 1kp-ERS1830142,ERS1830143 | Transcriptome |
| Chlorophyta | Ulvophyceae | Ignatius tetrasporus | 1kp- ERS1830151 | Transcriptome |
| Chlorophyta | Ulvophyceae | Caulerpa lentillifera | Arimoto et al. (2019) | Transcriptome |
| Chlorophyta | Ulvophyceae | Caulerpa taxifolia | Ranjan et al. (2015) | Transcriptome |
| Chlorophyta | Ulvophyceae | Caulerpa cylindracea | Unlu et al. (2019) | Transcriptome |
| Chlorophyta | Ulvophyceae | Codium fragile | 1kp-ERS1830154 | Transcriptome |
| Chlorophyta | Ulvophyceae | Bryopsis plumosa | 1kp-ERS1830153 | Transcriptome |
| Chlorophyta | Ulvophyceae | Bryopsis hypnoides | Hou et al. (2022) | Transcriptome |
| Chlorophyta | Ulvophyceae | Ostreobium quekettii | Hou et al. (2022) | Transcriptome |
| Chlorophyta | Chlorophyceae | Oedogonium cardiacum | 1kp-ERS1830083 | Transcriptome |
| Chlorophyta | Chlorophyceae | Oedogonium foveolatum | 1kp-ERS1830084 | Transcriptome |
| Chlorophyta | Chlorophyceae | Chaetopeltis orbicularis | 1kp-ERS1830072 | Transcriptome |
| Chlorophyta | Chlorophyceae | Stigeoclonium helveticum | 1kp-ERS1830076 | Transcriptome |
| Chlorophyta | Chlorophyceae | Aphanochaete repens | 1kp-ERS1830078 | Transcriptome |
| Chlorophyta | Chlorophyceae | Scenedesmus glucoliberatum PABB004 | Mancipe et al (2021) | Transcriptome |
| Chlorophyta | Chlorophyceae | Pediastrum duplex | 1kp-ERS3670374 | Transcriptome |
| Chlorophyta | Chlorophyceae | Chromochloris zofingiensis | JGI | Genome |
| Chlorophyta | Chlorophyceae | Golenkinia longispicula | 1kp-ERS1830080 | Transcriptome |
| Chlorophyta | Chlorophyceae | *Astrephomene gubernaculifera* NIES-418 | Lindsey et al. (2021) | Transcriptome |

| | | | | |
|---|---|---|---|---|
| Chlorophyta | Chlorophyceae | *Astrephomene perforate* NIES-564 | Lindsey et al. (2021) | Transcriptome |
| Chlorophyta | Chlorophyceae | *Basichlamys sacculifera* NIES-566 | Lindsey et al. (2021) | Transcriptome |
| Chlorophyta | Chlorophyceae | *Colemanosphaera charkowiensis* NIES-3383 | Lindsey et al. (2021) | Transcriptome |
| Chlorophyta | Chlorophyceae | *Chlamydomonas debaryana* SAG 11-55a | Lindsey et al. (2021) | Transcriptome |
| Chlorophyta | Chlorophyceae | *Chlamydomonas debaryana* SAG 70.81 | Lindsey et al. (2021) | Transcriptome |
| Chlorophyta | Chlorophyceae | *Chlamydomonas globosa* SAG 81.72 | Lindsey et al. (2021) | Transcriptome |
| Chlorophyta | Chlorophyceae | *Chlamydomonas moewusii* SAG 11-16f | Lindsey et al. (2021) | Transcriptome |
| Chlorophyta | Chlorophyceae | *Chlamydomonas schloesseri* | Lindsey et al. (2021) | Transcriptome |
| Chlorophyta | Chlorophyceae | *Eudorina cylindrica* NIES-722 | Lindsey et al. (2021) | Transcriptome |
| Chlorophyta | Chlorophyceae | *Eudorina elegans* NIES-456 | Lindsey et al. (2021) | Transcriptome |
| Chlorophyta | Chlorophyceae | *Eudorina elegans* NIES-458 | Lindsey et al. (2021) | Transcriptome |
| Chlorophyta | Chlorophyceae | *Eudorina elegans* NIES-568 | Lindsey et al. (2021) | Transcriptome |
| Chlorophyta | Chlorophyceae | *Eudorina elegans* NIES-717 | Lindsey et al. (2021) | Transcriptome |
| Chlorophyta | Chlorophyceae | *Eudorina elegans* NIES-719 | Lindsey et al. (2021) | Transcriptome |
| Chlorophyta | Chlorophyceae | *Eudorina elegans* NIES-720 | Lindsey et al. (2021) | Transcriptome |
| Chlorophyta | Chlorophyceae | *Eudorina illinoisensis* NIES-720 | Lindsey et al. (2021) | Transcriptome |
| Chlorophyta | Chlorophyceae | *Eudorina minodii* NIES-856 | Lindsey et al. (2021) | Transcriptome |
| Chlorophyta | Chlorophyceae | *Eudorina peripheralis* NIES-725 | Lindsey et al. (2021) | Transcriptome |
| Chlorophyta | Chlorophyceae | *Eudorina unicocca* SAG 24-1c | Lindsey et al. (2021) | Transcriptome |
| Chlorophyta | Chlorophyceae | *Gonium multicoccum* NIES-737 | Lindsey et al. (2021) | Transcriptome |
| Chlorophyta | Chlorophyceae | *Gonium octonarium* NIES-851 | Lindsey et al. (2021) | Transcriptome |
| Chlorophyta | Chlorophyceae | *Gonium quadratum* NIES-653 | Lindsey et al. (2021) | Transcriptome |
| Chlorophyta | Chlorophyceae | *Gonium viridistellatum* NIES-654 | Lindsey et al. (2021) | Transcriptome |
| Chlorophyta | Chlorophyceae | *Pandorina colemaniae* NIES-572 | Lindsey et al. (2021) | Transcriptome |
| Chlorophyta | Chlorophyceae | *Pandorina morum* NIES-890 | Lindsey et al. (2021) | Transcriptome |
| Chlorophyta | Chlorophyceae | *Platydorina caudata* NIES-728 | Lindsey et al. (2021) | Transcriptome |
| Chlorophyta | Chlorophyceae | *Pleodorina indica* NIES-736 | Lindsey et al. (2021) | Transcriptome |
| Chlorophyta | Chlorophyceae | *Pleodorina japonica* UTEX 2523 | Lindsey et al. (2021) | Transcriptome |
| Chlorophyta | Chlorophyceae | *Pleodorina starrii* NIES-1362 | Lindsey et al. (2021) | Transcriptome |
| Chlorophyta | Chlorophyceae | *Pleodorina starrii* NIES-1363 | Lindsey et al. (2021) | Transcriptome |
| Chlorophyta | Chlorophyceae | *Pleodorina thompsonii* NIES-4126 | Lindsey et al. (2021) | Transcriptome |

| | | | | |
|---|---|---|---|---|
| Chlorophyta | Chlorophyceae | *Vitreochlamys aulata* NIES-878 | Lindsey et al. (2021) | Transcriptome |
| Chlorophyta | Chlorophyceae | *Vitreochlamys aulata* SAG 80.81 | Lindsey et al. (2021) | Transcriptome |
| Chlorophyta | Chlorophyceae | *Vitreochlamys nekrassovii* SAG 11-10 | Lindsey et al. (2021) | Transcriptome |
| Chlorophyta | Chlorophyceae | *Vitreochlamys ordinate* NIES-882 | Lindsey et al. (2021) | Transcriptome |
| Chlorophyta | Chlorophyceae | *Volvox africanus* NIES-863 | Lindsey et al. (2021) | Transcriptome |
| Chlorophyta | Chlorophyceae | *Vitreochlamys aureus* NIES-541 | Lindsey et al. (2021) | Transcriptome |
| Chlorophyta | Chlorophyceae | *Vitreochlamys barberi* NIES-730 | Lindsey et al. (2021) | Transcriptome |
| Chlorophyta | Chlorophyceae | *Volvox carteri* f. *kawasakiensis* NIES-732 | Lindsey et al. (2021) | Transcriptome |
| Chlorophyta | Chlorophyceae | *Volvox carteri* f. *nagariensis* NIES-865 | Lindsey et al. (2021) | Transcriptome |
| Chlorophyta | Chlorophyceae | *Volvox carteri* f. *weismannia* NIES-866 | Lindsey et al. (2021) | Transcriptome |
| Chlorophyta | Chlorophyceae | *Volvox dissipatrix* NIES-4128 | Lindsey et al. (2021) | Transcriptome |
| Chlorophyta | Chlorophyceae | *Volvox ferrisii* NIES-3986 | Lindsey et al. (2021) | Transcriptome |
| Chlorophyta | Chlorophyceae | *Volvox gigas* NIES-867 | Lindsey et al. (2021) | Transcriptome |
| Chlorophyta | Chlorophyceae | *Volvox globator* SAG 199.80 | Lindsey et al. (2021) | Transcriptome |
| Chlorophyta | Chlorophyceae | *Volvox kirkiorum* NIES-543 | Lindsey et al. (2021) | Transcriptome |
| Chlorophyta | Chlorophyceae | *Volvox obversus* NIES-868 | Lindsey et al. (2021) | Transcriptome |
| Chlorophyta | Chlorophyceae | *Volvox ovalis* NIES-2569 | Lindsey et al. (2021) | Transcriptome |
| Chlorophyta | Chlorophyceae | *Volvox powersii* NIES-4127 | Lindsey et al. (2021) | Transcriptome |
| Chlorophyta | Chlorophyceae | *Volvox tertius* NIES-544 | Lindsey et al. (2021) | Transcriptome |
| Chlorophyta | Chlorophyceae | *Volvulina boldii* NIES-893 | Lindsey et al. (2021) | Transcriptome |
| Chlorophyta | Chlorophyceae | *Volvulina compacta* NIES-582 | Lindsey et al. (2021) | Transcriptome |
| Chlorophyta | Chlorophyceae | *Volvulina pringsheimii* NIES-895 | Lindsey et al. (2021) | Transcriptome |
| Chlorophyta | Chlorophyceae | *Volvulina steinii* SAG 90-1 | Lindsey et al. (2021) | Transcriptome |
| Chlorophyta | Chlorophyceae | *Colemanosphaera angeleri* FACHB 2363 | Hu et al. (2020) | Transcriptome |
| Chlorophyta | Chlorophyceae | *Colemanosphaera charkowiensis* FACHB 2326 | Hu et al. (2020) | Transcriptome |
| Chlorophyta | Chlorophyceae | *Eudorina cylindrica* FACHB 2322 | Hu et al. (2020) | Transcriptome |
| Chlorophyta | Chlorophyceae | *Eudorina elegans* FACHB 2321 | Hu et al. (2020) | Transcriptome |
| Chlorophyta | Chlorophyceae | *Pandorina colemaniae* FACHB 2361 | Hu et al. (2020) | Transcriptome |
| Chlorophyta | Chlorophyceae | *Pandorina morum* FACHB 2362 | Hu et al. (2020) | Transcriptome |
| Chlorophyta | Chlorophyceae | *Tetrabaena socialis* NIES-571 | Featherston et al. (2018) | Transcriptome |
| Chlorophyta | Chlorophyceae | *Tetrabaena socialis* NIES-691 | Zhang et al. (2019) | Transcriptome |
| Chlorophyta | Chlorophyceae | *Volvulina compacta* FACHB 2337 | Hu et al. (2020) | Transcriptome |

| | | | | |
|---|---|---|---|---|
| Chlorophyta | Chlorophyceae | *Yamagishiella unicocca* FACHB 2364 | Hu et al. (2020) | Transcriptome |
| Chlorophyta | Chlorophyceae | *Chlamydomonas reinhardtii* | NCBI | Proteome |
| Chlorophyta | Chlorophyceae | *Gonium pectorale* | NCBI | Proteome |
| Chlorophyta | Chlorophyceae | *Volvox carteri* f. *nagariensis* | NCBI | Proteome |
