## Supplementary material for "Fossil-calibrated inference of divergence times among the Volvocine algae enables reconstruction of the steps that led to differentiated multicellularity": Pairwise sequence identities of ITS seqs

**Supplementary Table 3.** ITS pairwise sequence comparisons via NCBI BLAST between *Tetrabaena* and *Volvox* strains for genes used in the present study.

| Gene Alignment | Taxa | Perc. Identity |
|---|---|---|
| ITS | T. socialis NIES-571 & NIES-691 | 92.47 |
| | P. starrii NIES-1362 & NIES-1363 | 91.25 |
| | V. carteri f. nagariensis NIES-865 & HK10 | 98.06 |
