## Supplementary material for "Fossil-calibrated inference of divergence times among the Volvocine algae enables reconstruction of the steps that led to differentiated multicellularity": Pairwise sequence identities of ITS seqs between Tetrabaena, Pleodorina, and Volvox strains

**Supplementary Table 4.** Pairwise sequence comparisons between *Tetrabaena*, *Pleodorina*, and *Volvox* strains for genes used in the present study.

| Gene Alignment | Taxa | Perc. Identity | Perc. Similarity |
|---|---|---|---|
| OG0004441 | T. socialis NIES-571 & NIES-691 | 97.08 | 97.92 |
| | P. starrii NIES-1362 & NIES-1363 | 100 | 100 |
| | V. carteri f. nagariensis NIES-865 & HK10 | 100 | 100 |
| OG0004570 | T. socialis NIES-571 & NIES-691 | 95.86 | 96.55 |
| | P. starrii NIES-1362 & NIES-1363 | 100 | 100 |
| | V. carteri f. nagariensis NIES-865 & HK10 | 100 | 100 |
| OG0004572 | T. socialis NIES-571 & NIES-691 | 90.72 | 98.84 |
| | P. starrii NIES-1362 & NIES-1363 | N/A | N/A |
| | V. carteri f. nagariensis NIES-865 & HK10 | 100 | 100 |
| OG0004581 | T. socialis NIES-571 & NIES-691 | N/A | N/A |
| | P. starrii NIES-1362 & NIES-1363 | 100 | 100 |
| | V. carteri f. nagariensis NIES-865 & HK10 | 100 | 100 |
| OG0004602 | T. socialis NIES-571 & NIES-691 | 99 | 99.5 |
| | P. starrii NIES-1362 & NIES-1363 | 100 | 100 |
| | V. carteri f. nagariensis NIES-865 & HK10 | 100 | 100 |
| OG0004910 | T. socialis NIES-571 & NIES-691 | N/A | N/A |
| | P. starrii NIES-1362 & NIES-1363 | 100 | 100 |
| | V. carteri f. nagariensis NIES-865 & HK10 | 100 | 100 |
| OG0004914 | T. socialis NIES-571 & NIES-691 | 97.79 | 99.12 |
| | P. starrii NIES-1362 & NIES-1363 | 100 | 100 |
| | V. carteri f. nagariensis NIES-865 & HK10 | 100 | 100 |
| OG0004916 | T. socialis NIES-571 & NIES-691 | N/A | N/A |
| | P. starrii NIES-1362 & NIES-1363 | 100 | 100 |
| | V. carteri f. nagariensis NIES-865 & HK10 | 100 | 100 |
| OG0005090 | T. socialis NIES-571 & NIES-691 | N/A | N/A |
| | P. starrii NIES-1362 & NIES-1363 | 100 | 100 |
| | V. carteri f. nagariensis NIES-865 & HK10 | 100 | 100 |
| OG0005111 | T. socialis NIES-571 & NIES-691 | 97.79 | 98.69 |
| | P. starrii NIES-1362 & NIES-1363 | 100 | 100 |
| | V. carteri f. nagariensis NIES-865 & HK10 | 100 | 100 |
| OG0005154 | T. socialis NIES-571 & NIES-691 | N/A | N/A |
| | P. starrii NIES-1362 & NIES-1363 | N/A | N/A |
| | V. carteri f. nagariensis NIES-865 & HK10 | 100 | 100 |
| OG0005180 | T. socialis NIES-571 & NIES-691 | 97.43 | 98.2 |
| | P. starrii NIES-1362 & NIES-1363 | 100 | 100 |
| | V. carteri f. nagariensis NIES-865 & HK10 | 100 | 100 |
| OG0005188 | T. socialis NIES-571 & NIES-691 | 97.69 | 98.85 |
| | P. starrii NIES-1362 & NIES-1363 | 100 | 100 |

| | | | |
|---|---|---|---|
| | V. carteri f. nagariensis NIES-865 & HK10 | 100 | 100 |
| OG0005211 | T. socialis NIES-571 & NIES-691 | 88.68 | 92.45 |
| | P. starrii NIES-1362 & NIES-1363 | 100 | 100 |
| | V. carteri f. nagariensis NIES-865 & HK10 | 100 | 100 |
| OG0005213 | T. socialis NIES-571 & NIES-691 | N/A | N/A |
| | P. starrii NIES-1362 & NIES-1363 | 100 | 100 |
| | V. carteri f. nagariensis NIES-865 & HK10 | 100 | 100 |
| OG0005213 | T. socialis NIES-571 & NIES-691 | 99.27 | 99.27 |
| | P. starrii NIES-1362 & NIES-1363 | 99.65 | 99.65 |
| | V. carteri f. nagariensis NIES-865 & HK10 | 100 | 100 |
