## Supplementary material for "Fossil-calibrated inference of divergence times among the Volvocine algae enables reconstruction of the steps that led to differentiated multicellularity": Fossil cross validation tests

| | | Normal run | | | |
|---|---|---|---|---|---|
| | Node | Mean date | Lower 95% | Upper 95% | Fossil age c |
| 8 genes, CB | | | | | |
| CIR | cyamer_Ensembl_chocri_Ensembl | 1108.77 | 989.85 | 1319.82 | MIN: 1047 |
| | graliv_1KPTI_chocri_Ensembl | 636.282 | 559.753 | 722.057 | MIN: 609 |
| | euaaff_1KPTI_zygsp_1KPTI | 556.882 | 421.832 | 668.861 | MIN: 350 |
| | marpol_Phytozome_zeamay_Phytozome | 467.31 | 449.339 | 493.85 | MIN:480 |
| | isoteg_1KPTI_zeamay_Phytozome | 421.59 | 419.16 | 423.083 | 423-419 |
| | ctesub_Qi_zeamay_Phytozome | 406.314 | 400.308 | 411.428 | MIN: 385 |
| | ginbil_1KPTI_zeamay_Phytozome | 324.392 | 322.713 | 328.349 | 330-323 |
| | ambtri_Phytozome_zeamay_Phytozome | 128.213 | 125.996 | 129.166 | 129-125 |
| | sargla_1KPTI_zeamay_Phytozome | 105.277 | 99.0281 | 110.928 | MIN: 125 |
| | aratha_Phytozome_zeamay_Phytozome | 89.27 | 81.9304 | 96.1691 | MIN: 113 |
| | botbra_Phytozome_botter_1KPTI | 356.992 | 356.001 | 358 | 358-356 |
| | bryhyp_Hou_volcar_Phytozome | 978.819 | 945.56 | 1042.22 | 1056-948 |
| | codfra_1KPTI_bryhyp_Hou | 561.466 | 497.171 | 645.889 | MIN: 541 |
| | acrsp_1KPTI_plasp_1KPTI | 464.043 | 457.97 | 470.033 | 470-458 |
| LN | cyamer_Ensembl_chocri_Ensembl | 1159.85 | 994.913 | 1425 | MIN: 1047 |
| | graliv_1KPTI_chocri_Ensembl | 660.431 | 544.313 | 782.352 | MIN: 609 |
| | euaaff_1KPTI_zygsp_1KPTI | 602.242 | 501.799 | 721.559 | MIN: 350 |
| | marpol_Phytozome_zeamay_Phytozome | 483.562 | 461.767 | 515.668 | MIN:480 |
| | isoteg_1KPTI_zeamay_Phytozome | 421.665 | 419.23 | 423.107 | 423-419 |
| | ctesub_Qi_zeamay_Phytozome | 403.404 | 397.384 | 409.095 | MIN: 385 |
| | ginbil_1KPTI_zeamay_Phytozome | 324.231 | 322.677 | 328.012 | 330-323 |
| | ambtri_Phytozome_zeamay_Phytozome | 128.138 | 125.78 | 129.161 | 129-125 |
| | sargla_1KPTI_zeamay_Phytozome | 101.396 | 94.3616 | 107.919 | MIN: 125 |
| | aratha_Phytozome_zeamay_Phytozome | 83.3457 | 75.0767 | 91.1795 | MIN: 113 |
| | botbra_Phytozome_botter_1KPTI | 357.003 | 356.001 | 358.001 | 358-356 |
| | bryhyp_Hou_volcar_Phytozome | 981.857 | 945.324 | 1046.58 | 1056-948 |
| | codfra_1KPTI_bryhyp_Hou | 566.887 | 508.585 | 657.879 | MIN: 541 |
| | acrsp_1KPTI_plasp_1KPTI | 463.773 | 457.972 | 469.997 | 470-458 |
| UGAM | cyamer_Ensembl_chocri_Ensembl | 1237.17 | 1043.86 | 1490.83 | MIN: 1047 |
| | graliv_1KPTI_chocri_Ensembl | 648.391 | 531.747 | 804.893 | MIN: 609 |
| | euaaff_1KPTI_zygsp_1KPTI | 498.656 | 352.225 | 688.264 | MIN: 350 |
| | marpol_Phytozome_zeamay_Phytozome | 501.173 | 457.999 | 576.022 | MIN:480 |
| | isoteg_1KPTI_zeamay_Phytozome | 421.143 | 419.029 | 423.033 | 423-419 |
| | ctesub_Qi_zeamay_Phytozome | 391.489 | 375.648 | 405.254 | MIN: 385 |
| | ginbil_1KPTI_zeamay_Phytozome | 325.909 | 322.914 | 329.878 | 330-323 |
| | ambtri_Phytozome_zeamay_Phytozome | 128.259 | 126.068 | 129.181 | 129-125 |
| | sargla_1KPTI_zeamay_Phytozome | 113.509 | 107.896 | 118.308 | MIN: 125 |
| | aratha_Phytozome_zeamay_Phytozome | 97.4493 | 89.8008 | 104.404 | MIN: 113 |
| | botbra_Phytozome_botter_1KPTI | 356.989 | 355.995 | 357.996 | 358-356 |
| | bryhyp_Hou_volcar_Phytozome | 968.636 | 942.785 | 1027.57 | 1056-948 |
| | codfra_1KPTI_bryhyp_Hou | 564.364 | 478.222 | 683.923 | MIN: 541 |
| | acrsp_1KPTI_plasp_1KPTI | 463.609 | 457.973 | 469.92 | 470-458 |
| WN | cyamer_Ensembl_chocri_Ensembl | 1519.95 | 1305.91 | 1756.44 | MIN: 1047 |
| | graliv_1KPTI_chocri_Ensembl | 587.438 | 497.992 | 672.502 | MIN: 609 |

| | | | | | |
|---|---|---|---|---|---|
| | euaaff_1KPTI_zygsp_1KPTI | 571.745 | 471.764 | 675.812 | MIN: 350 |
| | marpol_Phytozome_zeamay_Phytozome | 523.979 | 467.307 | 596.594 | MIN:480 |
| | isoteg_1KPTI_zeamay_Phytozome | 421.132 | 419.031 | 423.013 | 423-419 |
| | ctesub_Qi_zeamay_Phytozome | 394.933 | 369.891 | 415.822 | MIN: 385 |
| | ginbil_1KPTI_zeamay_Phytozome | 326.106 | 322.952 | 329.939 | 330-323 |
| | ambtri_Phytozome_zeamay_Phytozome | 127.92 | 125.448 | 129.127 | 129-125 |
| | sargla_1KPTI_zeamay_Phytozome | 122.357 | 115.785 | 126.965 | MIN: 125 |
| | aratha_Phytozome_zeamay_Phytozome | 113.528 | 103.437 | 121.982 | MIN: 113 |
| | botbra_Phytozome_botter_1KPTI | 356.985 | 355.993 | 358 | 358-356 |
| | bryhyp_Hou_volcar_Phytozome | 976.837 | 944.744 | 1040.37 | 1056-948 |
| | codfra_1KPTI_bryhyp_Hou | 540.86 | 463.239 | 616.331 | MIN: 541 |
| | acrsp_1KPTI_plasp_1KPTI | 463.977 | 457.977 | 469.991 | 470-458 |
| 8 genes, ML | | | | | |
| CIR | cyamer_Ensembl_chocri_Ensembl | 1122.39 | 941.284 | 1348.46 | MIN: 1047 |
| | graliv_1KPTI_chocri_Ensembl | 631.749 | 559.069 | 715.531 | MIN: 609 |
| | euaaff_1KPTI_zygsp_1KPTI | 518.025 | 377.963 | 627.084 | MIN: 350 |
| | marpol_Phytozome_zeamay_Phytozome | 464.312 | 445.528 | 492.542 | MIN: 480 |
| | isoteg_1KPTI_zeamay_Phytozome | 421.596 | 419.169 | 423.082 | 423-419 |
| | ctesub_Qi_zeamay_Phytozome | 403.201 | 396.649 | 409.008 | MIN: 385 |
| | ginbil_1KPTI_zeamay_Phytozome | 324.42 | 322.699 | 328.467 | 330-323 |
| | ambtri_Phytozome_zeamay_Phytozome | 128.24 | 125.964 | 129.186 | 129-125 |
| | sargla_1KPTI_zeamay_Phytozome | 107.591 | 101.479 | 113.114 | MIN: 125 |
| | aratha_Phytozome_zeamay_Phytozome | 91.8245 | 84.4808 | 98.6075 | MIN: 113 |
| | botbra_Phytozome_botter_1KPTI | 356.977 | 355.994 | 357.999 | 358-356 |
| | ulvmut_orcAE_volcar_Phytozome | 969.589 | 943.564 | 1026.73 | 1056-948 |
| | acrsp_1KPTI_plasp_1KPTI | 464.237 | 458.033 | 470.004 | 470-458 |
| | codfra_1KPTI_bryhyp_Hou | 559.356 | 496.783 | 637.673 | MIN: 541 |
| LN | cyamer_Ensembl_chocri_Ensembl | 1161.31 | 1019.27 | 1402.89 | MIN: 1047 |
| | graliv_1KPTI_chocri_Ensembl | 666.323 | 574.054 | 774.607 | MIN: 609 |
| | euaaff_1KPTI_zygsp_1KPTI | 549.13 | 462.488 | 646.949 | MIN: 350 |
| | marpol_Phytozome_zeamay_Phytozome | 477.499 | 457.414 | 503.744 | MIN: 480 |
| | isoteg_1KPTI_zeamay_Phytozome | 421.647 | 419.208 | 423.086 | 423-419 |
| | ctesub_Qi_zeamay_Phytozome | 400.155 | 393.664 | 406.23 | MIN: 385 |
| | ginbil_1KPTI_zeamay_Phytozome | 324.208 | 322.638 | 327.81 | 330-323 |
| | ambtri_Phytozome_zeamay_Phytozome | 128.177 | 125.85 | 129.173 | 129-125 |
| | sargla_1KPTI_zeamay_Phytozome | 103.245 | 96.1049 | 109.789 | MIN: 125 |
| | aratha_Phytozome_zeamay_Phytozome | 85.0434 | 76.5844 | 92.9169 | MIN: 113 |
| | botbra_Phytozome_botter_1KPTI | 357.015 | 356.003 | 358.006 | 358-356 |
| | ulvmut_orcAE_volcar_Phytozome | 977.833 | 944.997 | 1041.4 | 1056-948 |
| | acrsp_1KPTI_plasp_1KPTI | 463.826 | 457.981 | 470.007 | 470-458 |
| | codfra_1KPTI_bryhyp_Hou | 553.43 | 496.047 | 617.988 | MIN: 541 |
| UGAM | cyamer_Ensembl_chocri_Ensembl | 1223.92 | 1035.5 | 1486 | MIN: 1047 |
| | graliv_1KPTI_chocri_Ensembl | 639.487 | 528.472 | 784.493 | MIN: 609 |
| | euaaff_1KPTI_zygsp_1KPTI | 482.34 | 351.657 | 646.127 | MIN: 350 |
| | marpol_Phytozome_zeamay_Phytozome | 494.817 | 453.403 | 565.535 | MIN: 480 |
| | isoteg_1KPTI_zeamay_Phytozome | 421.148 | 419.03 | 423.022 | 423-419 |
| | ctesub_Qi_zeamay_Phytozome | 389.396 | 373.153 | 403.378 | MIN: 385 |

| | | | | | |
|---|---|---|---|---|---|
| | ginbil_1KPTI_zeamay_Phytozome | 325.874 | 322.907 | 329.859 | 330-323 |
| | ambtri_Phytozome_zeamay_Phytozome | 128.264 | 126.121 | 129.181 | 129-125 |
| | sargla_1KPTI_zeamay_Phytozome | 114.437 | 108.796 | 119.146 | MIN: 125 |
| | aratha_Phytozome_zeamay_Phytozome | 98.7207 | 91.0944 | 105.484 | MIN: 113 |
| | botbra_Phytozome_botter_1KPTI | 356.995 | 356.002 | 358 | 358-356 |
| | ulvmut_orcAE_volcar_Phytozome | 967.637 | 943.327 | 1023.64 | 1056-948 |
| | acrsp_1KPTI_plasp_1KPTI | 463.713 | 457.962 | 469.973 | 470-458 |
| | codfra_1KPTI_bryhyp_Hou | 550.489 | 461.043 | 648.054 | MIN: 541 |
| WN | cyamer_Ensembl_chocri_Ensembl | #NAME? | 1356.83 | 1806.74 | MIN: 1047 |
| | graliv_1KPTI_chocri_Ensembl | #NAME? | 499.936 | 670.845 | MIN: 609 |
| | euaaff_1KPTI_zygsp_1KPTI | #NAME? | 459.326 | 659.03 | MIN: 350 |
| | marpol_Phytozome_zeamay_Phytozome | #NAME? | 465.431 | 590.764 | MIN: 480 |
| | isoteg_1KPTI_zeamay_Phytozome | #NAME? | 419.047 | 423.028 | 423-419 |
| | ctesub_Qi_zeamay_Phytozome | #NAME? | 368.8 | 415.57 | MIN: 385 |
| | ginbil_1KPTI_zeamay_Phytozome | #NAME? | 322.956 | 329.939 | 330-323 |
| | ambtri_Phytozome_zeamay_Phytozome | #NAME? | 125.473 | 129.131 | 129-125 |
| | sargla_1KPTI_zeamay_Phytozome | #NAME? | 115.614 | 126.883 | MIN: 125 |
| | aratha_Phytozome_zeamay_Phytozome | #NAME? | 103.856 | 122.01 | MIN: 113 |
| | botbra_Phytozome_botter_1KPTI | #NAME? | 356.004 | 357.995 | 358-356 |
| | ulvmut_orcAE_volcar_Phytozome | #NAME? | 944.617 | 1035.75 | 1056-948 |
| | acrsp_1KPTI_plasp_1KPTI | #NAME? | 458.015 | 470.037 | 470-458 |
| | codfra_1KPTI_bryhyp_Hou | #NAME? | 456.121 | 612.328 | MIN: 541 |
| 16 genes, CB | | | | | |
| CIR | cyamer_Ensembl_chocri_Ensembl | 1131.1 | 966.263 | 1375.31 | MIN: 1047 |
| | graliv_1KPTI_chocri_Ensembl | 640.866 | 560.305 | 735.014 | MIN: 609 |
| | euaaff_1KPTI_zygsp_1KPTI | 521.616 | 407.905 | 616.987 | MIN: 350 |
| | marpol_Phytozome_zeamay_Phytozome | 458.704 | 444.374 | 480.311 | MIN:480 |
| | isoteg_1KPTI_zeamay_Phytozome | 421.647 | 419.233 | 423.087 | 423-419 |
| | ctesub_Qi_zeamay_Phytozome | 403.552 | 397.989 | 408.458 | MIN: 385 |
| | ginbil_1KPTI_zeamay_Phytozome | 324.24 | 322.678 | 327.963 | 330-323 |
| | ambtri_Phytozome_zeamay_Phytozome | 128.245 | 126.092 | 129.189 | 129-125 |
| | sargla_1KPTI_zeamay_Phytozome | 100.116 | 93.9534 | 105.596 | MIN: 125 |
| | aratha_Phytozome_zeamay_Phytozome | 84.3725 | 77.2805 | 90.8907 | MIN: 113 |
| | botbra_Phytozome_botter_1KPTI | 356.99 | 355.999 | 358.003 | 358-356 |
| | bryhyp_Hou_volcar_Phytozome | 980.611 | 945.9 | 1045.02 | 1056-948 |
| | codfra_1KPTI_bryhyp_Hou | 554.187 | 493.531 | 627.173 | MIN: 541 |
| | acrsp_1KPTI_plasp_1KPTI | 464.16 | 458.004 | 470.059 | 470-458 |
| LN | cyamer_Ensembl_chocri_Ensembl | 1176.93 | 1008.32 | 1411.48 | MIN: 1047 |
| | graliv_1KPTI_chocri_Ensembl | 659.64 | 559.393 | 771.775 | MIN: 609 |
| | euaaff_1KPTI_zygsp_1KPTI | 560.485 | 482.758 | 671.63 | MIN: 350 |
| | marpol_Phytozome_zeamay_Phytozome | 470.869 | 453.53 | 494.824 | MIN:480 |
| | isoteg_1KPTI_zeamay_Phytozome | 421.71 | 419.26 | 423.098 | 423-419 |
| | ctesub_Qi_zeamay_Phytozome | 401.197 | 395.907 | 406.207 | MIN: 385 |
| | ginbil_1KPTI_zeamay_Phytozome | 324.144 | 322.643 | 327.713 | 330-323 |
| | ambtri_Phytozome_zeamay_Phytozome | 128.128 | 125.729 | 129.165 | 129-125 |
| | sargla_1KPTI_zeamay_Phytozome | 95.5552 | 88.7853 | 101.838 | MIN: 125 |
| | aratha_Phytozome_zeamay_Phytozome | 78.2807 | 70.4461 | 85.7146 | MIN: 113 |

| | | | | | |
|---|---|---|---|---|---|
| | botbra_Phytozome_botter_1KPTI | 357.003 | 355.999 | 358.001 | 358-356 |
| | bryhyp_Hou_volcar_Phytozome | 983.758 | 946.004 | 1047.57 | 1056-948 |
| | codfra_1KPTI_bryhyp_Hou | 548.356 | 498.086 | 613.119 | MIN: 541 |
| | acrsp_1KPTI_plasp_1KPTI | 463.876 | 458.009 | 469.973 | 470-458 |
| UGAM | cyamer_Ensembl_chocri_Ensembl | #NAME? | 1041.4 | 1473.41 | MIN: 1047 |
| | graliv_1KPTI_chocri_Ensembl | #NAME? | 527.95 | 799.73 | MIN: 609 |
| | euaaff_1KPTI_zygsp_1KPTI | #NAME? | 348.809 | 668.059 | MIN: 350 |
| | marpol_Phytozome_zeamay_Phytozome | #NAME? | 448.725 | 557.336 | MIN:480 |
| | isoteg_1KPTI_zeamay_Phytozome | #NAME? | 419.021 | 423.017 | 423-419 |
| | ctesub_Qi_zeamay_Phytozome | #NAME? | 372.862 | 404.408 | MIN: 385 |
| | ginbil_1KPTI_zeamay_Phytozome | #NAME? | 322.943 | 329.874 | 330-323 |
| | ambtri_Phytozome_zeamay_Phytozome | #NAME? | 126.181 | 129.177 | 129-125 |
| | sargla_1KPTI_zeamay_Phytozome | #NAME? | 105.412 | 115.837 | MIN: 125 |
| | aratha_Phytozome_zeamay_Phytozome | #NAME? | 88.305 | 102.222 | MIN: 113 |
| | botbra_Phytozome_botter_1KPTI | #NAME? | 356.003 | 358.006 | 358-356 |
| | bryhyp_Hou_volcar_Phytozome | #NAME? | 943.988 | 1030.58 | 1056-948 |
| | codfra_1KPTI_bryhyp_Hou | #NAME? | 463.393 | 657.022 | MIN: 541 |
| | acrsp_1KPTI_plasp_1KPTI | #NAME? | 457.922 | 469.903 | 470-458 |
| WN | cyamer_Ensembl_chocri_Ensembl | 1588.44 | 1381.46 | 1824.85 | MIN: 1047 |
| | graliv_1KPTI_chocri_Ensembl | 591.388 | 503.996 | 671.737 | MIN: 609 |
| | euaaff_1KPTI_zygsp_1KPTI | 561.427 | 464.967 | 654.049 | MIN: 350 |
| | marpol_Phytozome_zeamay_Phytozome | 507.775 | 458.099 | 573.907 | MIN:480 |
| | isoteg_1KPTI_zeamay_Phytozome | 421.131 | 419.024 | 423.026 | 423-419 |
| | ctesub_Qi_zeamay_Phytozome | 392.981 | 367.457 | 415.131 | MIN: 385 |
| | ginbil_1KPTI_zeamay_Phytozome | 326.059 | 322.938 | 329.895 | 330-323 |
| | ambtri_Phytozome_zeamay_Phytozome | 127.925 | 125.441 | 129.118 | 129-125 |
| | sargla_1KPTI_zeamay_Phytozome | 122.385 | 115.899 | 126.969 | MIN: 125 |
| | aratha_Phytozome_zeamay_Phytozome | 113.97 | 104.2 | 122.186 | MIN: 113 |
| | botbra_Phytozome_botter_1KPTI | 356.971 | 355.996 | 358 | 358-356 |
| | bryhyp_Hou_volcar_Phytozome | 989.835 | 946.781 | 1051.2 | 1056-948 |
| | codfra_1KPTI_bryhyp_Hou | 531.838 | 446.248 | 603.729 | MIN: 541 |
| | acrsp_1KPTI_plasp_1KPTI | 463.927 | 457.987 | 469.994 | 470-458 |
| 16 genes, ML | | | | | |
| CIR | cyamer_Ensembl_chocri_Ensembl | 1089.37 | 926.805 | 1331.98 | MIN: 1047 |
| | graliv_1KPTI_chocri_Ensembl | 638.132 | 564.623 | 725.578 | MIN: 609 |
| | euaaff_1KPTI_zygsp_1KPTI | 528.962 | 394.568 | 634.544 | MIN: 350 |
| | marpol_Phytozome_zeamay_Phytozome | 462.13 | 446.79 | 485.517 | MIN: 480 |
| | isoteg_1KPTI_zeamay_Phytozome | 421.659 | 419.2 | 423.097 | 423-419 |
| | ctesub_Qi_zeamay_Phytozome | 403.746 | 398.094 | 408.529 | MIN: 385 |
| | ginbil_1KPTI_zeamay_Phytozome | 324.261 | 322.683 | 328.086 | 330-323 |
| | ambtri_Phytozome_zeamay_Phytozome | 128.264 | 126.077 | 129.176 | 129-125 |
| | sargla_1KPTI_zeamay_Phytozome | 100.456 | 94.4046 | 105.953 | MIN: 125 |
| | aratha_Phytozome_zeamay_Phytozome | 85.0202 | 78.2047 | 91.5794 | MIN: 113 |
| | botbra_Phytozome_botter_1KPTI | 356.996 | 356 | 358.002 | 358-356 |
| | ulvmut_orcAE_volcar_Phytozome | 979.384 | 945.431 | 1042.6 | 1056-948 |
| | acrsp_1KPTI_plasp_1KPTI | 464.129 | 458.029 | 469.997 | 470-458 |
| | codfra_1KPTI_bryhyp_Hou | 543.778 | 483.383 | 610.362 | MIN: 541 |

| | | | | | |
|---|---|---|---|---|---|
| LN | cyamer_Ensembl_chocri_Ensembl | 1132.71 | 975.461 | 1342.13 | MIN: 1047 |
| | graliv_1KPTI_chocri_Ensembl | 658.921 | 565.28 | 753.681 | MIN: 609 |
| | euaaff_1KPTI_zygsp_1KPTI | 564.817 | 484.914 | 679.523 | MIN: 350 |
| | marpol_Phytozome_zeamay_Phytozome | 474.18 | 456.795 | 497.951 | MIN: 480 |
| | isoteg_1KPTI_zeamay_Phytozome | 421.717 | 419.253 | 423.099 | 423-419 |
| | ctesub_Qi_zeamay_Phytozome | 401.124 | 395.906 | 406.074 | MIN: 385 |
| | ginbil_1KPTI_zeamay_Phytozome | 324.168 | 322.631 | 327.833 | 330-323 |
| | ambtri_Phytozome_zeamay_Phytozome | 128.146 | 125.79 | 129.17 | 129-125 |
| | sargla_1KPTI_zeamay_Phytozome | 95.2551 | 88.2655 | 101.596 | MIN: 125 |
| | aratha_Phytozome_zeamay_Phytozome | 78.1337 | 69.989 | 85.6711 | MIN: 113 |
| | botbra_Phytozome_botter_1KPTI | 356.997 | 356.002 | 357.999 | 358-356 |
| | ulvmut_orcAE_volcar_Phytozome | 987.507 | 946.663 | 1051.59 | 1056-948 |
| | acrsp_1KPTI_plasp_1KPTI | 463.893 | 457.984 | 469.933 | 470-458 |
| | codfra_1KPTI_bryhyp_Hou | 541.256 | 484.161 | 594.573 | MIN: 541 |
| UGAM | cyamer_Ensembl_chocri_Ensembl | 1216.33 | 1039.04 | 1454.19 | MIN: 1047 |
| | graliv_1KPTI_chocri_Ensembl | 644.591 | 535.02 | 795.853 | MIN: 609 |
| | euaaff_1KPTI_zygsp_1KPTI | 493.738 | 352.502 | 670.363 | MIN: 350 |
| | marpol_Phytozome_zeamay_Phytozome | 493.29 | 451.394 | 566.974 | MIN: 480 |
| | isoteg_1KPTI_zeamay_Phytozome | 421.104 | 419.01 | 423.011 | 423-419 |
| | ctesub_Qi_zeamay_Phytozome | 389.661 | 372.755 | 404.186 | MIN: 385 |
| | ginbil_1KPTI_zeamay_Phytozome | 325.966 | 322.914 | 329.886 | 330-323 |
| | ambtri_Phytozome_zeamay_Phytozome | 128.303 | 126.191 | 129.205 | 129-125 |
| | sargla_1KPTI_zeamay_Phytozome | 111.5 | 106.023 | 116.242 | MIN: 125 |
| | aratha_Phytozome_zeamay_Phytozome | 96.14 | 88.873 | 102.674 | MIN: 113 |
| | botbra_Phytozome_botter_1KPTI | 356.993 | 355.993 | 357.998 | 358-356 |
| | ulvmut_orcAE_volcar_Phytozome | 969.125 | 943.643 | 1027.45 | 1056-948 |
| | acrsp_1KPTI_plasp_1KPTI | 463.725 | 457.968 | 469.959 | 470-458 |
| | codfra_1KPTI_bryhyp_Hou | 541.841 | 448.686 | 631.963 | MIN: 541 |
| WN | cyamer_Ensembl_chocri_Ensembl | 1519.33 | 1316.87 | 1740.33 | MIN: 1047 |
| | graliv_1KPTI_chocri_Ensembl | 594.154 | 508.986 | 673.558 | MIN: 609 |
| | euaaff_1KPTI_zygsp_1KPTI | 572.617 | 478.274 | 664.017 | MIN: 350 |
| | marpol_Phytozome_zeamay_Phytozome | 515.665 | 463.195 | 583.763 | MIN: 480 |
| | isoteg_1KPTI_zeamay_Phytozome | 421.141 | 419.021 | 423.021 | 423-419 |
| | ctesub_Qi_zeamay_Phytozome | 393.051 | 367.433 | 415.226 | MIN: 385 |
| | ginbil_1KPTI_zeamay_Phytozome | 326.119 | 322.946 | 329.947 | 330-323 |
| | ambtri_Phytozome_zeamay_Phytozome | 127.937 | 125.464 | 129.134 | 129-125 |
| | sargla_1KPTI_zeamay_Phytozome | 122.427 | 115.997 | 126.904 | MIN: 125 |
| | aratha_Phytozome_zeamay_Phytozome | 114.027 | 104.275 | 122.02 | MIN: 113 |
| | botbra_Phytozome_botter_1KPTI | 356.975 | 355.999 | 357.997 | 358-356 |
| | ulvmut_orcAE_volcar_Phytozome | 985.031 | 946.161 | 1047.2 | 1056-948 |
| | acrsp_1KPTI_plasp_1KPTI | 463.996 | 458.005 | 469.996 | 470-458 |
| | codfra_1KPTI_bryhyp_Hou | 516.452 | 422.254 | 589.645 | MIN: 541 |

Normal run - Removed fossils

| | Node | Mean date | Lower 95% | Upper 95% | Fossil age c |
|---|---|---|---|---|---|
| 8 genes, CB | | | | | |
| CIR | caulen_OIST_caucyl_Unlu | 413.881 | 308.162 | 504.154 | MIN: 505 |

| | | | | | |
|---|---|---|---|---|---|
| | oedcar_1KPTI_oedfov_1KPTI | 385.033 | 381.654 | 392.024 | 393-382 |
| | stihel_1KPTI_aphrep_1KPTI | 108.34 | 103.081 | 110.649 | 110-97 |
| LN | caulen_OIST_caucyl_Unlu | 491.272 | 385.452 | 558.602 | MIN: 505 |
| | oedcar_1KPTI_oedfov_1KPTI | 386.737 | 381.882 | 392.93 | 393-382 |
| | stihel_1KPTI_aphrep_1KPTI | 106.8 | 98.5285 | 110.57 | 110-97 |
| UGAM | caulen_OIST_caucyl_Unlu | 515.11 | 417.461 | 616.733 | MIN: 505 |
| | oedcar_1KPTI_oedfov_1KPTI | 387.195 | 381.974 | 392.948 | 393-382 |
| | stihel_1KPTI_aphrep_1KPTI | 103.702 | 97.0804 | 110.08 | 110-97 |
| WN | caulen_OIST_caucyl_Unlu | 408.765 | 291.14 | 511.338 | MIN: 505 |
| | oedcar_1KPTI_oedfov_1KPTI | 386.445 | 381.798 | 392.692 | 393-382 |
| | stihel_1KPTI_aphrep_1KPTI | 104.059 | 97.0082 | 110.209 | 110-97 |
| 8 genes, ML | | | | | |
| CIR | caulen_OIST_caucyl_Unlu | 396.869 | 285.636 | 494.854 | MIN: 505 |
| | oedcar_1KPTI_oedfov_1KPTI | 385.104 | 381.628 | 392.145 | 393-382 |
| | stihel_1KPTI_aphrep_1KPTI | 108.35 | 103.242 | 110.721 | 110-97 |
| LN | caulen_OIST_caucyl_Unlu | 474.614 | 358.82 | 552.137 | MIN: 505 |
| | oedcar_1KPTI_oedfov_1KPTI | 386.483 | 381.894 | 392.825 | 393-382 |
| | stihel_1KPTI_aphrep_1KPTI | 106.482 | 97.7906 | 110.605 | 110-97 |
| UGAM | caulen_OIST_caucyl_Unlu | 509.189 | 405.478 | 606.142 | MIN: 505 |
| | oedcar_1KPTI_oedfov_1KPTI | 387.037 | 381.898 | 392.979 | 393-382 |
| | stihel_1KPTI_aphrep_1KPTI | 104.066 | 97.1068 | 110.18 | 110-97 |
| WN | caulen_OIST_caucyl_Unlu | 399.055 | 283.965 | 508.146 | MIN: 505 |
| | oedcar_1KPTI_oedfov_1KPTI | 386.383 | 381.899 | 392.867 | 393-382 |
| | stihel_1KPTI_aphrep_1KPTI | 104.222 | 97.1346 | 110.144 | 110-97 |
| 16 genes, CB | | | | | |
| CIR | caulen_OIST_caucyl_Unlu | 375.665 | 260.471 | 461.974 | MIN: 505 |
| | oedcar_1KPTI_oedfov_1KPTI | 384.894 | 381.678 | 391.754 | 393-382 |
| | stihel_1KPTI_aphrep_1KPTI | 109.03 | 105.665 | 110.978 | 110-97 |
| LN | caulen_OIST_caucyl_Unlu | 479.946 | 359.26 | 550.935 | MIN: 505 |
| | oedcar_1KPTI_oedfov_1KPTI | 386.689 | 381.959 | 392.919 | 393-382 |
| | stihel_1KPTI_aphrep_1KPTI | 107.229 | 99.3214 | 110.53 | 110-97 |
| UGAM | caulen_OIST_caucyl_Unlu | 514.466 | 418.231 | 605.673 | MIN: 505 |
| | oedcar_1KPTI_oedfov_1KPTI | 387.147 | 382.004 | 392.962 | 393-382 |
| | stihel_1KPTI_aphrep_1KPTI | 104.09 | 97.2736 | 110.17 | 110-97 |
| WN | caulen_OIST_caucyl_Unlu | 373.529 | 272.235 | 485.025 | MIN: 505 |
| | oedcar_1KPTI_oedfov_1KPTI | 386.374 | 381.862 | 392.721 | 393-382 |
| | stihel_1KPTI_aphrep_1KPTI | 104.44 | 97.4831 | 110.114 | 110-97 |
| 16 genes, ML | | | | | |
| CIR | caulen_OIST_caucyl_Unlu | 351.177 | 250.985 | 443.95 | MIN: 505 |
| | oedcar_1KPTI_oedfov_1KPTI | 384.866 | 381.449 | 391.807 | 393-382 |
| | stihel_1KPTI_aphrep_1KPTI | 109.065 | 105.314 | 111.08 | 110-97 |
| LN | caulen_OIST_caucyl_Unlu | 466.975 | 346.922 | 547.145 | MIN: 505 |
| | oedcar_1KPTI_oedfov_1KPTI | 386.772 | 381.898 | 392.851 | 393-382 |
| | stihel_1KPTI_aphrep_1KPTI | 107.443 | 99.9763 | 110.577 | 110-97 |
| UGAM | caulen_OIST_caucyl_Unlu | 507.449 | 403.101 | 597.306 | MIN: 505 |
| | oedcar_1KPTI_oedfov_1KPTI | 387.269 | 381.987 | 393.057 | 393-382 |
| | stihel_1KPTI_aphrep_1KPTI | 103.958 | 96.9547 | 110.266 | 110-97 |

| | | | | | |
|---|---|---|---|---|---|
| WN | caulen_OIST_caucyl_Unlu | 354.671 | 257.32 | 474.437 | MIN: 505 |
| | oedcar_1KPTI_oedfov_1KPTI | 386.31 | 381.678 | 392.697 | 393-382 |
| | stihel_1KPTI_aphrep_1KPTI | 104.45 | 97.1732 | 110.187 | 110-97 |

Fossil cross validation run

| Node | Mean date | Lower 95% | Upper 95% HPDI |
|---|---|---|---|
| cyamer_En | 952.754 | 877.105 | 1058.41 |
| graliv_1KP | 566.615 | 361.991 | 706.595 |
| euaaff_1KF | 537.815 | 397.252 | 667.063 |
| marpol_Ph | 462.543 | 447.377 | 486.529 |
| isoteg_1KP | 620.792 | 544.026 | 704.688 |
| ctesub_Qi_ | 406.549 | 400.621 | 412.06 |
| ginbil_1KP | 202.955 | 184.757 | 225.667 |
| ambtri_Phy | 260.388 | 244.092 | 275.015 |
| sargla_1KP | 103.385 | 96.4085 | 109.64 |
| aratha_Phy | 86.1042 | 78.6394 | 93.5776 |
| botbra_Phy | 300.992 | 205.501 | 392.715 |
| bryhyp_Ho | 969.85 | 944.037 | 1029.52 |
| codfra_1KF | 460.462 | 318.694 | 627.277 |
| acrsp_1KP | 402.202 | 254.784 | 611.382 |
| cyamer_En | 991.513 | 858.215 | 1157.61 |
| graliv_1KP | 518.605 | 340.987 | 720.116 |
| euaaff_1KF | 600.235 | 474.043 | 723.097 |
| marpol_Ph | 477.555 | 457.249 | 507.995 |
| isoteg_1KP | 613.705 | 552.097 | 684.982 |
| ctesub_Qi_ | 403.803 | 397.532 | 409.281 |
| ginbil_1KP | 203.845 | 185.967 | 224.696 |
| ambtri_Phy | 252.535 | 232.63 | 269.552 |
| sargla_1KP | 98.4792 | 90.5993 | 105.985 |
| aratha_Phy | 78.1526 | 68.8216 | 87.3476 |
| botbra_Phy | 411.692 | 250.787 | 547.675 |
| bryhyp_Ho | 977.473 | 945.168 | 1041.7 |
| codfra_1KF | 511.633 | 334.441 | 642.898 |
| acrsp_1KP | 390.511 | 279.518 | 533.471 |
| cyamer_En | 993.897 | 849.154 | 1164.64 |
| graliv_1KP | 322.361 | 202.647 | 514.77 |
| euaaff_1KF | 465.749 | 297.654 | 650.916 |
| marpol_Ph | 487.069 | 449.414 | 560.295 |
| isoteg_1KP | 482.641 | 422.788 | 572.841 |
| ctesub_Qi_ | 391.19 | 372.052 | 405.675 |
| ginbil_1KP | 231.066 | 181.507 | 296.479 |
| ambtri_Phy | 265.96 | 244.209 | 285.055 |
| sargla_1KP | 111.607 | 104.658 | 117.129 |
| aratha_Phy | 93.9419 | 84.8295 | 102.225 |
| botbra_Phy | 175.941 | 85.5151 | 360.975 |
| bryhyp_Ho | 965.219 | 942.588 | 1016.44 |
| codfra_1KF | 315.054 | 179.061 | 510.523 |
| acrsp_1KP | 326.935 | 230.326 | 431.51 |
| cyamer_En | 1515.57 | 1150.07 | 1779.78 |
| graliv_1KP | 428.532 | 316.374 | 559.394 |

| | | | |
|---|---|---|---|
| euaaff_1KF | 560.897 | 466.606 | 663.797 |
| marpol_Ph | 518.69 | 449.306 | 595.411 |
| isoteg_1KP | 571.63 | 479.609 | 662.558 |
| ctesub_Qi_ | 392.301 | 354.173 | 415.823 |
| ginbil_1KPT | 261.624 | 199.439 | 327.518 |
| ambtri_Phy | 293.587 | 259.472 | 320.594 |
| sargla_1KP | 120.269 | 112.428 | 125.979 |
| aratha_Phy | 109.566 | 92.6081 | 121.597 |
| botbra_Phy | 272.455 | 185.99 | 369.104 |
| bryhyp_Ho | 972.551 | 944.721 | 1032.43 |
| codfra_1KF | 425.322 | 313.673 | 573.426 |
| acrsp_1KPT | 456.047 | 357.675 | 564.529 |
| cyamer_En | 968.561 | 821.927 | 1167.21 |
| graliv_1KPT | 530.194 | 347.899 | 688.558 |
| euaaff_1KF | 494.368 | 371.262 | 616.002 |
| marpol_Ph | 458.16 | 443.314 | 480.746 |
| isoteg_1KP | 612.538 | 542.169 | 691.643 |
| ctesub_Qi_ | 403.55 | 397.461 | 408.699 |
| ginbil_1KPT | 201.75 | 182.417 | 227.494 |
| ambtri_Phy | 260.823 | 243.999 | 274.85 |
| sargla_1KP | 105.88 | 99.501 | 111.833 |
| aratha_Phy | 88.3127 | 80.6419 | 96.0216 |
| botbra_Phy | 288.795 | 184.5 | 376.223 |
| ulvmut_orc | 860.079 | 761.919 | 965.852 |
| acrsp_1KPT | 421.882 | 262.062 | 609.196 |
| codfra_1KF | 466.799 | 312 | 600.452 |
| cyamer_En | 1034.47 | 910.463 | 1210.77 |
| graliv_1KPT | 496.547 | 298.407 | 682.236 |
| euaaff_1KF | 534.353 | 434.11 | 648.129 |
| marpol_Ph | 472.164 | 453.312 | 499.375 |
| isoteg_1KP | 634.638 | 562.2 | 715.253 |
| ctesub_Qi_ | 400.545 | 393.835 | 406.672 |
| ginbil_1KPT | 203.234 | 185.625 | 226.447 |
| ambtri_Phy | 252.241 | 232.518 | 269.816 |
| sargla_1KP | 100.727 | 93.2888 | 107.565 |
| aratha_Phy | 80.0408 | 71.172 | 88.6051 |
| botbra_Phy | 383.526 | 222.225 | 528.343 |
| ulvmut_orc | 883.338 | 789.212 | 998.671 |
| acrsp_1KPT | 397.606 | 264.9 | 544.128 |
| codfra_1KF | 474.178 | 316.368 | 599.976 |
| cyamer_En | 1290.31 | 945.61 | 1811.38 |
| graliv_1KPT | 327.096 | 212.33 | 505.245 |
| euaaff_1KF | 460.57 | 311.79 | 620.859 |
| marpol_Ph | 479.162 | 445.69 | 545.38 |
| isoteg_1KP | 487.132 | 427.998 | 586.139 |
| ctesub_Qi_ | 388.356 | 370.819 | 403.502 |

| | | | |
|---|---|---|---|
| ginbil_1KP | 229.798 | 180.824 | 300.184 |
| ambtri_Phy | 266.324 | 244.271 | 284.618 |
| sargla_1KP | 112.333 | 106.17 | 117.644 |
| aratha_Phy | 95.1662 | 85.6905 | 103.088 |
| botbra_Phy | 169.229 | 86.4677 | 350.505 |
| ulvmut_orc | 802.362 | 709.158 | 912.656 |
| acrsp_1KP | 345.931 | 246.14 | 494.284 |
| codfra_1KF | 307.777 | 186.775 | 487.737 |
| cyamer_En | 1616.28 | 1430.7 | 1827.68 |
| graliv_1KP | 436.117 | 305.081 | 574.632 |
| euaaff_1KF | 547.87 | 455.133 | 648.247 |
| marpol_Ph | 513.735 | 448.357 | 586.479 |
| isoteg_1KP | 581.76 | 498.439 | 671.945 |
| ctesub_Qi_ | 391.058 | 351.373 | 416.3 |
| ginbil_1KP | 264.947 | 204.929 | 328.446 |
| ambtri_Phy | 290.274 | 256.183 | 319.669 |
| sargla_1KP | 120.387 | 112.312 | 126.174 |
| aratha_Phy | 110.849 | 94.5254 | 122.095 |
| botbra_Phy | 264.473 | 171.869 | 358.723 |
| ulvmut_orc | 889.293 | 787.758 | 1008.51 |
| acrsp_1KP | 480.526 | 375.136 | 583.346 |
| codfra_1KF | 415.719 | 309.274 | 553.57 |
| cyamer_En | 1035.8 | 892.92 | 1195.25 |
| graliv_1KP | 583.95 | 414.536 | 729.13 |
| euaaff_1KF | 495.731 | 394.098 | 617.186 |
| marpol_Ph | 455.269 | 443.038 | 472.375 |
| isoteg_1KP | 589.969 | 512.004 | 676.002 |
| ctesub_Qi_ | 403.63 | 398.01 | 408.392 |
| ginbil_1KP | 210.987 | 192.811 | 233.983 |
| ambtri_Phy | 251.26 | 235.386 | 265.304 |
| sargla_1KP | 98.4234 | 91.3139 | 104.301 |
| aratha_Phy | 81.0201 | 74.2431 | 87.4056 |
| botbra_Phy | 293.286 | 184.096 | 385.085 |
| bryhyp_Ho | 978.883 | 945.8 | 1041.59 |
| codfra_1KF | 477.885 | 322.195 | 613.754 |
| acrsp_1KP | 363.605 | 225.027 | 569.356 |
| cyamer_En | 1042.6 | 877.893 | 1205.74 |
| graliv_1KP | 481.638 | 303.236 | 635.196 |
| euaaff_1KF | 554.26 | 460.557 | 672.654 |
| marpol_Ph | 465.549 | 451.208 | 486.382 |
| isoteg_1KP | 622.451 | 556.118 | 721.449 |
| ctesub_Qi_ | 401.261 | 396.141 | 406.045 |
| ginbil_1KP | 211.666 | 195.375 | 228.759 |
| ambtri_Phy | 244.014 | 227.685 | 258.094 |
| sargla_1KP | 92.7495 | 84.6288 | 99.5887 |
| aratha_Phy | 73.2701 | 64.6175 | 81.633 |

| | | | |
|---|---|---|---|
| botbra_Phy | 393.728 | 232.51 | 534.491 |
| bryhyp_Ho | 985.764 | 945.928 | 1051.22 |
| codfra_1KF | 461.695 | 307.467 | 581.744 |
| acrsp_1KPT | 358.069 | 242.745 | 482.933 |
| cyamer_En | 1307.57 | 950.061 | 1645.5 |
| graliv_1KP | 309.405 | 200.57 | 501.187 |
| euaaff_1KF | 451.152 | 292.657 | 605.6 |
| marpol_Ph | 473.817 | 441.796 | 537.848 |
| isoteg_1KP | 469.535 | 417.555 | 551.08 |
| ctesub_Qi_ | 388.255 | 369.803 | 404.165 |
| ginbil_1KPT | 228.141 | 178.74 | 294.57 |
| ambtri_Phy | 267.643 | 248.531 | 285.729 |
| sargla_1KP | 109.344 | 102.712 | 114.394 |
| aratha_Phy | 91.9391 | 83.3564 | 100.156 |
| botbra_Phy | 145.229 | 76.1317 | 304.625 |
| bryhyp_Ho | 966.633 | 943.007 | 1019.81 |
| codfra_1KF | 288.263 | 161.113 | 461.219 |
| acrsp_1KPT | 322.159 | 215.648 | 449.557 |
| cyamer_En | 1676.71 | 1454.21 | 1896.68 |
| graliv_1KP | 452.862 | 332.72 | 587.865 |
| euaaff_1KF | 549.885 | 459.055 | 642.331 |
| marpol_Ph | 497.46 | 441.615 | 575.456 |
| isoteg_1KP | 531.15 | 452.585 | 612.455 |
| ctesub_Qi_ | 389.013 | 351.055 | 415.04 |
| ginbil_1KPT | 260.289 | 199.377 | 323.288 |
| ambtri_Phy | 290.577 | 252.734 | 318.264 |
| sargla_1KP | 120.368 | 113.207 | 126.227 |
| aratha_Phy | 110.782 | 93.7865 | 122.961 |
| botbra_Phy | 253.209 | 159.979 | 342.878 |
| bryhyp_Ho | 984.682 | 945.908 | 1050.65 |
| codfra_1KF | 393.077 | 289.398 | 508.717 |
| acrsp_1KPT | 452.667 | 346.877 | 567.009 |
| cyamer_En | 989.912 | 842.626 | 1155.24 |
| graliv_1KP | 573.385 | 424.342 | 715.036 |
| euaaff_1KF | 522.403 | 388.753 | 639.001 |
| marpol_Ph | 457.201 | 443.505 | 475.595 |
| isoteg_1KP | 590.487 | 524.401 | 664.066 |
| ctesub_Qi_ | 403.695 | 398.007 | 408.752 |
| ginbil_1KPT | 204.302 | 186.266 | 226.654 |
| ambtri_Phy | 254.984 | 239.868 | 270.003 |
| sargla_1KP | 98.3489 | 91.5966 | 104.651 |
| aratha_Phy | 81.6565 | 73.7152 | 89.495 |
| botbra_Phy | 298.188 | 180.292 | 390.907 |
| ulvmut_orc | 859.319 | 776.244 | 927.312 |
| acrsp_1KPT | 406.872 | 260.033 | 593.036 |
| codfra_1KF | 441.682 | 317.82 | 571.291 |

| | | | |
|---|---|---|---|
| cyamer_En | 971.816 | 837.749 | 1099.43 |
| graliv_1KP | 503.586 | 335.449 | 669.27 |
| euaaff_1KF | 558.352 | 458.83 | 669.394 |
| marpol_Ph | 468.401 | 453.504 | 491.809 |
| isoteg_1KP | 625.495 | 566.528 | 703.812 |
| ctesub_Qi_ | 401.115 | 395.693 | 406.085 |
| ginbil_1KP | 205.272 | 188.546 | 224.597 |
| ambtri_Phy | 248.362 | 230.477 | 264.355 |
| sargla_1KP | 92.7671 | 85.5522 | 99.4067 |
| aratha_Phy | 72.9788 | 64.4727 | 81.8903 |
| botbra_Phy | 388.573 | 217.667 | 528.077 |
| ulvmut_orc | 902.349 | 797.241 | 998.431 |
| acrsp_1KPT | 370.745 | 244.189 | 531.717 |
| codfra_1KF | 408.864 | 257.098 | 549.319 |
| cyamer_En | 1378.52 | 986.724 | 1653.23 |
| graliv_1KP | 323.765 | 212.879 | 490.754 |
| euaaff_1KF | 473.07 | 318.36 | 626.502 |
| marpol_Ph | 475.967 | 445.515 | 539.582 |
| isoteg_1KP | 473.74 | 416.111 | 563.914 |
| ctesub_Qi_ | 388.554 | 368.138 | 404.518 |
| ginbil_1KP | 226.707 | 175.341 | 297.569 |
| ambtri_Phy | 270.904 | 251.517 | 287.466 |
| sargla_1KP | 109.806 | 103.557 | 115.465 |
| aratha_Phy | 92.7664 | 83.6386 | 100.453 |
| botbra_Phy | 155.735 | 77.0947 | 311.346 |
| ulvmut_orc | 813.06 | 716.458 | 933.745 |
| acrsp_1KPT | 326.084 | 220.526 | 468.915 |
| codfra_1KF | 267.527 | 152.147 | 443.702 |
| cyamer_En | 1595.8 | 1392.45 | 1831.59 |
| graliv_1KP | 462.711 | 332.36 | 587.992 |
| euaaff_1KF | 561.812 | 465.023 | 663.093 |
| marpol_Ph | 510.779 | 449.871 | 591.586 |
| isoteg_1KP | 545.804 | 459.988 | 630.03 |
| ctesub_Qi_ | 389.817 | 350.065 | 415.948 |
| ginbil_1KP | 258.834 | 195.142 | 317.925 |
| ambtri_Phy | 294.655 | 260.913 | 321.338 |
| sargla_1KP | 120.337 | 112.596 | 125.855 |
| aratha_Phy | 110.833 | 93.3744 | 121.757 |
| botbra_Phy | 259.756 | 161.679 | 347.313 |
| ulvmut_orc | 940.791 | 840.629 | 1054.37 |
| acrsp_1KPT | 471.61 | 367.18 | 584.546 |
| codfra_1KF | 371.292 | 282.308 | 489.936 |

il cross validation run - Removed fc

| | Mean date | Lower 95% | Upper 95% HPDI |
|---|---|---|---|
| caulen_OIS | 198.698 | 113.535 | 327.891 |

| | | | |
|---|---|---|---|
| oedcar_1KI | 66.6015 | 41.4781 | 103.728 |
| stihel_1KP | 498.913 | 378.012 | 611.5 |
| caulen_OIS | 239.259 | 142.897 | 384.288 |
| oedcar_1KI | 74.3321 | 44.7977 | 114.596 |
| stihel_1KP | 568.632 | 471.829 | 637.825 |
| caulen_OIS | 161.687 | 94.4988 | 308.344 |
| oedcar_1KI | 76.138 | 38.1205 | 127.213 |
| caulen_OIS | 225.594 | 124.611 | 328.976 |
| oedcar_1KI | 104.102 | 48.6765 | 163.416 |
| stihel_1KP | 327.781 | 214.543 | 429.635 |
| caulen_OIS | 209.669 | 113.736 | 336.908 |
| oedcar_1KI | 71.5391 | 42.6155 | 108.683 |
| stihel_1KP | 493.622 | 382.817 | 598.593 |
| caulen_OIS | 251.32 | 156.537 | 364.929 |
| oedcar_1KI | 81.9049 | 47.9545 | 130.953 |
| stihel_1KP | 575.897 | 473.179 | 645.887 |
| caulen_OIS | 162.786 | 97.114 | 298.064 |
| oedcar_1KI | 80.1214 | 41.3946 | 142.513 |
| stihel_1KP | 235.176 | 97.5333 | 466.721 |
| caulen_OIS | 222.868 | 126.284 | 326.133 |
| oedcar_1KI | 109.803 | 52.3767 | 173.521 |
| stihel_1KP | 338.573 | 229.812 | 451.493 |
| caulen_OIS | 198.044 | 113.029 | 315.26 |
| oedcar_1KI | 57.4257 | 36.1327 | 89.4941 |
| stihel_1KP | 592.925 | 463.292 | 681.582 |
| caulen_OIS | 235.615 | 148.737 | 349.956 |
| oedcar_1KI | 68.6022 | 41.6347 | 110.18 |
| stihel_1KP | 603.341 | 527.012 | 669.248 |
| caulen_OIS | 137.436 | 80.3688 | 273.924 |
| oedcar_1KI | 69.8496 | 33.043 | 122.83 |
| stihel_1KP | 261.747 | 104.58 | 467.918 |
| caulen_OIS | 198.56 | 110.714 | 301.72 |
| oedcar_1KI | 104.334 | 47.8335 | 169.647 |
| stihel_1KP | 364.232 | 258.511 | 479.844 |
| caulen_OIS | 200.863 | 117.999 | 300.811 |
| oedcar_1KI | 58.9204 | 37.6046 | 89.453 |
| stihel_1KP | 613.969 | 505.909 | 701.03 |
| caulen_OIS | 251.32 | 156.537 | 364.929 |
| oedcar_1KI | 81.9049 | 47.9545 | 130.953 |
| stihel_1KP | 575.897 | 473.179 | 645.887 |
| caulen_OIS | 162.786 | 97.114 | 298.064 |
| oedcar_1KI | 80.1214 | 41.3946 | 142.513 |
| stihel_1KP | 235.176 | 97.5333 | 466.721 |

| | | | |
|---|---|---|---|
| caulen_OIS | 222.868 | 126.284 | 326.133 |
| oedcar_1KI | 109.803 | 52.3767 | 173.521 |
| stihel_1KP | 338.573 | 229.812 | 451.493 |
