## Supplementary figures and images for "Fossil-calibrated inference of divergence times among the Volvocine algae enables reconstruction of the steps that led to differentiated multicellularity"

### asymmetric_division_tree.plot.pdf

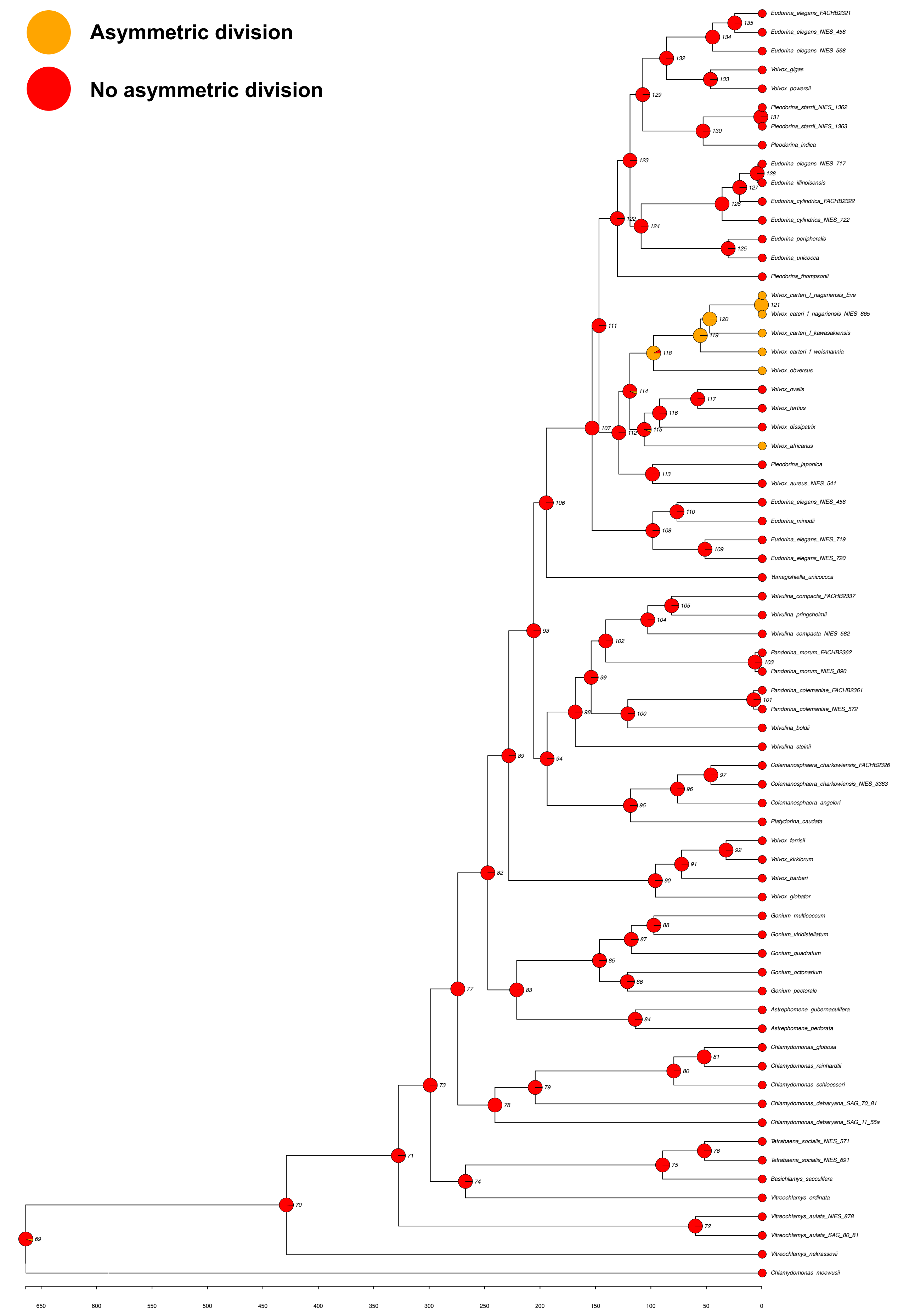

### bbr_tree.plot.pdf

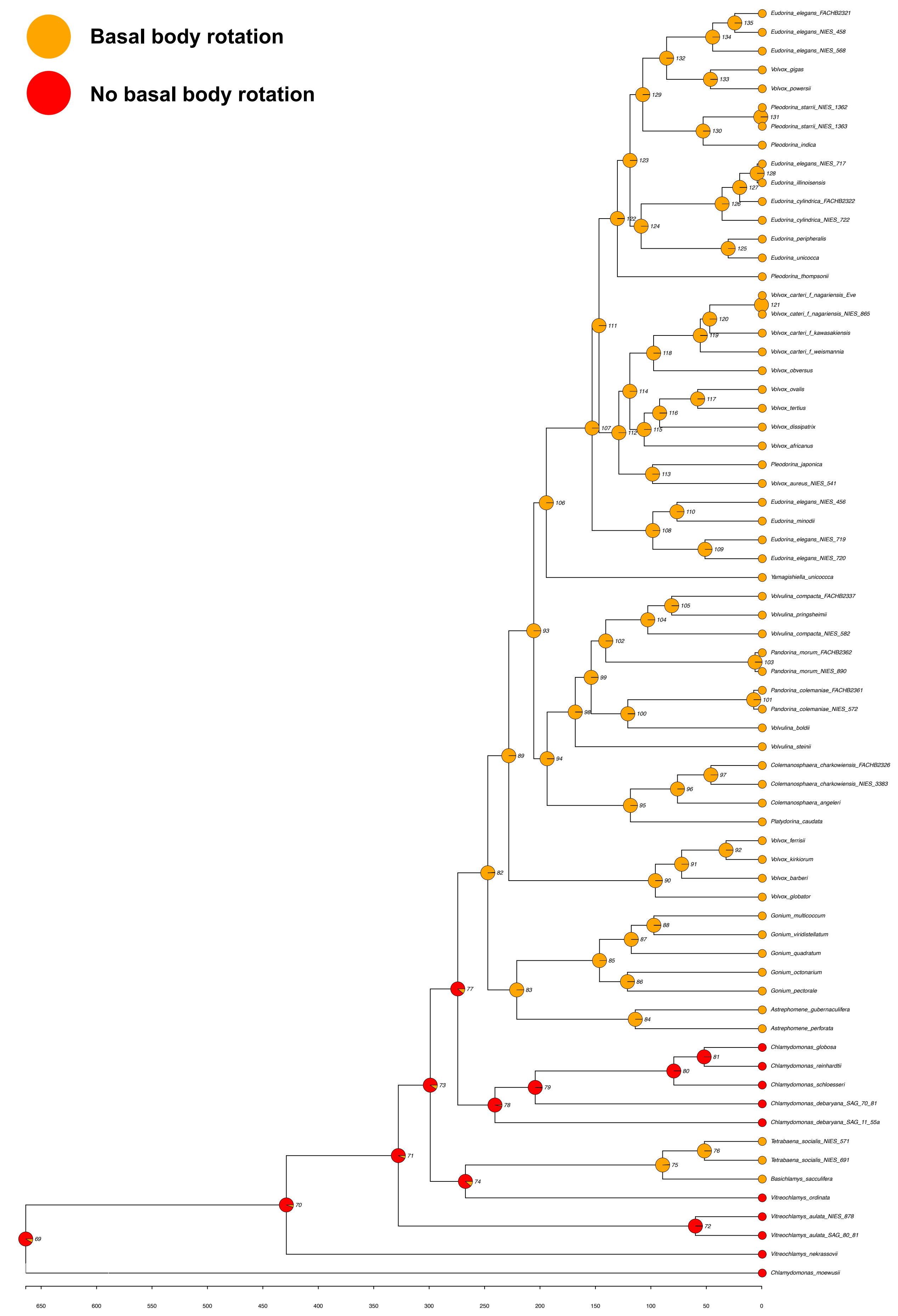

### bifurcated_cell_div_prog_tree.plot.pdf

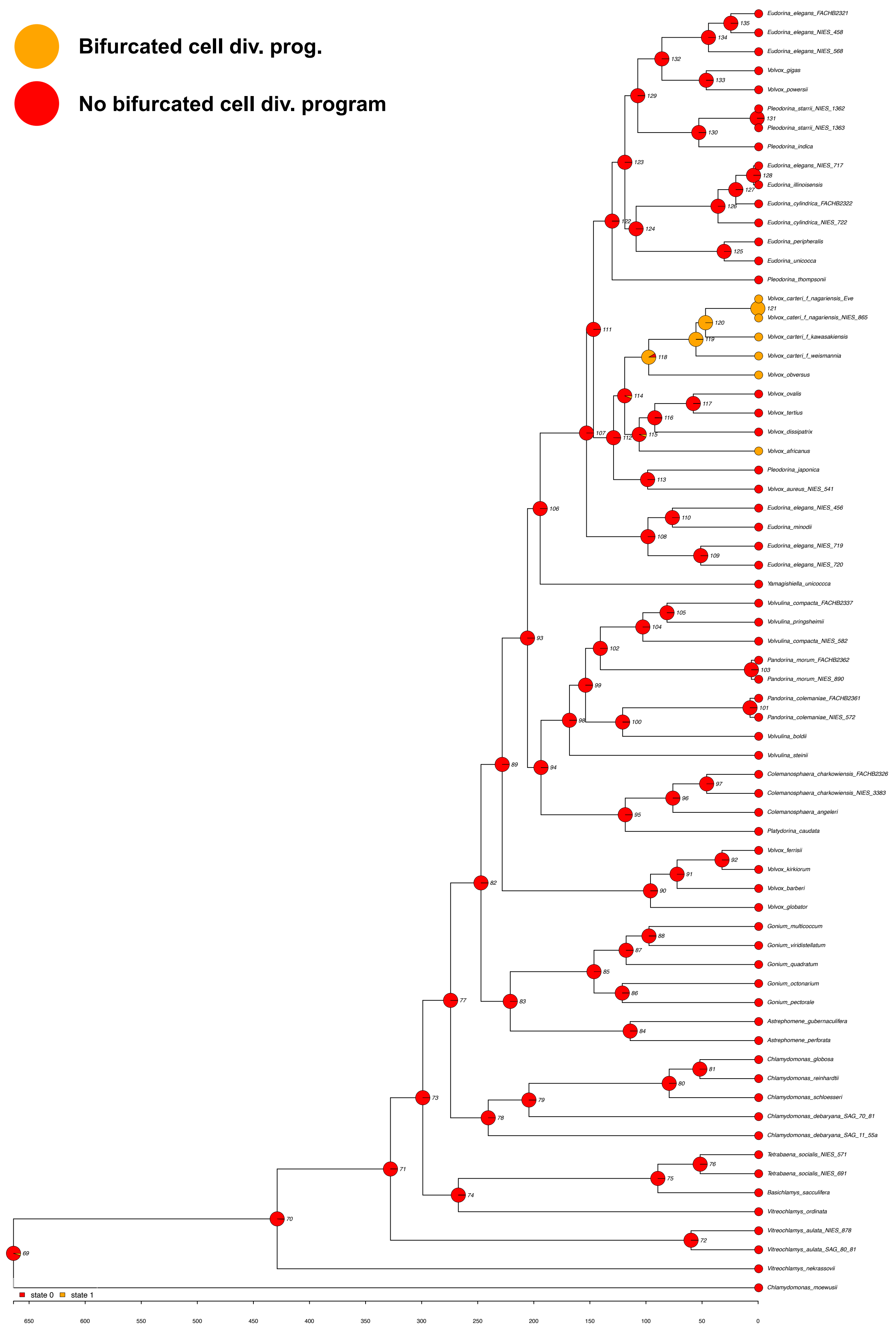

### cell_walls_to_ECM_tree.plot.pdf

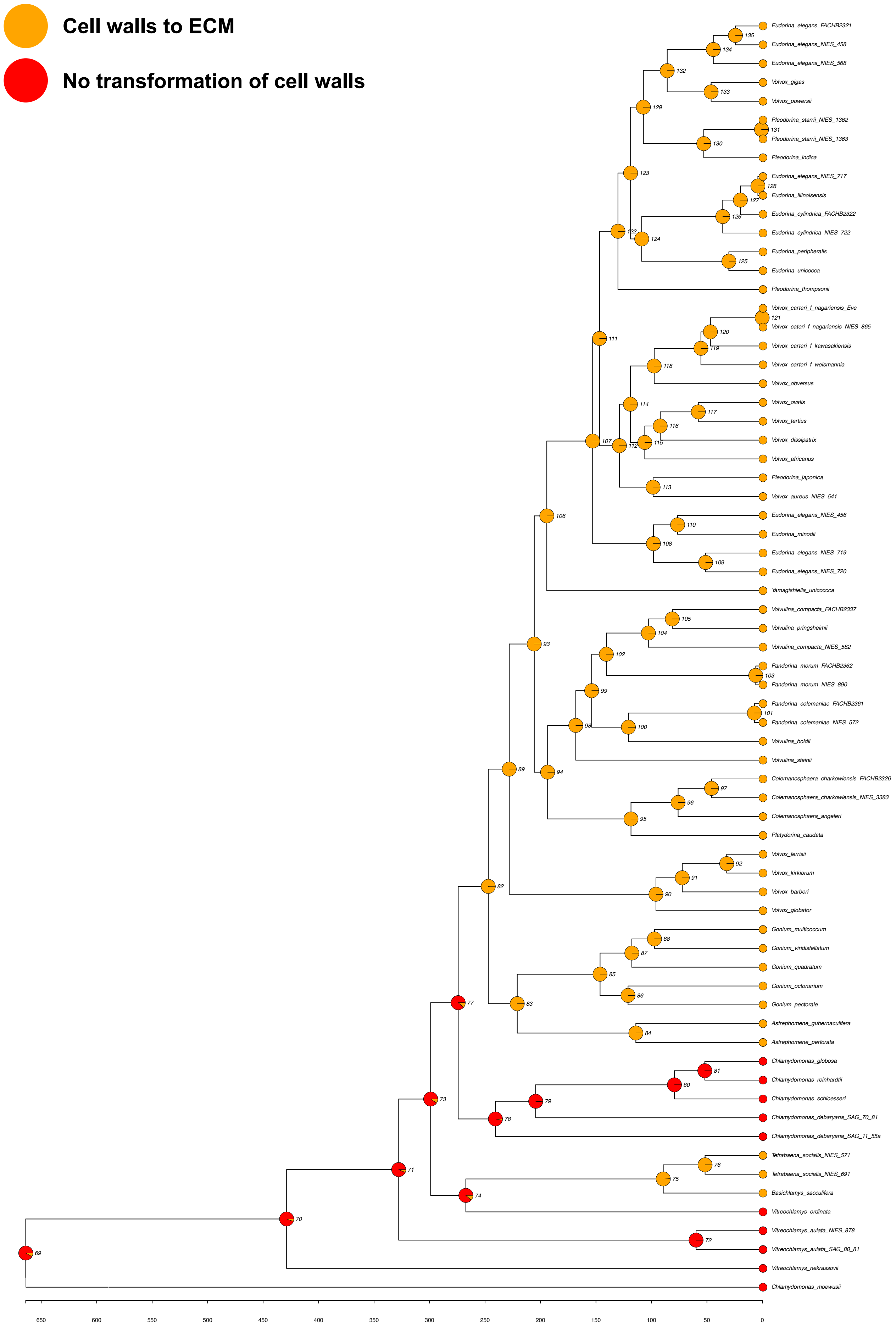

### cellularity_tree.plot.pdf

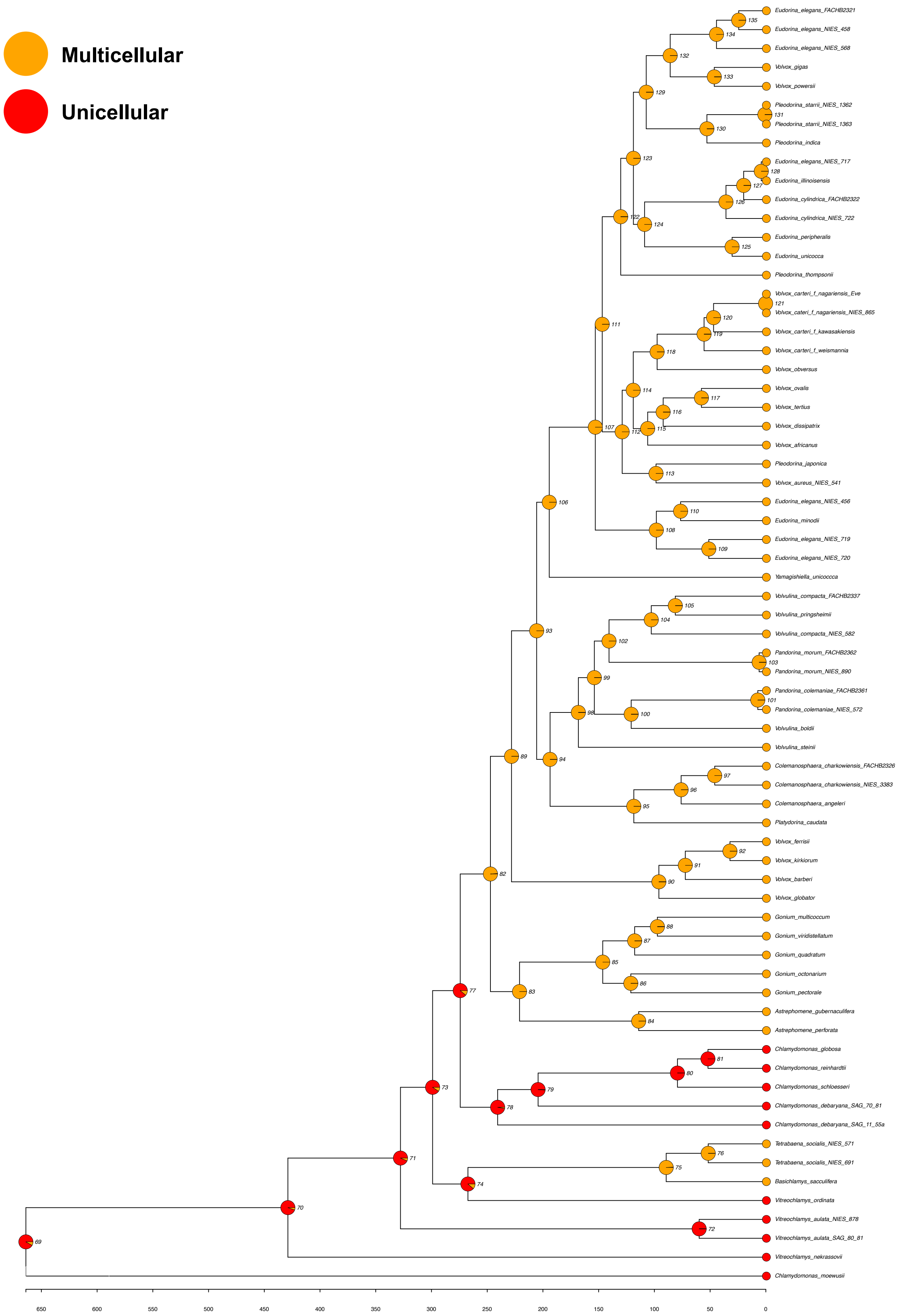

### cytokinesis_tree.plot.pdf

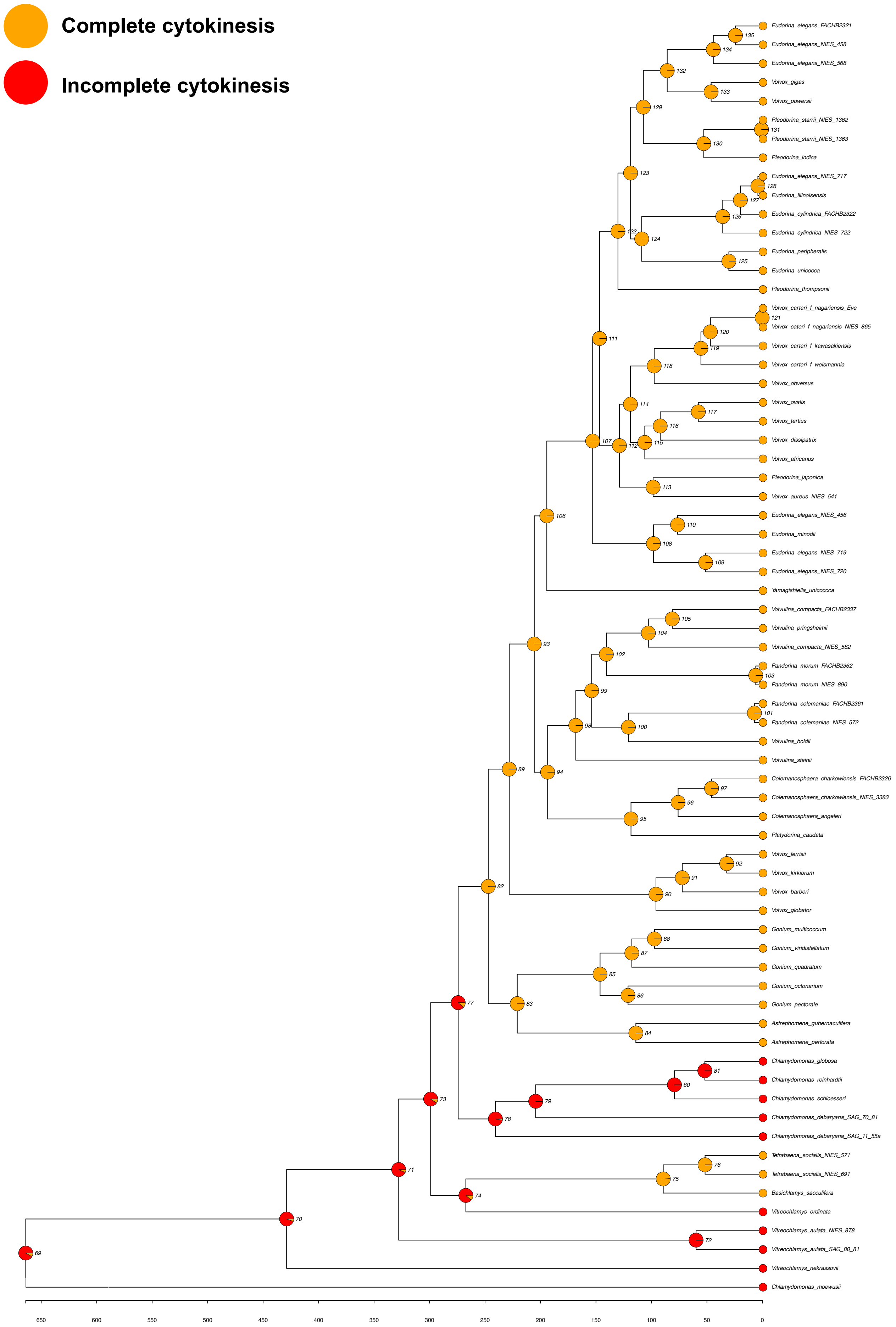

### differentiation3states_COMPLETE_tree.plot.pdf

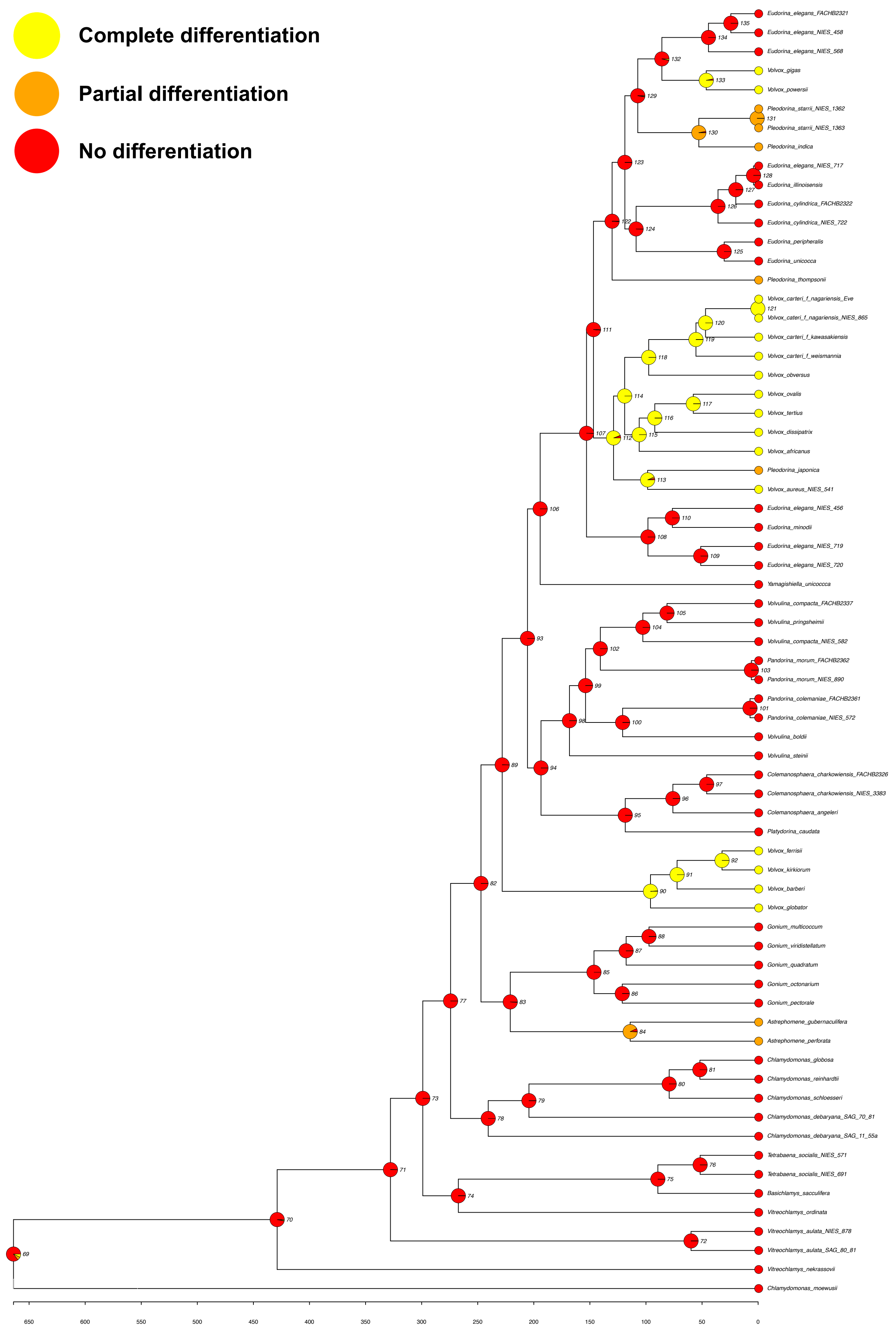

### dwarfmales_tree.plot.pdf

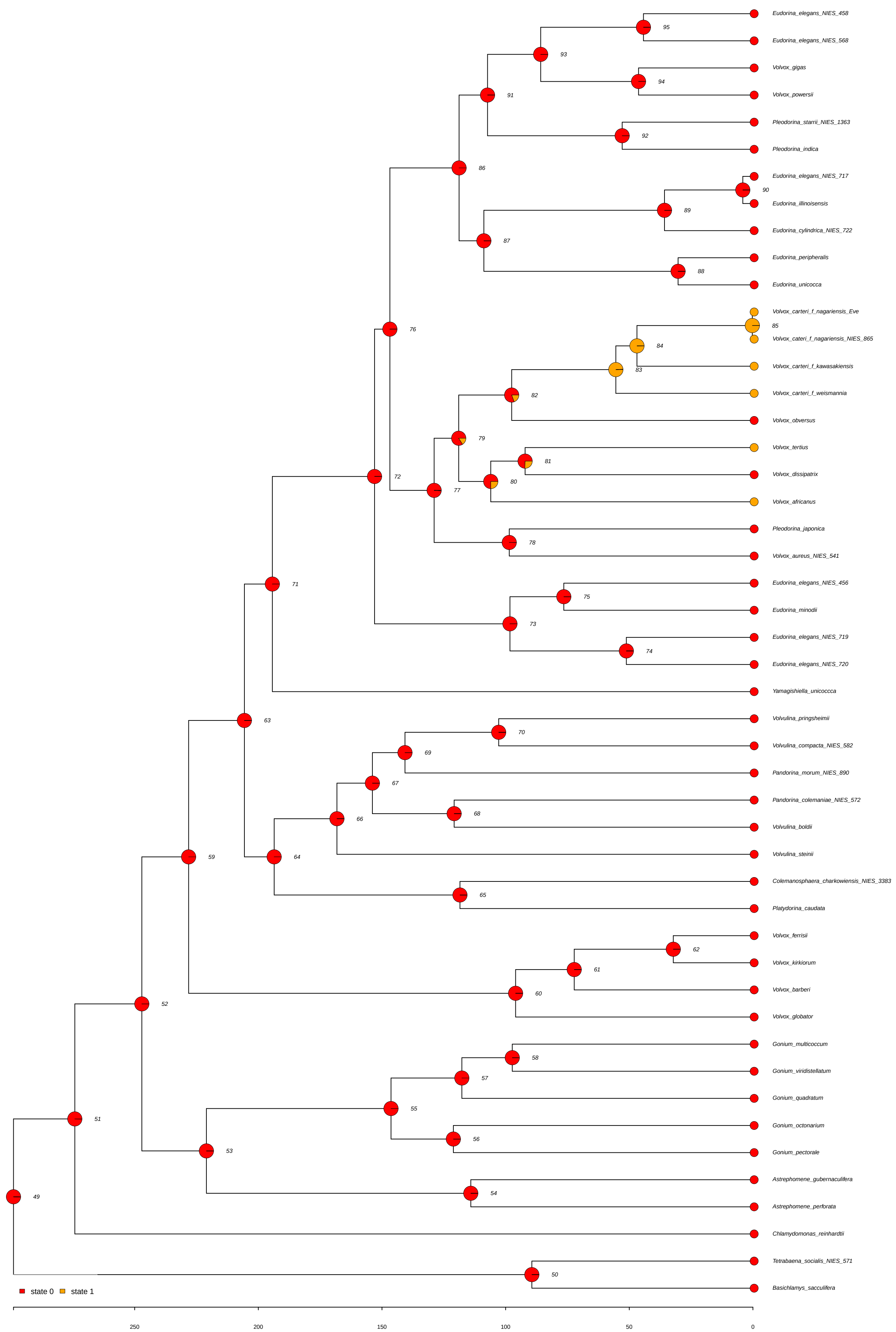

### extrafertile_females_tree.plot.pdf

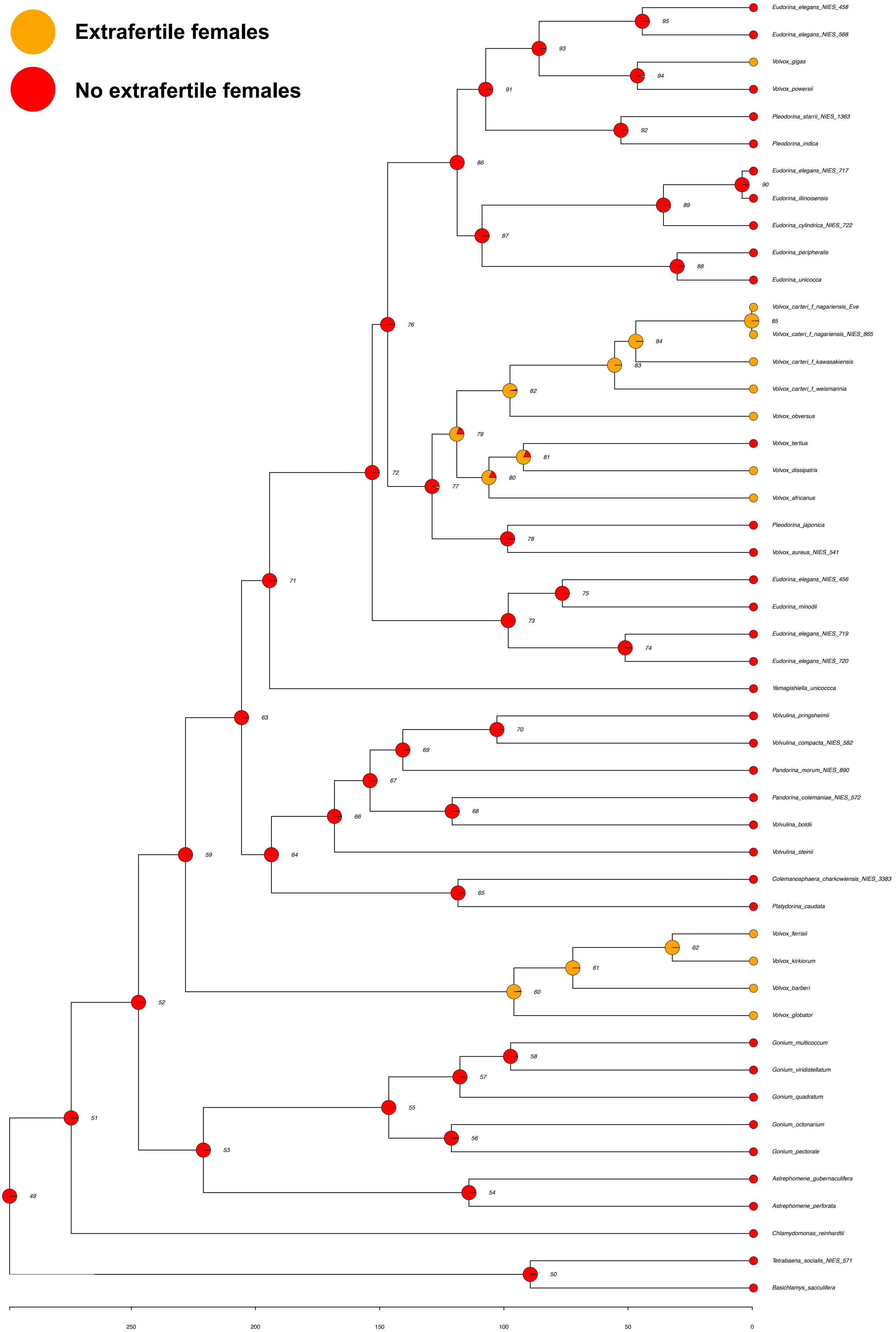

### fertilization_tree.plot.pdf

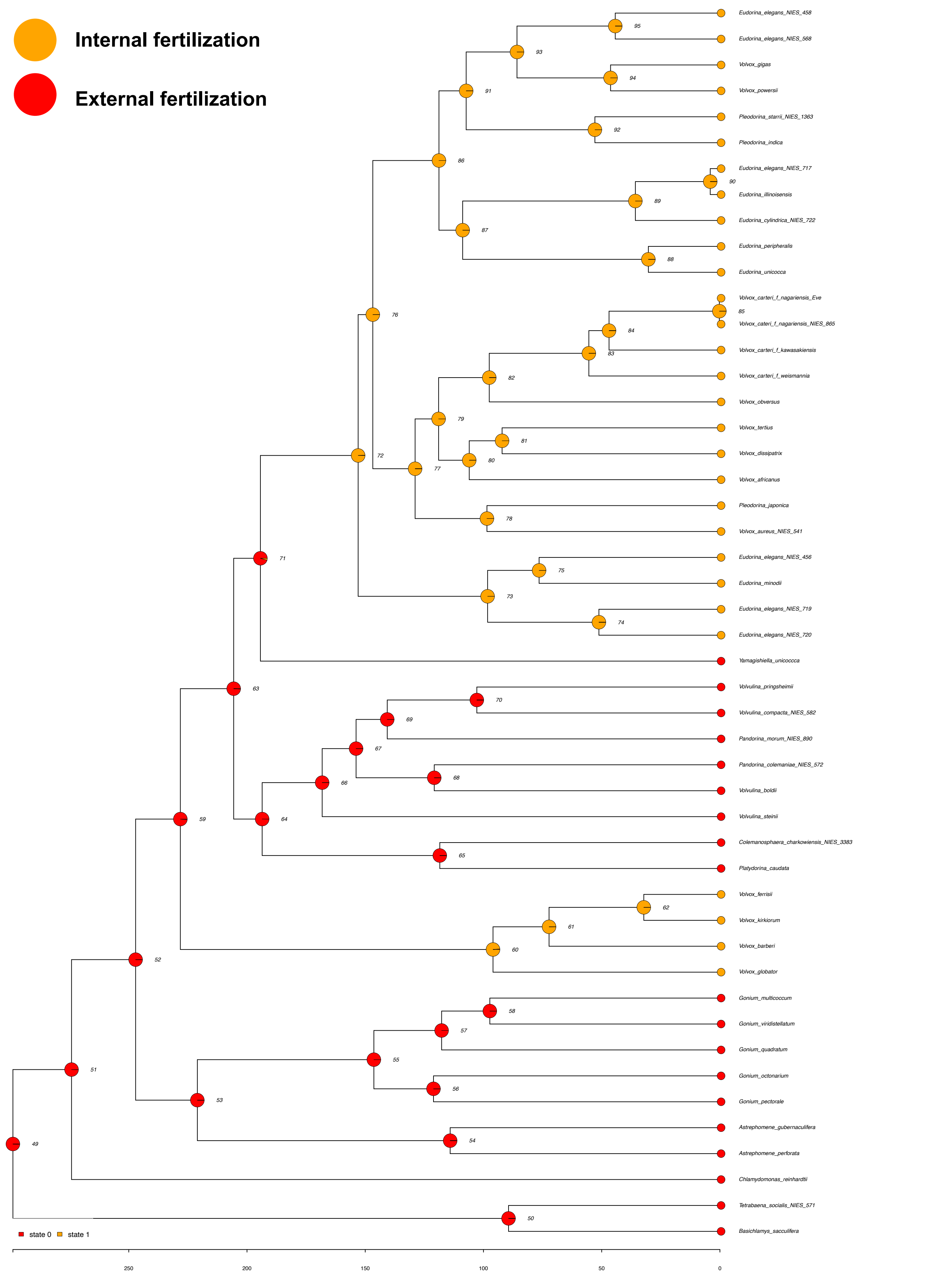

### gametic_diff_3states_tree.plot.pdf

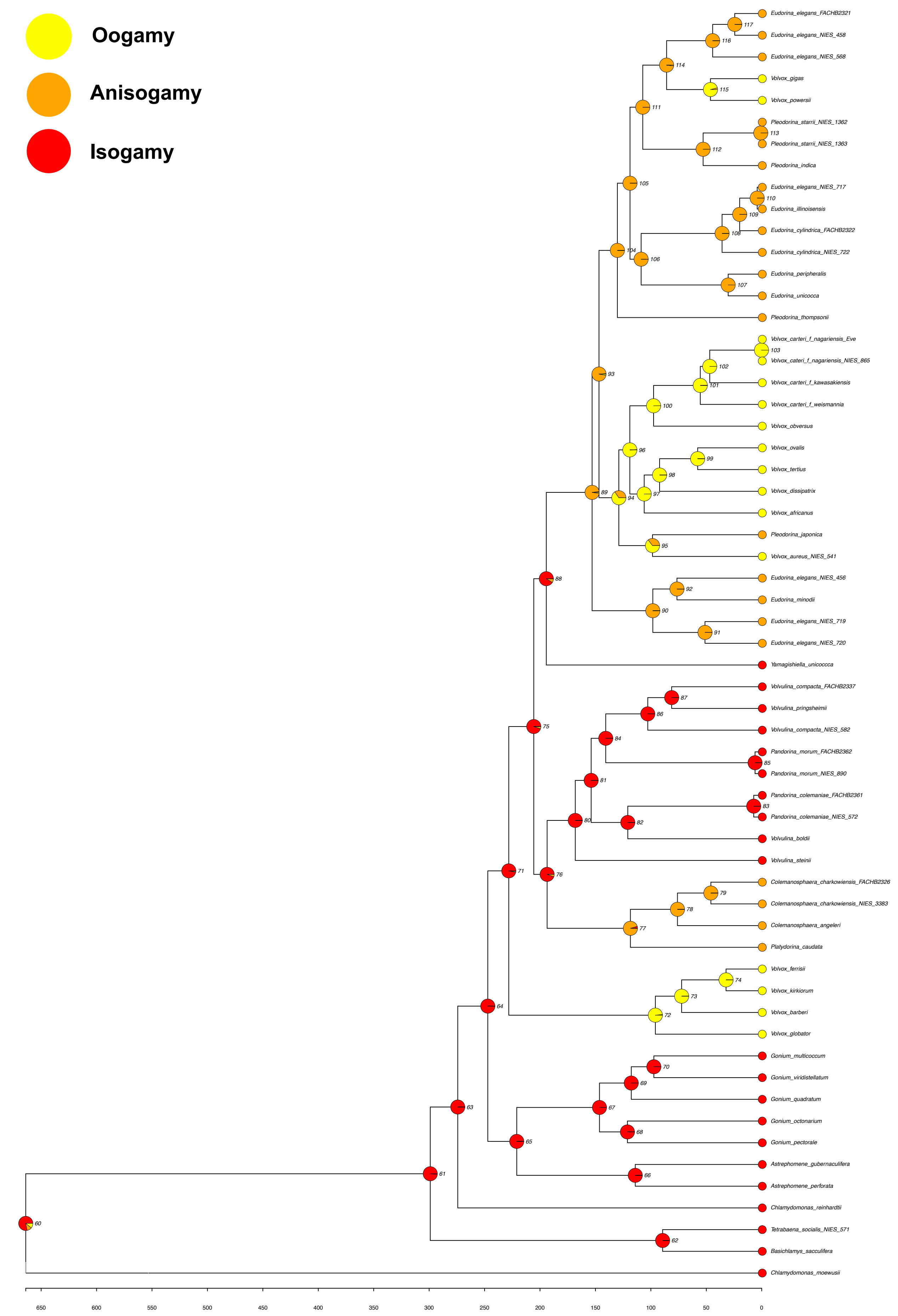

### gametic_differentiation_2states_tree.plot.pdf

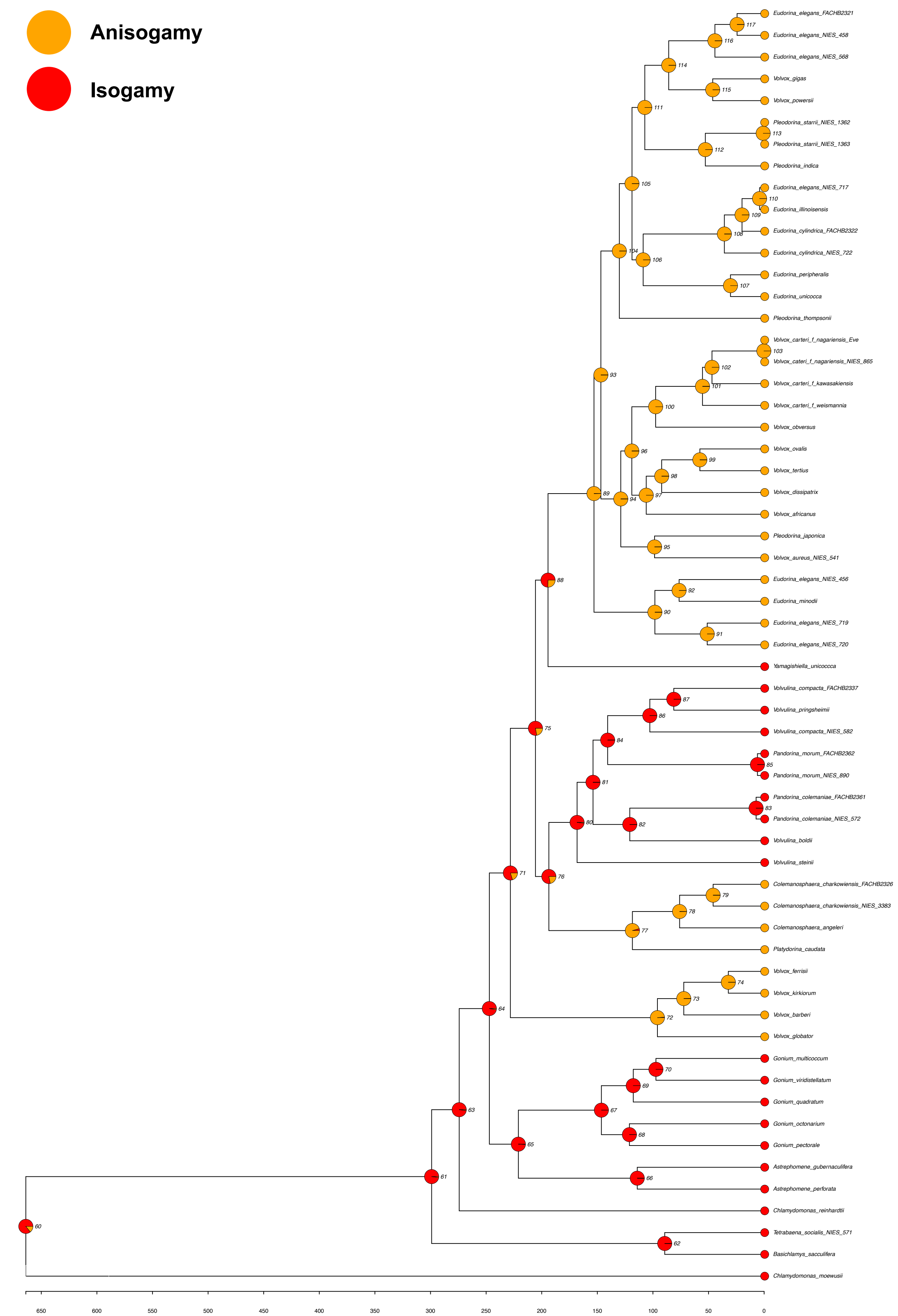

### gen_mod_cell_cum_tree.plot.pdf

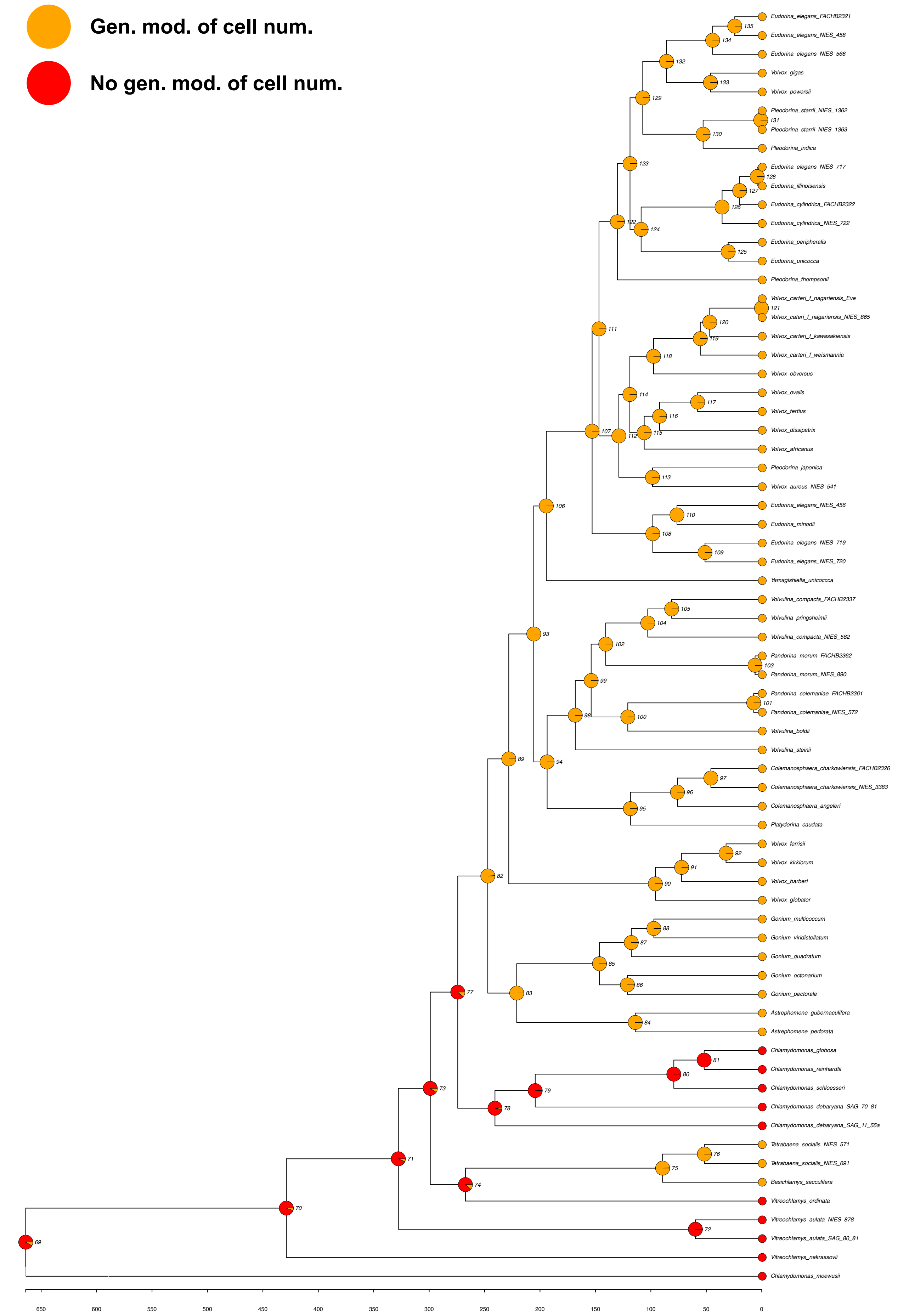

### inversion_2states_tree.plot.pdf

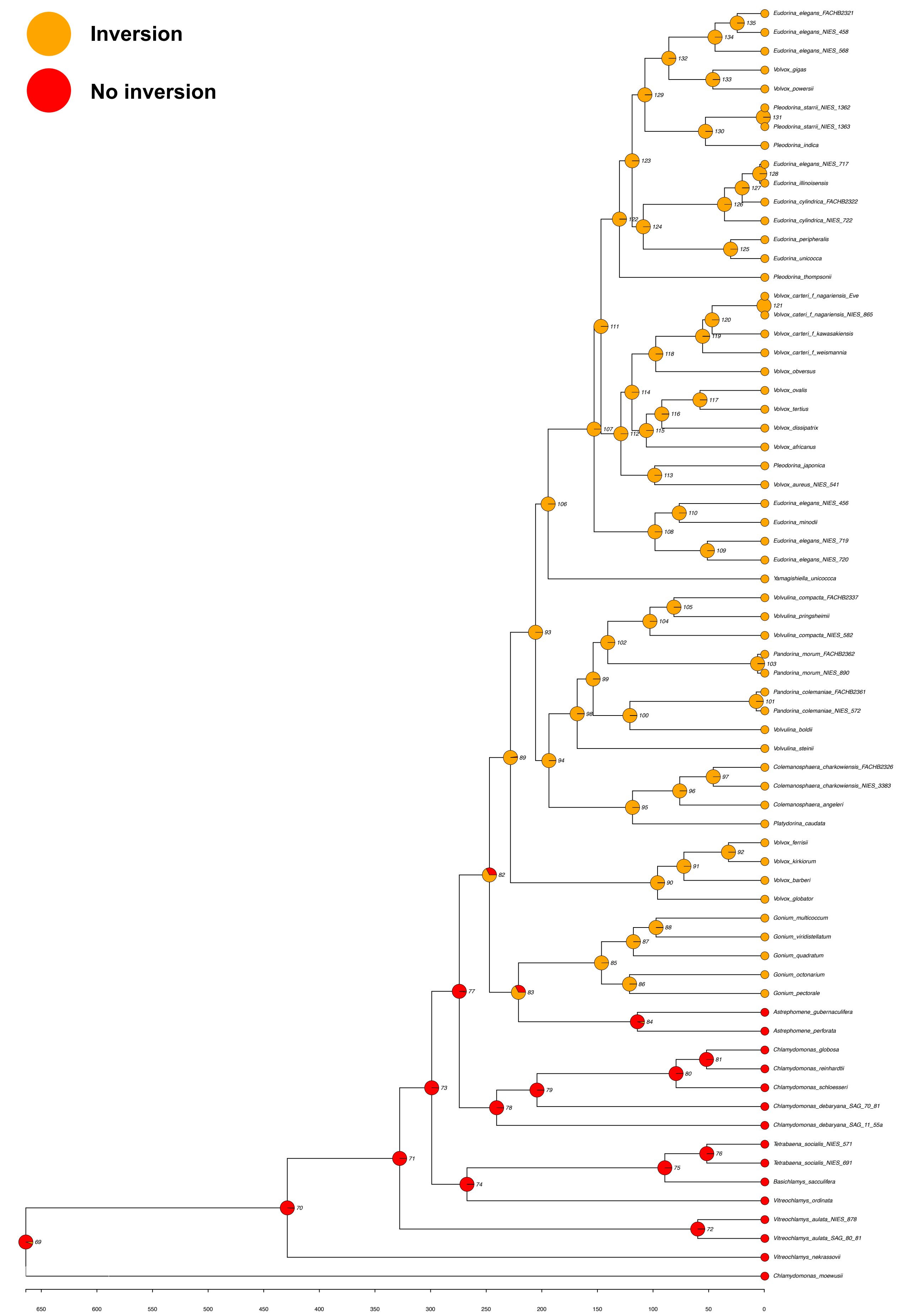

### inversion_3states_ordered_tree.plot.pdf

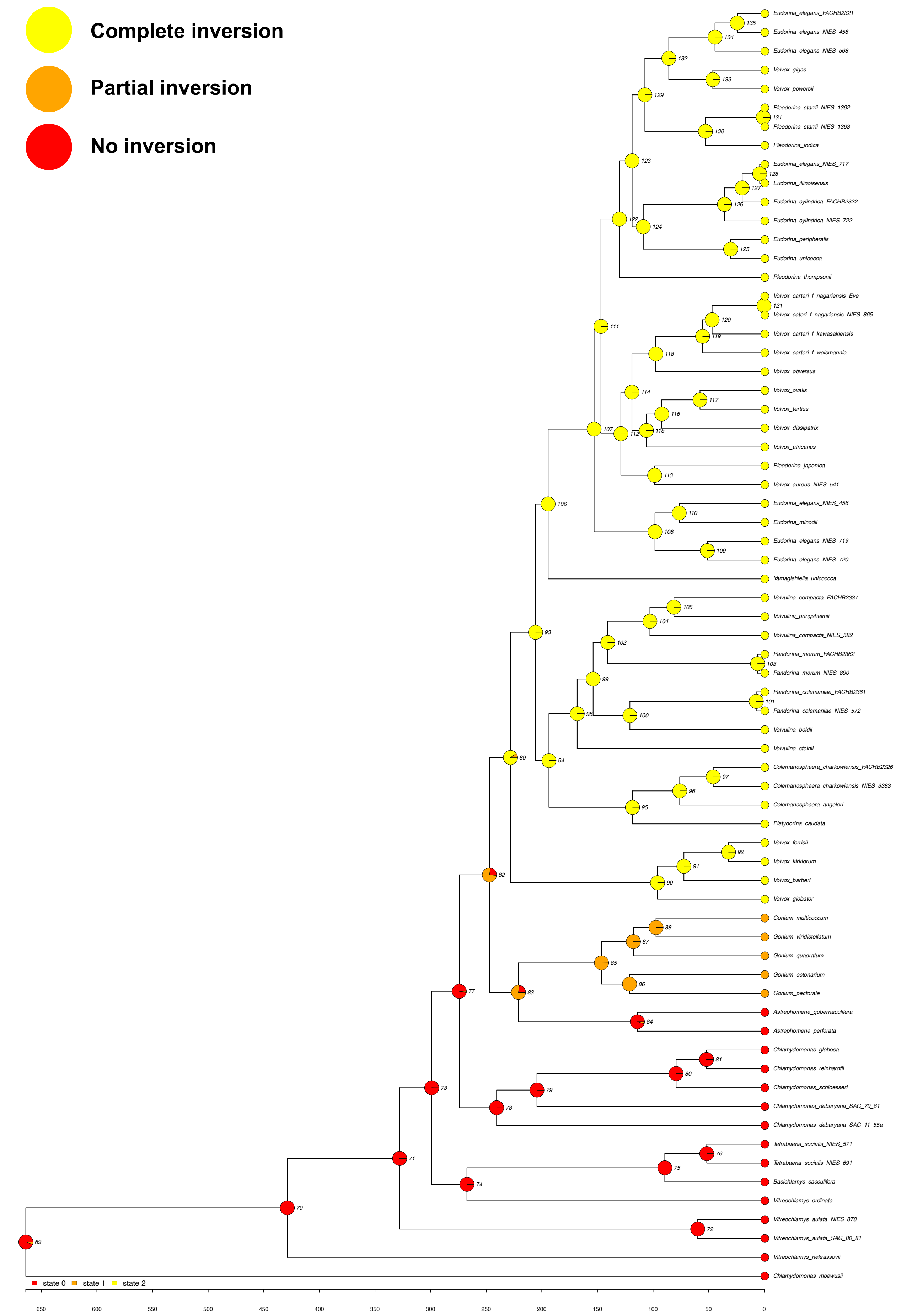

### inversion_3states_tree.plot.pdf

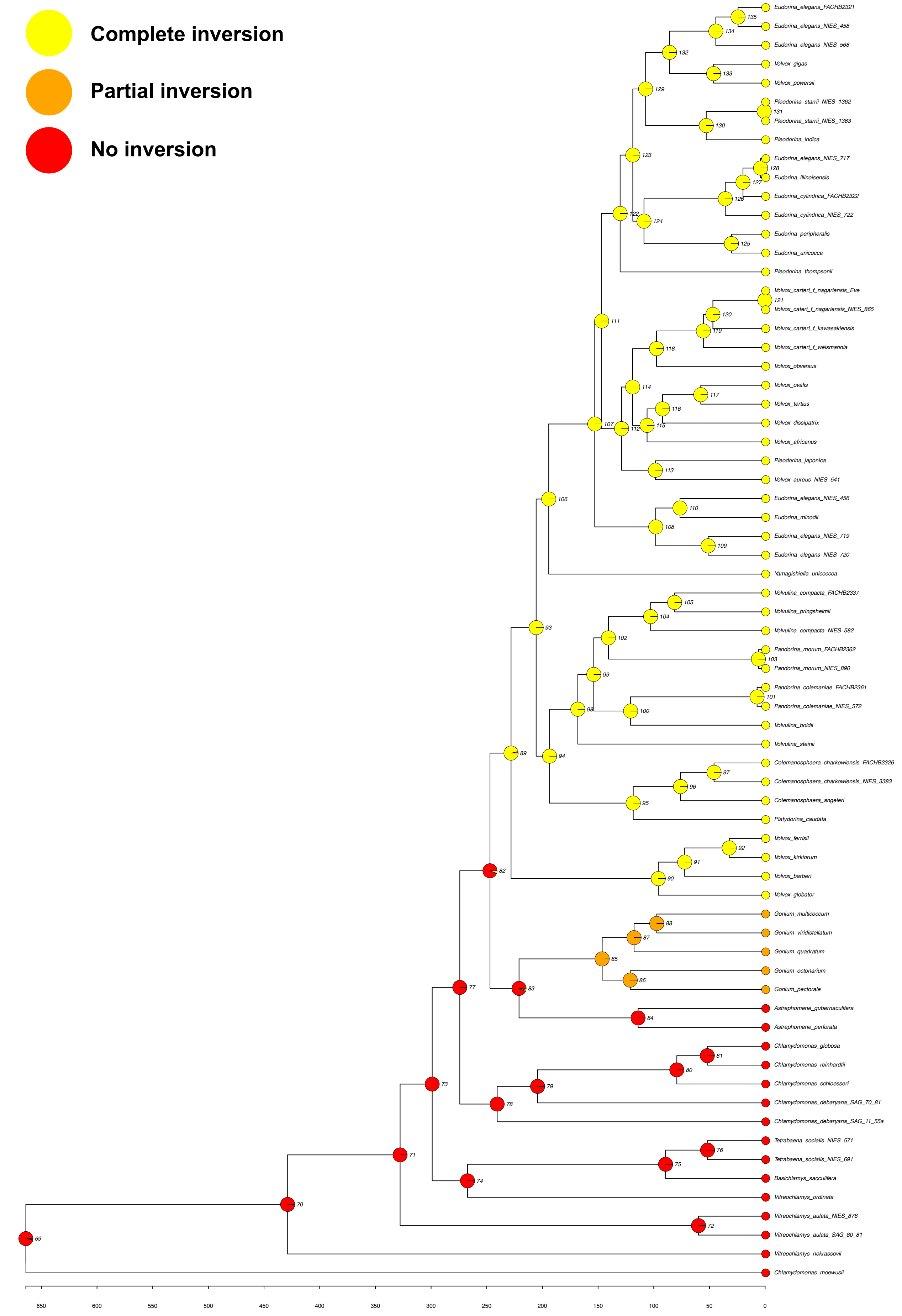

### meiotic_product_tree.plot.pdf

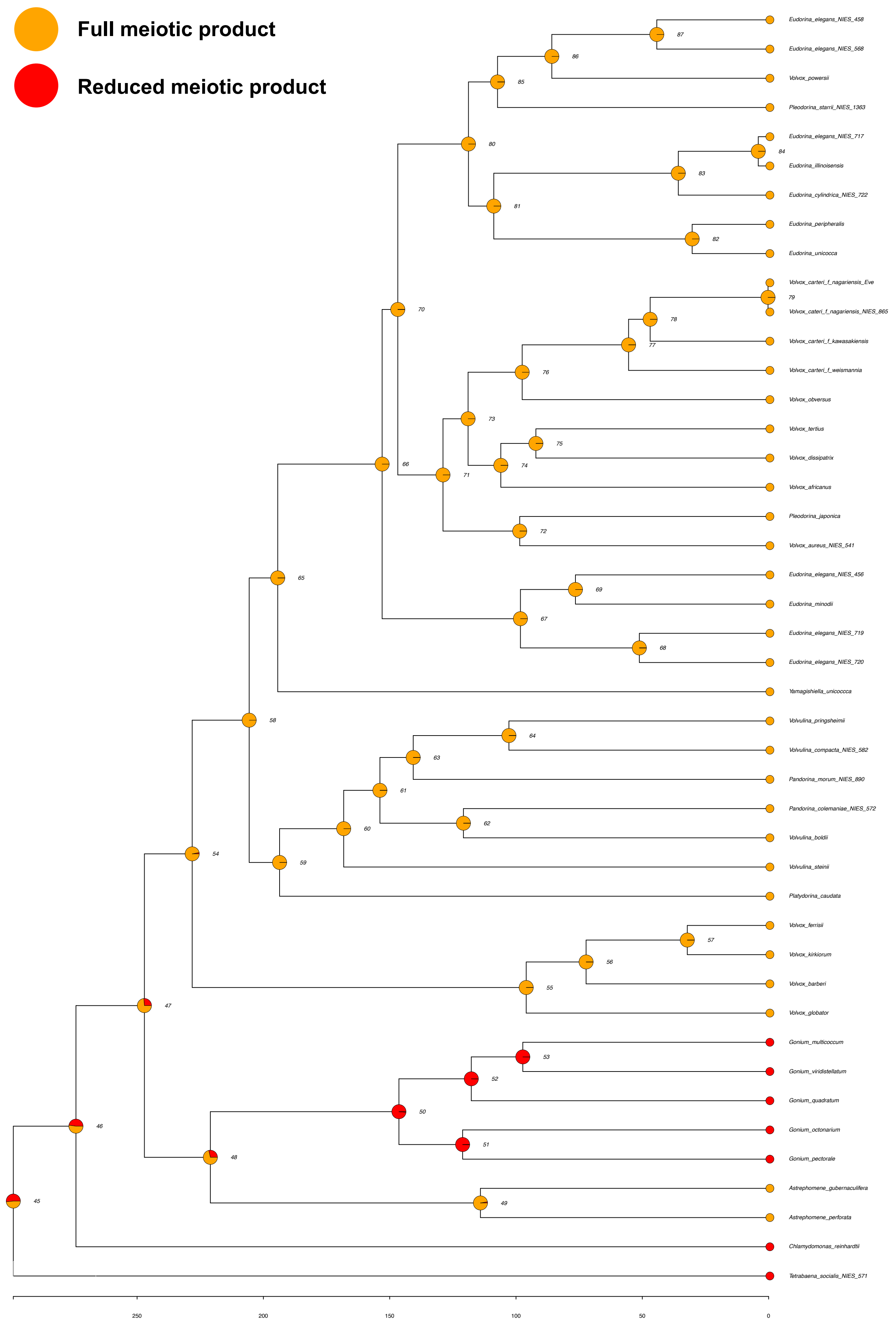

### mon_vs_di_tree.plot.pdf

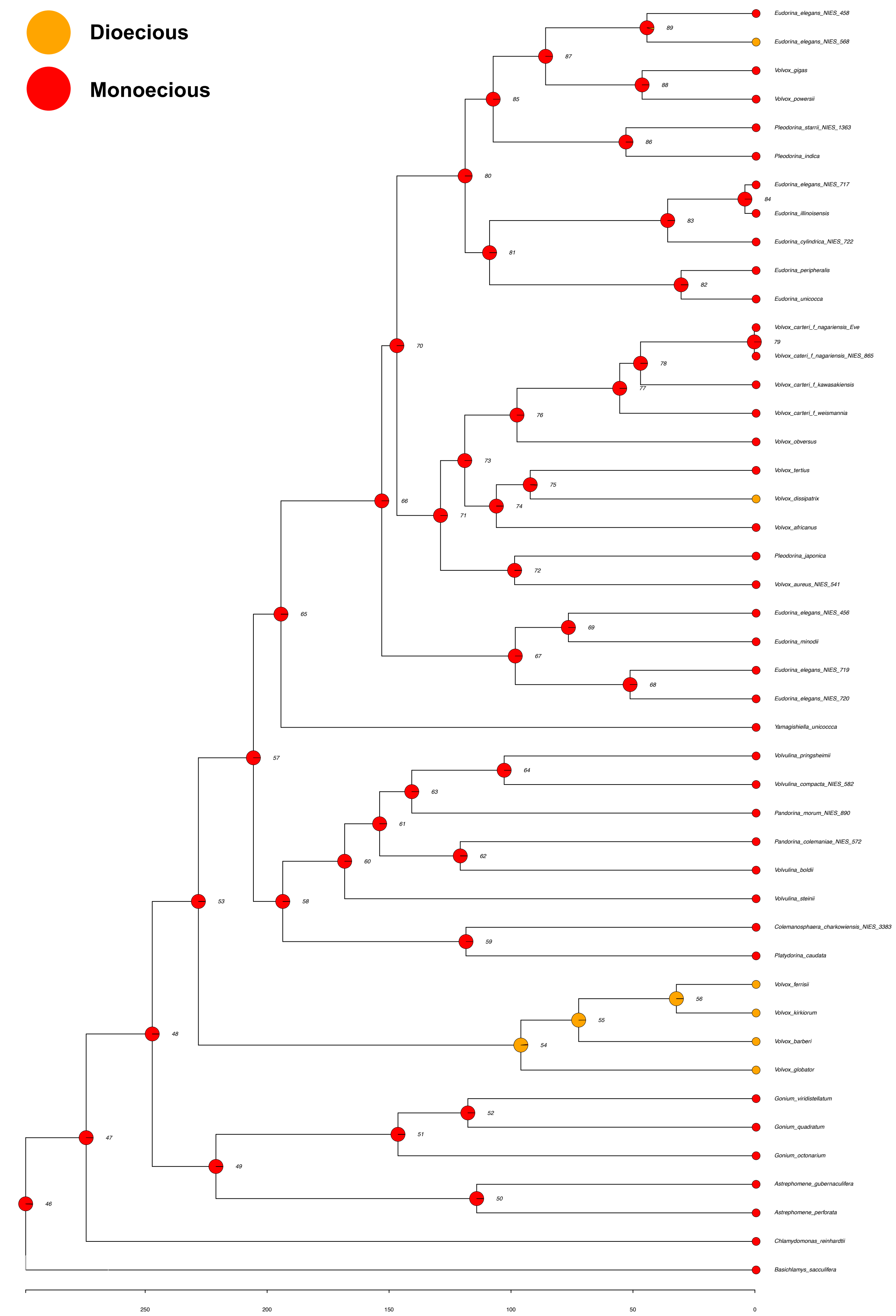

### polarity_tree.plot.pdf

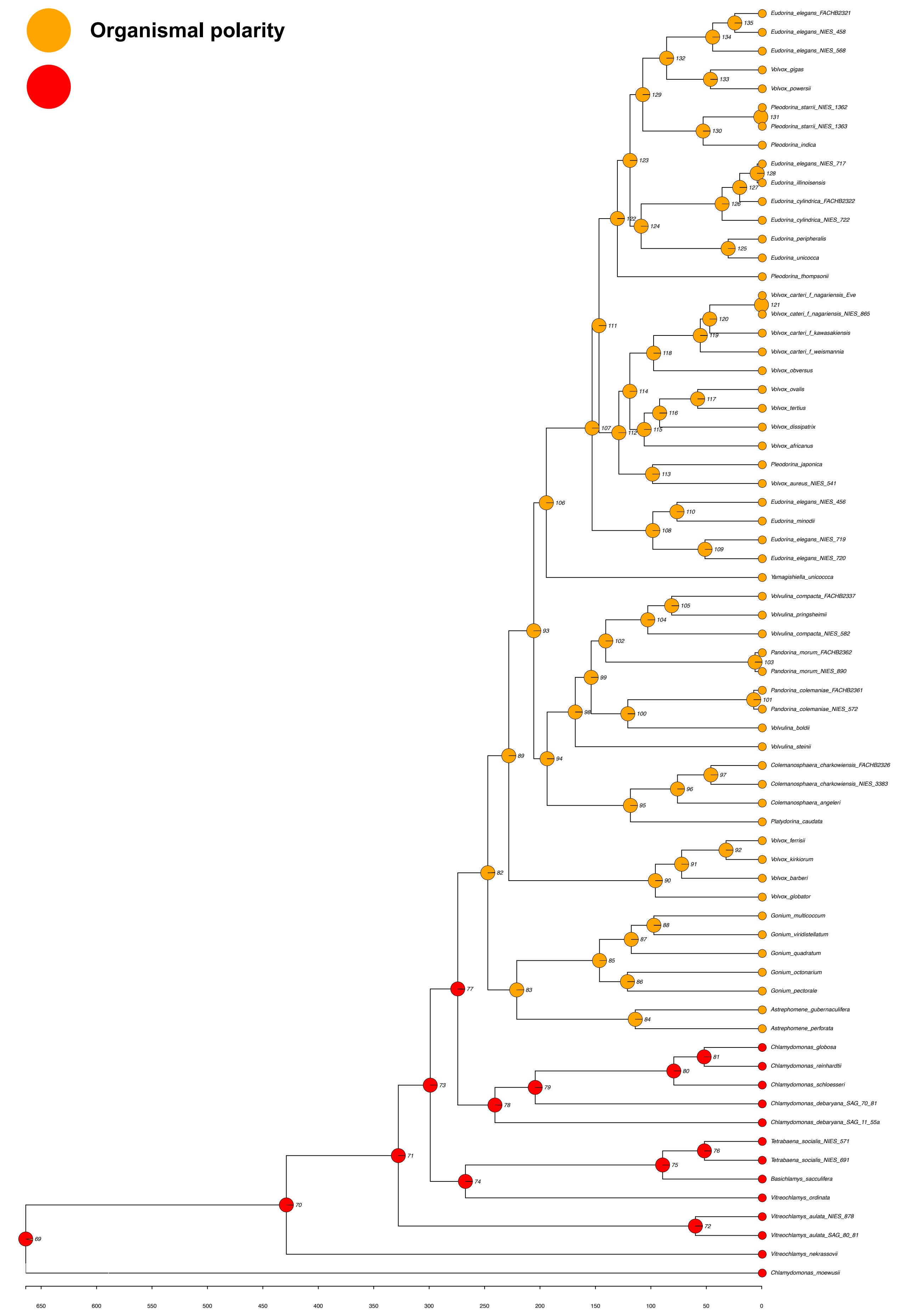

### selfing_tree.plot.pdf

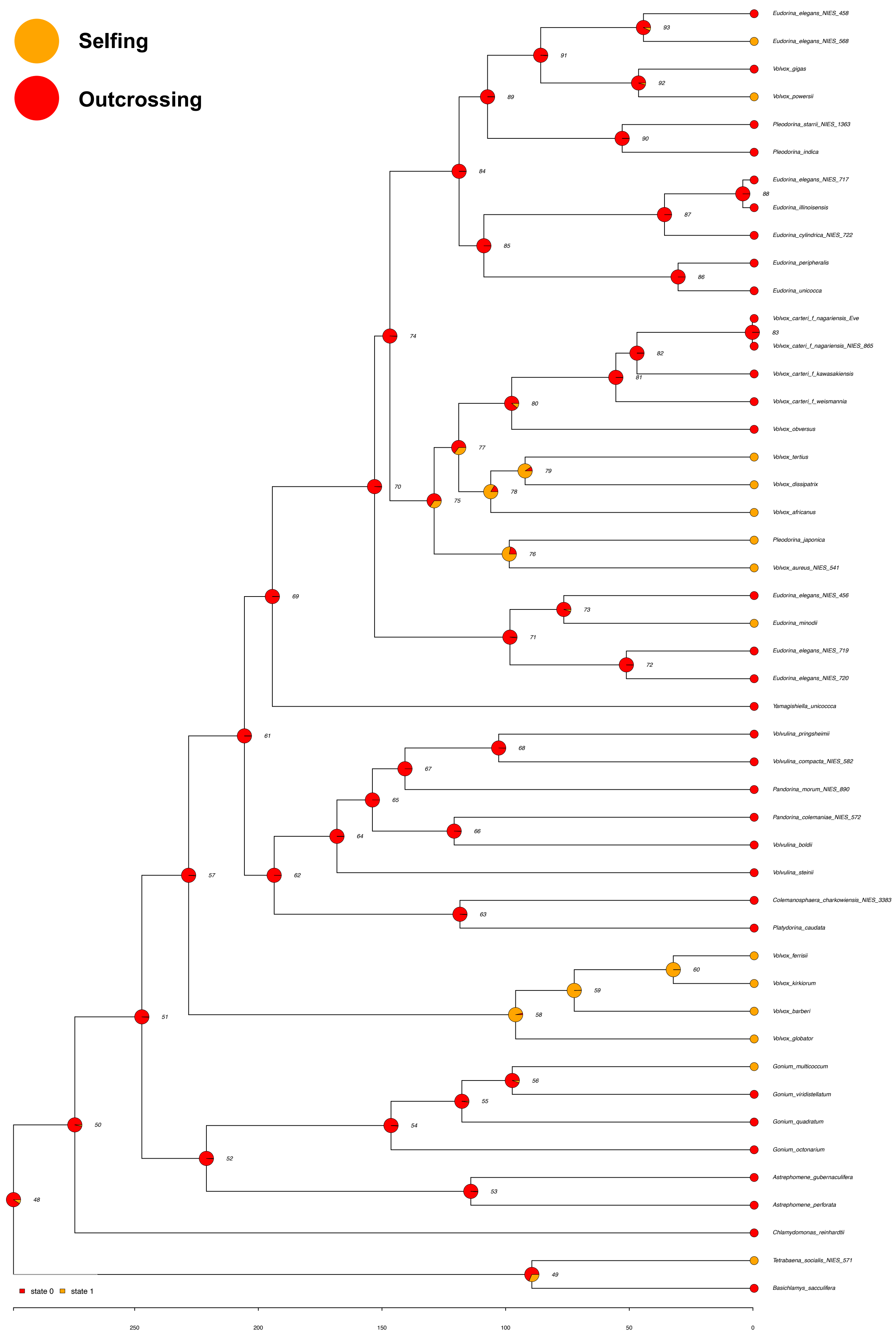

### spheroidal_body_plan_tree.plot.pdf

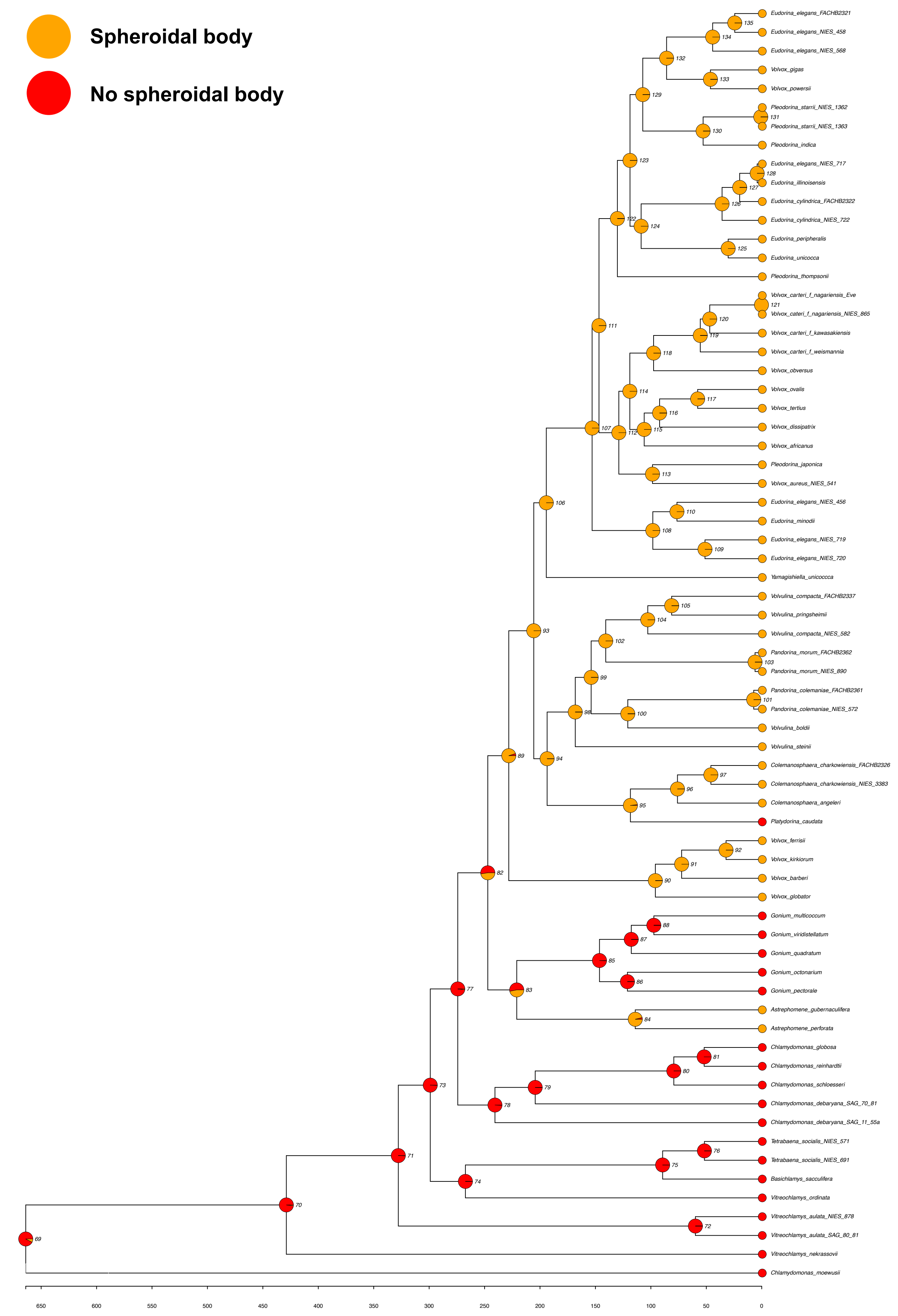
