## Supplementary methods for "Fossil-calibrated inference of divergence times among the Volvocine algae enables reconstruction of the steps that led to differentiated multicellularity"

***Fossil selections and node calibrations make possible molecular-clock analyses and the construction of an evolutionary time tree.*** Multiple sources of information can be used to calibrate a phylogenetic tree when performing a molecular dating analysis. These forms of information fall into three distinct categories:[1] verifiable terrestrial geological events,[2] secondary calibrations (those inferred in a separate study under another molecular dataset and calibration point),[3] and/or fossil taxa. While no calibration source is without its limitations, well-preserved fossils with distinguishing features that are either assignable to or that establish a relationship with extant taxa are highly favored. The fossilized remains of a taxon usually represent when a particular organism rose to abundance rather than when it emerged, thereby providing a minimum age constraint on the node to which it has been assigned.[4]

Since no reliable fossils exist for the volvocine algae, we selected 14 fossil taxa across the Archaeplastida where reliable fossils are abundant (**Figure 2** and **Table 1**). For each primary fossil calibration used in this study, each calibration point was constrained to a range rather than a fixed-point estimate, acknowledging the inherent uncertainty in fossil ages. With each date range, we specified a soft bound where 2.5% of the total probability mass is positioned outside of the specified lower and/or upper bounds.[5]

The root age of our tree, the divergence between red and green algae, was maximally constrained to 2000 MYA. This maximum constraint was informed by geological evidence indicating a rise in atmospheric oxygen ~2400 MYA;[6] we reason that it is unlikely that a common ancestor of red and green algae existed prior to elevated oxygen levels in the atmosphere and at the ocean surface. Since this date is most likely removed from the true root age of the Archaeplastida by hundreds of millions of years, this maximum is fitting for an upper constraint.

Two fossil calibrations were assigned within the Rhodophyta (red algae). The oldest divergence within the rhodophyte lineage was constrained to 1060-1030 MYA[7] based on *Bangiomorpha pubscens,*[8] the oldest red algal fossil reliably identified as such. Florideophyte divergence (620-600 MYA) was established by the age of unambiguous florideophytes in the Doushantuo Formation, China.[9]

Within Streptophyta, the clade composed of charophyte green algae and land plants, 8 nodes were calibrated using the fossil record. The oldest divergence within the Zygnemataceae was constrained to a minimum age of 350 MYA based on the oldest fossils assigned to this clade[10]. The dawn of land plants was informed by newly discovered fossil spores from the Ordovician[11] (MIN:480 MYA). *Cooksonia* with tracheids[12] (423-419 MYA) aided in estimating the emergence of vascular plants (i.e. tracheophytes), while the oldest known seeds[13] (MIN: 385 MYA) were used to calibrate the fern-seed plant split. The gymnosperm-angiosperm bifurcation was constrained to 330-323 MYA, based on the oldest *Cordaite,* a sister to coifiers[14], whereas the age of the *Amborella* and Nymplaeles divergence from other angiosperms was informed by Barremian flowers and pollen[15] (129-125 MYA). Finally, the split between chloroanthaceae/magnoliid and monocots + eudicots was calibrated using the oldest known chloranthaceous fossils[16] (MIN: 125 MYA), while the monocot-eudicot split was constrained to a minimum age of 113 MYA based on *Liliacidites* monocot pollen.[17]

Among the Chlorophyta, a major Archaeplastida clade consisting of the unicellular and multicellular green algae, 8 nodes were calibrated. The oldest prasinophyte divergence within the Micromonadophyceae was constrained to 600-580 MYA based on the oldest prasinophyte phycomata with punctate walls.[18] Within Trebouxiophyceae, the oldest *Botryococcus* divergence was informed by the earliest known fossils assigned to that genus (358-356 MYA).[10] *Proterocladus*, a cladophoracean fossil, was used to constrain the divergence between the Ulvophyceae and Bryopsidales + Chlorophyceae to 1056-948 MYA.[19] Within the Ulvophyceae, the earliest Ulotrichale divergence was constrained to 470-458 MYA based on *Vermiporella.*[20]

Three fossil taxa were discarded from our analysis because they yielded either highly variable estimates when fossil cross validation tests were performed. These validation tests involve a "leave-one-out" approach where a node's calibration is removed and the resulting inferred mean date and 95% HPD interval is compared to the fossil date. The oldest split within the Caulerpaceae was intended to be informed by *Margerita dorus*[21] (MIN: 505), the deepest divergence in the Caulerpaceae was estimated to have a mean date of ~200 MYA when this fossil taxa was removed (**Supplemental Table 2**). ). This fossil's phylogenetic placement is reliable, but nodes ages inferred using our *Caulerpa* molecular data do not accord with the age of the *Margerita dorus* fossil. An earlier inferred date compared to the fossil age suggests that this fossil was formed during a later period; therefore, we also deemed this fossil to be uninformative. The *Oedogonium* chlorophyte clade was originally going to be constrained to 393-382 MYA, based on *Paleoodogonium*;[22] however, fossil validation tests estimated ~58 MYA as the deepest divergence for the genus *Oedogonium* (**Supplemental Table 2**). Based on their morphology[22,23] and similarities in degradation[23] to extant *Oedogonium*, *Paleoodogoinum* fossils appear to be a reliable ancestor of *Oedogonium*. Our inferred validation dates indicate that *Paleoodogoinum* may be even more distantly related to *Oedogonium* than previously thought, thus we removed this fossil from consideration as a calibration point. The earliest known divergence within the Chaetophorales was intended to be informed by *Electrophycus* *astroplethus* (110-97 MYA),[24] however, the oldest divergence within the Chaetophorales was estimated to be much older at ~613 MYA (**Supplemental Table 2**).

***Key Archaeplastida divergence times are largely congruent with previous studies.***

Beginning at the root, red and green algae diverged from their last common ancestor in the Early to Middle Mesoproterozoic (**Figure 2**), most likely ~1385 MYA (**Figure 2**), consistent with a Middle Mesoproterozoic estimate by Lang et al.[25] The red-green algal divergence has also been inferred to have occurred as early as the Late Paleoproterozoic,[26–28] in the Early Mesoproterozoic,[29–31] or as late at the Early Neoproterozoic.[28] Our 95% HPD intervals for the red green algal divergence (1634-1187 MYA) complement these earlier reports. Branching from the root, we find that two crown groups, Rhodophyta and Chlorophyta, emerge in the Late Mesoproterozoic, and Streptophyta, a crown group containing land plants, originated in the Early Neoproterozoic (**Figure 2**), as reported in Sánchez-Baracaldo et al.[26] When considering the 95% HPD interval for each of the crown groups, our estimates encompass those of other studies. For example, Rhodophyta has a 95% HPD interval between 1331 (upper bound postdates the first fossil appearance of red algae by ~100 MY) and 926 MYA (**Figure 3B**), and this overlaps with estimates from a study exclusively sampling taxa within the Archaeplastida[32] as well as another that samples across all 3 domains of life.[29] 95% HPD Confidence intervals for Chlorophyta and Streptophyta are between 1308 and 1102 MYA and 1038 and 858 MYA, respectively, and comfortably overlap with those from previous reports.[31,33,34]

Key divergences within two crown groups overlap with inferred age estimates elsewhere. The Florideophyceae, the largest and most diverse clade of rhodophytes, were estimated to have arisen sometime between 724 to 564 MYA (**Table 1**). Yang et al.,[32] who performed dense taxonomic sampling of Rhodophyta, inferred that the Florideophyceae emerged in the Neoproterozoic, 879-681 MYA. Among the Streptophytes, land plants appear to originate either in the Late Cambrian or Late Ordovician, according to a 95% HPD interval spanning 485-446 MYA (**Table 1**). These estimates support claims by Strother and Foster[11] who hypothesized that embryophytes may have emerged earlier than the Ordovician following their discovery of a land plant fossil dating to the Early Ordovician. Additionally, our inferred 95% HPD estimates overlap those for Embryophyta reported by Morris et al.[34] Within Chlorophyta, the divergence between the Ulvophyceae I clade and Ulvophyceae II + other chlorophytes is estimated to have occurred sometime between 1042 and 945 MYA. The reason our 95% HPD intervals encompass the Early Neoproterozoic, and are outside those reported in other studies indicating a Mid Neoproterozoic divergence,[35,36] is because we imposed a fossil age constraint on *Proterocladus* of 1056-948 MYA.

20. Kozlowski, R., and Kazmierczak, J. ON TWO ORDOVICIAN CALCAREOUS ALGAE. 45.

21. Morris, S.C., and Robison, R.A. MORE SOFT-BODIED ANIMALS AND ALGAE FROM THE MIDDLE CAMBRIAN OF UTAH AND BRITISH COLUMBIA'. 48.

22. Baschnagel, R.A. (1966). New Fossil Algae from the Middle Devonian of New York. Transactions of the American Microscopical Society *85*, 297. 10.2307/3224643.

23. Graham, L.E., Cook, M.E., Wilcox, L.W., Graham, J., Taylor, W., Wellman, C.H., and Lewis, L. (2013). Resistance of Filamentous Chlorophycean, Ulvophycean, and Xanthophycean Algae to Acetolysis: Testing Proterozoic and Paleozoic Microfossil Attributions. International Journal of Plant Sciences *174*, 947–957. 10.1086/670591.

24. Poinar, G., and Brown, A.E. (2021). A green algae (Chaetophorales: Chaetophoraceae) in Burmese amber. Historical Biology *33*, 323–327. 10.1080/08912963.2019.1616719.

25. Lang, D., Weiche, B., Timmerhaus, G., Richardt, S., Riaño-Pachón, D.M., Corrêa, L.G.G., Reski, R., Mueller-Roeber, B., and Rensing, S.A. (2010). Genome-Wide Phylogenetic Comparative Analysis of Plant Transcriptional Regulation: A Timeline of Loss, Gain, Expansion, and Correlation with Complexity. Genome Biology and Evolution *2*, 488–503. 10.1093/gbe/evq032.

26. Sánchez-Baracaldo, P., Raven, J.A., Pisani, D., and Knoll, A.H. (2017). Early photosynthetic eukaryotes inhabited low-salinity habitats. Proc. Natl. Acad. Sci. U.S.A. *114*. 10.1073/pnas.1620089114.

27. Strassert, J.F.H., Irisarri, I., Williams, T.A., and Burki, F. (2021). A molecular timescale for eukaryote evolution with implications for the origin of red algal-derived plastids. Nat Commun *12*, 1879. 10.1038/s41467-021-22044-z.

28. Eme, L., Sharpe, S.C., Brown, M.W., and Roger, A.J. (2014). On the Age of Eukaryotes: Evaluating Evidence from Fossils and Molecular Clocks. Cold Spring Harbor Perspectives in Biology *6*, a016139–a016139. 10.1101/cshperspect.a016139.

29. Betts, H.C., Puttick, M.N., Clark, J.W., Williams, T.A., Donoghue, P.C.J., and Pisani, D. (2018). Integrated genomic and fossil evidence illuminates life's early evolution and eukaryote origin. Nat Ecol Evol *2*, 1556–1562. 10.1038/s41559-018-0644-x.

30. Lutzoni, F., Nowak, M.D., Alfaro, M.E., Reeb, V., Miadlikowska, J., Krug, M., Arnold, A.E., Lewis, L.A., Swofford, D.L., Hibbett, D., et al. (2018). Contemporaneous radiations of fungi and plants linked to symbiosis. Nat Commun *9*, 5451. 10.1038/s41467-018-07849-9.

31. Parfrey, L.W., Lahr, D.J.G., Knoll, A.H., and Katz, L.A. (2011). Estimating the timing of early eukaryotic diversification with multigene molecular clocks. Proc. Natl. Acad. Sci. U.S.A. *108*, 13624–13629. 10.1073/pnas.1110633108.

32. Yang, E.C., Boo, S.M., Bhattacharya, D., Saunders, G.W., Knoll, A.H., Fredericq, S., Graf, L., and Yoon, H.S. (2016). Divergence time estimates and the evolution of major lineages in the florideophyte red algae. Sci Rep *6*, 21361. 10.1038/srep21361.

33. Becker, B. (2013). Snow ball earth and the split of Streptophyta and Chlorophyta. Trends in Plant Science *18*, 180–183. 10.1016/j.tplants.2012.09.010.

34. Morris, J.L., Puttick, M.N., Clark, J.W., Edwards, D., Kenrick, P., Pressel, S., Wellman, C.H., Yang, Z., Schneider, H., and Donoghue, P.C.J. (2018). The timescale of early land plant evolution. Proc. Natl. Acad. Sci. U.S.A. *115*. 10.1073/pnas.1719588115.

35. Herron, M.D., Hackett, J.D., Aylward, F.O., and Michod, R.E. (2009). Triassic origin and early radiation of multicellular volvocine algae. Proceedings of the National Academy of Sciences *106*, 3254–3258. 10.1073/pnas.0811205106.

36. Del Cortona, A., Jackson, C.J., Bucchini, F., Van Bel, M., D’hondt, S., Škaloud, P., Delwiche, C.F., Knoll, A.H., Raven, J.A., Verbruggen, H., et al. (2020). Neoproterozoic origin and multiple transitions to macroscopic growth in green seaweeds. Proc. Natl. Acad. Sci. U.S.A. *117*, 2551–2559. 10.1073/pnas.1910060117.
